# Two structurally different oomycete lipophilic MAMPs induce distinctive plant immune responses

**DOI:** 10.1101/2021.10.22.465218

**Authors:** Mohammad Shahjahan Monjil, Hiroaki Kato, Satomi Ota, Kentaro Matsuda, Natsumi Suzuki, Shiho Tenhiro, Ayane Tatsumi, Sreynich Pring, Atsushi Miura, Maurizio Camagna, Takamasa Suzuki, Aiko Tanaka, Ryohei Terauchi, Ikuo Sato, Sotaro Chiba, Kazuhito Kawakita, Makoto Ojika, Daigo Takemoto

## Abstract

Plants recognize a variety of external signals and induce appropriate mechanisms to increase their tolerance to biotic and abiotic stresses. Precise recognition of attacking pathogens and induction of effective resistance mechanisms are critical functions for plant survival. Some molecular patterns unique to a certain group of microbes, microbe-associated molecular patterns (MAMPs), are sensed by plant cells as non-self molecules via pattern recognition receptors. While MAMPs of bacterial and fungal origin have been identified, reports on oomycete MAMPs are relatively limited. This study aimed to identify MAMPs from an oomycete pathogen *Phytophthora infestans*, the causal agent of potato late blight. Using reactive oxygen species (ROS) production and phytoalexin production in potato as markers, two structurally different groups of elicitors, namely ceramides and diacylglycerols were identified. *P. infestans* ceramides (Pi-Cer A, B and D) induced ROS production, while diacylglycerol (Pi-DAG A and B), containing eicosapentaenoic acid (EPA) as a substructure, induced phytoalexins production in potato. The molecular patterns in Pi-Cers and Pi-DAGs essential for defense induction were identified as 9-methyl-4,8-sphingadienine (9Me-Spd) and 5,8,11,14-tetraene-type fatty acid (5,8,11,14-TEFA), respectively. These structures are not found in plants, but in oomycetes and fungi, indicating that they are microbe molecular patterns recognized by plants. When Arabidopsis was treated with Pi-Cer D and EPA, partially overlapping but different sets of genes were induced. Furthermore, expression of some genes is upregulated only after the simultaneous treatment with Pi-Cer D and EPA, indicating that plants combine the signals from simultaneously recognized MAMPs to adapt the defense response to pathogens.

## Introduction

During the course of their evolution, plants were always faced with the challenges of environmental microorganisms but have survived by developing physical barriers or inducible resistance strategies against pathogens. The first step of inducible plant defense against potential pathogens is the recognition of molecular patterns of microbes, referred to as MAMPs or PAMPs (microbe- or pathogen-associated molecular patterns). Plant cells recognize molecular patterns unique to a certain group of microbes as non-self molecules via pattern recognition receptors (PRRs) (Ranf, 2017; Ngou et al., 2022). In Arabidopsis, for instance, chitin, a major component of the fungal cell wall, is recognized by a PRR complex containing LysM-RLK and co-receptor CERK1, while a conserved sequence of bacterial flagellin (flg22) and an elongation factor EF-Tu (elf18) induce plant defense via PRRs, FLS2 and EFR, respectively, in conjuncture with co-receptor BAK1 (Ranf, 2017). While a variety of MAMPs of bacterial and fungal origin, as well as the mechanisms for their recognition in plants have been studied, reports of MAMPs from oomycete pathogens are relatively limited.

Oomycete plant pathogens resemble fungal pathogens in various aspects. Besides morphological similarities, such as filamentous hyphae, they are also able to form appressoria and haustoria during infection in host plants. These similarities are however merely the result of convergent evolution, since oomycetes are phylogenetically closely related to diatoms and brown algae (Stramenopiles), and only distantly related to fungi (Opisthokonta). Therefore, the natures of oomycetes and true fungi are rather different. For example, the somatic thallus of oomycetes is diploid, while it is haploid (or dikaryotic) for true fungi, and oomycetes have mitochondria with tubular cristae whereas true fungi have flattened ones. More importantly in terms of interaction with plants, the composition of the cell wall and cell membrane of oomycete and fungi is distinctive. Although varying from species to species, the fungal cell wall mainly contains β-1,3-glucan and chitin (β-1,4-N-acetylglucosamine) (Free, 2013), whereas the oomycete cell wall consists mainly of cellulose (β-1,4-glucan) and β-1,3-glucan (Mélida et al., 2013). The plasma membrane of oomycetes contains fucosterol as the end sterol, but ergosterol is the major end sterol for fungi (Gaulin et al., 2010). Consequently, plants may have distinct mechanisms to recognize oomycete and fungal pathogens to activate appropriate defense reactions.

Of the various oomycete genera, *Phytophthora*, *Pythium* and *Peronospora* /*Hyaloperonospora* (downy mildew) are particularly noteworthy, as they include many species that are problematic plant pathogens (Kamoun et al., 2015). These plant pathogenic oomycete genera are predicted to have evolved from a common phytopathogenic ancestor species (Thines and Kamoun, 2010). Undoubtedly the most infamous representative of pathogenic oomycetes is *Phytophthora infestans*, the causal agent of potato late blight, which is responsible for the Great Famine of the 1840s in Ireland. This pathogen is an ongoing problem even today, and the total costs for control efforts and yield losses by *P. infestans* are estimated in the range of multi-billion dollars annually (Garelik, 2002; Fry, 2008). Problems caused by *P. infestans* are particularly serious in developing countries where fungicides are used as the primary solution for disease management. Although potato cultivars with introduced resistance (*R*) gene(s) have been employed, due to selection pressure on these effectors in the pathogen population, *R*-gene dependent resistances have been lost in many varieties (Forbes, 2012).

Previously, Bostock et al. (1981) identified polyunsaturated fatty acids (PUFAs) such as eicosapentaenoic acid (EPA) and arachidonic acid (AA) from *P. infestans* as elicitor molecules. EPA can be considered as a MAMP of oomycete pathogens, as EPA is not found in higher plants, but is an essential component of the plasma membrane of oomycete cells (Bostock et al., 2011). In this study, we purified two lipophilic elicitors from *P. infestans*. Using induction of reactive oxygen species (ROS) production or phytoalexin production in potato as markers for the purification of elicitors, we identified two structurally different groups of elicitors, namely ceramides and diacylglycerols. *P. infestans* ceramide (Pi-Cer) elicitors induced ROS production, while diacylglycerol (Pi-DAG) elicitors induced the accumulation of phytoalexins in potato. RNAseq analysis was performed in Arabidopsis treated with Pi-Cer, EPA (a substructure of Pi-DAG), or the mixture of both elicitors, to investigate the difference in the activity of the two elicitors on the induction of defense genes, and the effect of simultaneous recognition of both MAMP elicitors. We have previously reported that a *P. infestans* ceramide Pi-Cer D is cleaved by an apoplastic ceramidase NCER2 and the resulting 9-methyl-branched sphingoid base is recognized by a lectin receptor-like kinase RDA2 in Arabidopsis (Kato et al., 2022). Cerebrosides (a group of glycosphingolipids) have been identified as elicitor molecules from rice blast pathogen *Pyricularia oryzae*, which also contain 9-methyl-branched sphingoid base (9Me-Spd) as part of their epitope structure (Koga et al., 1998, Umemura et al., 2000).

## RESULTS

### Purification of *Phytophthora infestans* elicitors which induce ROS production of potato suspension-cultured cells

Previously, we have reported that crude elicitor derived from the mycelia of *P. infestans* extracted by methanol (Pi-MEM) can induce the resistance reactions in potato (Monjil et al., 2015). Treatment with Pi-MEM can induce the ROS production in potato suspension cells and leaves, and production of sesquiterpenoid phytoalexins (rishitin, lubimin and oxylubimin) in potato tubers (Fig. 1). To purify the elicitor molecules from the mycelia of *P. infestans* and determine their structure, Pi-MEM was first fractionated by its solubility in water and butanol. In this study, the lipophilic (butanol soluble) fraction was further separated by column chromatography, and the elicitor fractions were selected using their ROS inducing activity in cultured potato cells (Fig. S1). Only fractions with clear elicitor activity were used for further purification processes. Six fractions with significant ROS-producing activity were further purified (Fig. S1) and the chemical structures of these elicitors were analyzed by NMR and mass spectroscopy (Figs. S2, S3, S5-7 and S9). To compare the structural difference between elicitors and inactive related substances, two fractions with no significant elicitor activity were also structurally analyzed (Figs. S1, S4 and S8). Further structural analyses (Figures S10-S16, and Supplemental document) showed that these eight substances could be divided into two groups, ceramide (Cer) and ceramide-phosphoethanolamine (CerPE). We designated the four ceramide compounds Pi-Cer A-D (*<u>P</u>. infestans* <u>Cer</u>amide) and the remaining four compounds Pi-CerPE A-D respectively. Except for the phosphoethanolamine in Pi-CerPE, compounds that share the same alphabetical suffix are otherwise structurally identical (Fig. 2 and S17A). Pi-Cer A, B and D were able to induce ROS production, whereas Pi-Cer C had only a marginal ROS inducing activity (Figs. 2 and S1). Similarly, Pi-CerPE A, B and D were active elicitors for ROS production (Figs. S1 and S17). For both Pi-Cers and Pi-CerPEs, the D type ones were the most active, followed by B and A in that order (Figs. 2 and S17). An Arabidopsis transformant containing a *LUC* transgene under the control of *WRKY33* (AT2G38470) promoter (pWRKY33-LUC) (Kato et al., 2022) was employed for the detection of elicitor activity of Pi-Cers and Pi-CerPEs in Arabidopsis. Treatment of Pi-Cers and Pi-CerPEs can induce the transient activation of the promoter with a peak at approx. 90 min after elicitor treatment. Same as in the case of ROS production in potato suspension cells (Figs. 2 and S17), Pi-Cer D and Pi-CerPE D showed the highest elicitor activity among other Pi-Cers and Pi-CerPEs, and clearly lower elicitor activity was detected for Pi-Cer C and Pi-Cer PE C in Arabidopsis (Fig. S18, as shown by Kato et al. 2022).

**Figure 1.**
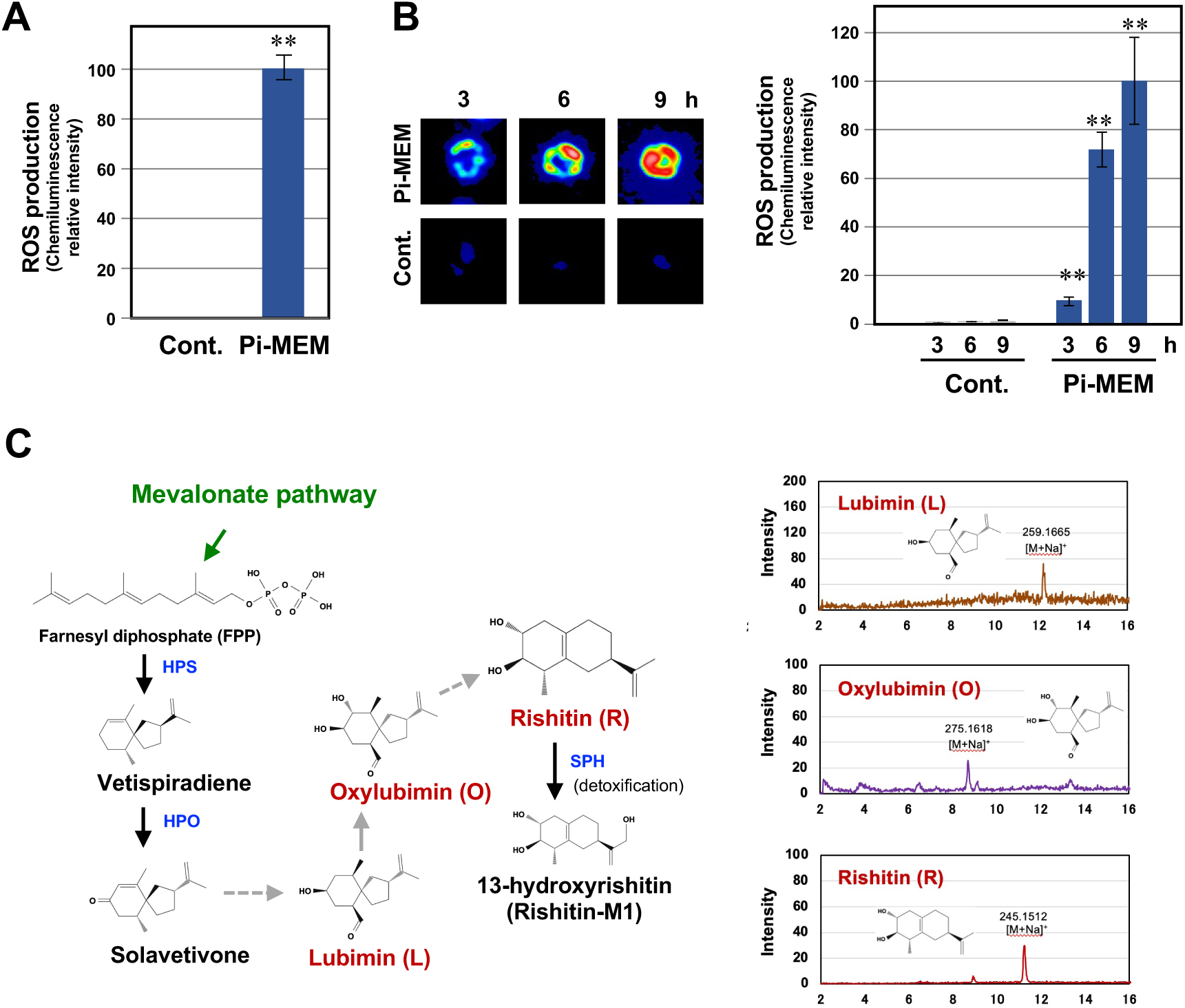
Elicitor activity of methanol extract of *Phytophthora infestans* mycelia (Pi-MEM) in potato. (A) Potato suspension cultured cells were treated with 0.3% DMSO (Cont.) or 30 µg/ml Pi-MEM and production of reactive oxygen species (ROS) was detected as L-012-mediated chemiluminescence 3 h after the treatment. Data are means ± SE (n = 3). Scores shown are chemiluminescence intensities relative to that of Pi-MEM-treated cells (B) Potato leaves were treated with 1 % DMSO (Cont.) or 100 µg/ml Pi-MEM and production of ROS was detected as L-012-mediated chemiluminescence. Data are means ± SE (n = 3). Scores shown are chemiluminescence intensities relative to that of Pi-MEM-treated leaves at 9 h. Data marked with asterisks are significantly different from control as assessed by two-tailed Student’s *t* tests: \*\**P* < 0.01. (C) Left, putative biosynthetic pathway of potato phytoalexins, lubimin, oxylubimin and rishitin. HPS, *Hyoscyamus muticus* premnaspirodiene synthase; HPO *H. muticus* premnaspirodiene oxygenase; SPH, sesquiterpenoid phytoalexins hydroxylase (Camagna et al., 2020). Potato tubers were treated with 3 % DMSO (Cont.) or 1 mg/ml Pi-MEM and phytoalexins were extracted 48 h after treatment. Produced phytoalexins were detected by thin-layer chromatography (middle) or LC/MS (right). Data shown are the representative results of at least 3 separate experiments.

**Figure 2.**
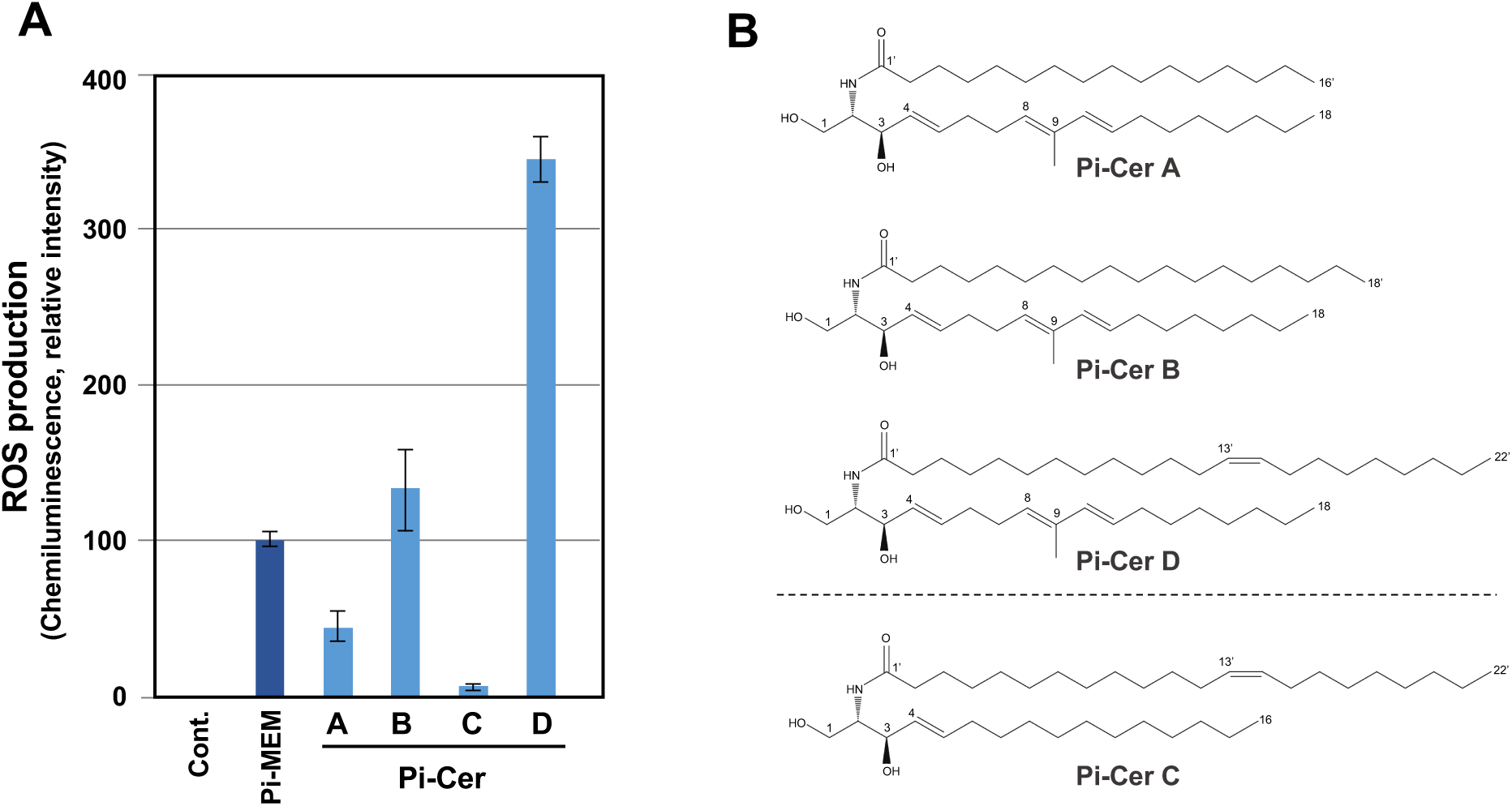
*Phytophthora infestans* ceramide elicitors (Pi-Cer) can induce the production of reactive oxygen species (ROS) in potato suspension cultured cells. (A) Potato suspension cultured cells were treated with 0.3% DMSO (Cont.), 30 µg/ml methanol extract of *P. infestans* mycelia (Pi-MEM), 3 µg/ml Pi-Cer A, B, C or D and production of ROS was detected as L-012 mediated chemiluminescence 3 h after the treatment. Data are means ± SE (n = 3). Scores are chemiluminescence intensities relative to that of Pi-MEM-treated cells at 3 h. Data shown are the representative results of at least 3 separate experiments. (B) Structures of Pi-Cer A, B, C and D. See Supplemental Figure S1 for the procedures of purification of elicitors and Supplemental Figures S2-5, S10-15 and supplemental document for details of their structural analysis.

### Pi-Cers are MAMPs of oomycete pathogens

To investigate whether the substances corresponding to Pi-Cers are contained in other phytopathogenic oomycetes, a modified purification process was applied to partially purify the Pi-Cers from *Pythium aphanidermatum*. *Py. aphanidermatum* is an oomycete plant pathogen with a wide host range, causing seedling damping-off, root and stem rots and blights of a wide range of important crops (e.g. tomato, soybean, cucumber and cotton). Although the composition of Pi-Cers was different from that in *P. infestans*, Pi-Cer A, B, C and D were detected in *Py. aphanidermatum*. On the other hand, Pi-Cers were not detected in several tested fungal plant pathogens, such as *Botrytis cinerea, Fusarium oxysporum* and *Colletotrichum orbiculare* (Fig. S19). These results suggest that the Pi-Cers are molecules specific to oomycetes.

### Pi-Cer D and Pi-CerPE D are elicitors of ROS production in dicot and monocot plants

Because Pi-Cer D and Pi-CerPE D had the highest elicitor activity among Pi-Cers and Pi-CerPEs, respectively, and were also more abundantly purified than the A and B type substances (Fig. S1), Pi-Cer D and Pi-CerPE D were mainly used in the following experiments. In potato suspension-cultured cells, Pi-MEM induced ROS production peaks approx. 3 h after treatment (Fig. S20). The same pattern was observed after Pi-Cer D and Pi-CerPE D treatment (Fig. S20), suggesting that Pi-Cers and Pi-CerPEs could be the major ROS inducing substances in Pi-MEM. Potato leaves were treated with Pi-Cer D, Pi-CerPE D or Pi-MEM by syringe infiltration and induction of ROS production was detected. ROS production was induced by these elicitors with a peak at approx. 12 h after the treatment. Similarly, ROS production was detected in leaves of *A. thaliana* and rice treated with Pi-Cer D or Pi-CerPE D 12 h after the treatment (Fig. 3). Pi-Cer D, but not Pi-Cer C, induced ROS production in *N. benthamiana* (Fig. S21). These results indicated that Pi-Cer D are recognized by both dicot (potato, Arabidopsis, *N. benthamiana*) and monocot (rice) plants as MAMPs.

**Figure 3.**
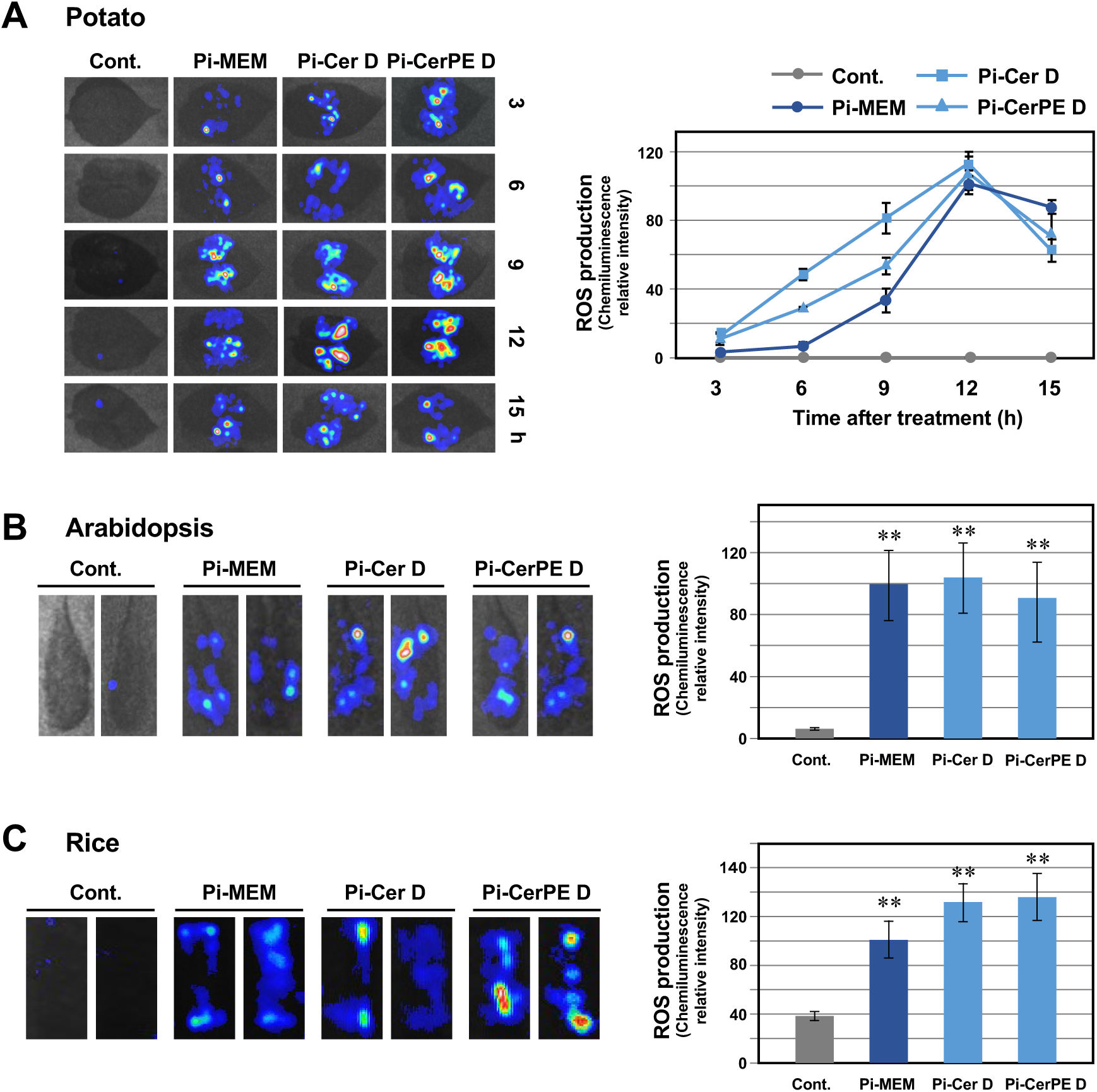
Pi-Cer D and Pi-CerPE D can induce the production of reactive oxygen species (ROS) in leaves of different plant species. Scores shown are chemiluminescence intensities relative to that of Pi-MEM-treated leaves. (A) (Left) Potato leaves were treated with 1% DMSO (Cont.), 100 µg/ml methanol extract of *P. infestans* mycelia (Pi-MEM), 10 µg/ml Pi-Cer D or Pi-CerPE D by syringe infiltration, and production of ROS was detected as L-012 mediated chemiluminescence 3-15 h after the treatment. Data are means ± SE (n = 3). (B) (Left) Arabidopsis thaliana leaves were treated with 1% DMSO (Cont.), 100 µg/ml methanol extract of *P. infestans* (Pi-MEM), 10 µg/ml Pi-Cer D or Pi-CerPE D by syringe infiltration, and production of ROS was detected as L-012 mediated chemiluminescence 12 h after the treatment. Data are means ± SE (n = 3). (C) (Left) Rice leaves were treated with 1% DMSO (Cont.), 100 µg/ml methanol extract of *P. infestans* (Pi-MEM), 10 µg/ml Pi-Cer D or Pi-CerPE D by vacuum infiltration, and production of ROS was detected as L-012 mediated chemiluminescence 12 h after the treatment. Data are means ± SE (n = 3). Data marked with asterisks are significantly different from control as assessed by two-tailed Student’s *t* tests: \*\**P* < 0.01.

### Treatment of Pi-Cer D and Pi-CerPE D enhances the resistance of potato against *P. infestans*

The effect of pre-treatment with ceramide elicitors on the resistance of potato against *P. infestans* was investigated (Fig. 4). Potato leaves were treated with 10 µg/ml Pi-Cer D, Pi-CerPE D or 100 µg/ml Pi-MEM, and inoculated with a zoospore suspension of *P. infestans* at 24 h after the elicitor treatment. Within 3 days after the inoculation, leaves of the control plant showed water-soaked disease symptoms, and the lesions on most leaves extended over the entire leaves within 7 days. In contrast, fewer and smaller spots were developed in leaves treated with Pi-Cer D, Pi-CerPE D or Pi-MEM. Evaluation of disease symptoms up to 7 days after the inoculation indicated that pretreatment with Pi-Cer D and Pi-CerPE D can enhance the resistance of potato leaves against *P. infestans* (Fig. 4). Inoculated leaves were stained with aniline blue to visualize the penetration of *P. infestans* as fluorescent spots of deposited callose beneath the penetration sites at 24 h after the inoculation. Compared with control leaves, fewer fluorescence spots were detected in leaves pretreated with Pi-Cer D, Pi-CerPE D or Pi-MEM elicitors (Fig. 4D), which indicates that Pi-Cer D and Pi-CerPE D can enhance the pre-penetration resistance of potato leaves against *P. infestans*.

**Figure 4.**
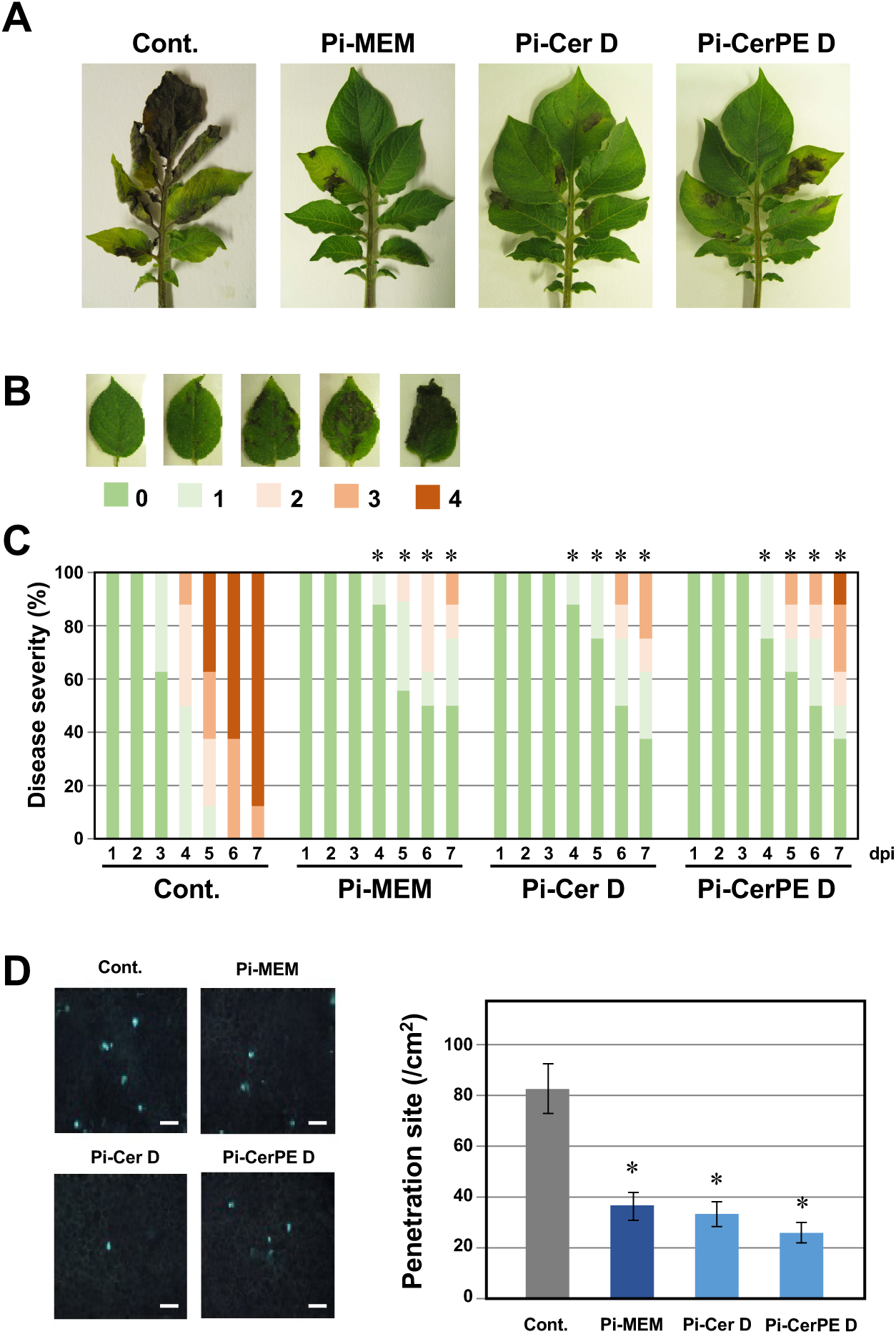
Pretreatment with Pi-Cer D and Pi-CerPE D enhances the resistance of potato leaves against *P. infestans*. Potato leaves were treated with 0.5 % DMSO (Cont.), 100 µg/ml Pi-MEM, 10 µg/ml Pi-Cer D or Pi-CerPE D and incubated for 24 h, and inoculated with a spore suspension of *P. infestans*. (A) Disease symptoms of *P. infestans* on potato leaves treated with DMSO, Pi-MEM, Pi-Cer D or Pi-CerPE D. Photographs were taken 6 days post inoculation. (B) Leaves representative of the disease severities used for classification. (C) Plots showing percentages of potato leaves with disease symptom severities using the classification depicted in (B). Leaves were pretreated with DMSO or elicitors, and disease severity of a subsequent *P. infestans* inoculation was observed from 1 - 7 days post inoculation (dpi). (n = 8). Data marked with asterisks are significantly different from control as assessed by one-tailed Mann–Whitney U-tests: \**P* < 0.05. (D) Left, Penetration sites of *P. infestans* in elicitor treated leaf-discs were detected as callose depositions by aniline blue staining 24 h after inoculation. Bars = 50 µm. Right, Number of fluorescent spots were counted in elicitor-treated leaf discs inoculated with *P. infestans* 24 h after inoculation. Data are means ± SE (n = 3). Data marked with asterisks are significantly different from control as assessed by two-tailed Student’s *t* tests: \**P* < 0.01.

### Isolation of Pi-DAGs as elicitors for the induction of phytoalexin production in potato tuber

In potato tubers, accumulation of phytoalexins (rishitin, lubimin and oxylubimin) was induced by the treatment with the Pi-MEM (Fig. 1). However, treatment of Pi-Cers or Pi-CerPEs did not induce the production of phytoalexins in potato tubers (Fig. S22), which suggested that Pi-MEM includes other elicitors that can induce phytoalexin production. The initial step of our purification protocol consisted of fractioning the butanol-soluble compounds via silica gel flash column chromatography, yielding eight fractions. Of these eight fractions, the third (RM-I-130-3) and sixth (RM-I-130-6) fraction had shown ROS inducing activity and were used for the isolation of Pi-Cers and Pi-CerPEs (Figs. S1 and 22). The same eight fractions were now used to evaluate their ability to induce phytoalexin production in potato tubers. Substantial elicitor activity was only observed in the first fraction (RM-I-130-1) and it was therefore further purified (Fig. S23). Among the five fractions obtained from the following flash column chromatography, the third (ST-I-8-3) and fourth (ST-I-8-4) fraction showed elicitor activity that resulted in phytoalexin production. These fractions were later found to contain mainly 1,3-DAG (diacylglycerol) and 1,2-DAG, respectively. Both ST-I-8-3 and -4 were further separated by HPLC using an ODS column and at least 5 out of 9 resultant fractions showed elicitor activity (Fig. S23). These results indicated that several 1,3- and 1,2-DAGs derived from *P. infestans* are elicitors that can induce the phytoalexin production of potato tubers. Two fractions (ST-I-14-4 and ST-I-23-4) with relatively higher elicitor activity, yield and purity were selected for further structural analyses. Two additional fractions (ST-I-14-8 and ST-I-23-8) which showed almost no elicitor activity were also subjected for structural analyses in order to narrow down the molecular patterns that may be critical for the elicitor activity. Further structural analyses (Figs. S24-27) revealed that the active substances from ST-I-14-4 and ST-I-23-4 were 1,2- and 1,3-DAGs which both contained an eicosapentaenoic acid (EPA) and linoleic acid as fatty acid chains each, and were designated as Pi-DAG A and B, respectively (Figs. 5 and S23). The inactive substances (ST-I-14-8 and ST-I-23-8) were found to be 1,2- and 1,3-DAGs containing palmitic acid and linoleic acid as fatty acid chains, and were designated as Pi-DAGs C and D, respectively. A variety of Pi-DAGs such as Pi-DAG A-D, are also contained in *Py. aphanidermatum*, suggesting that Pi-DAGs are a group of molecules commonly found in oomycete plant pathogens (Fig. S28).

**Figure 5.**
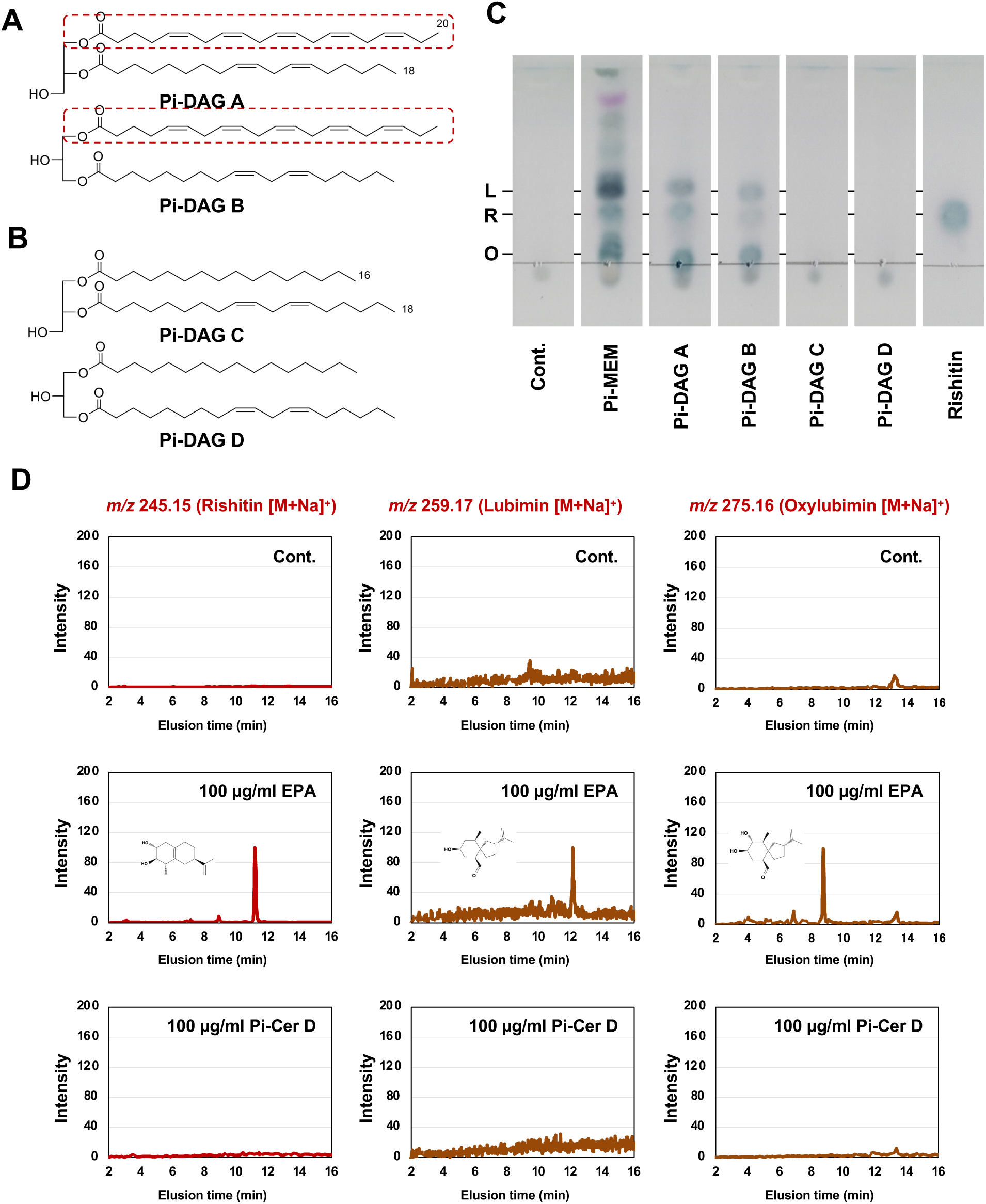
*Phytophthora infestans* diacylglycerol (Pi-DAG) induce the production of phytoalexins in potato tubers. (A) Structures of Pi-DAG A and B. Structures equivalent to eicosapentaenoic acid (EPA) are shown in red, dotted boxes. (B) Structures of Pi-DAG C and D which have significantly weaker elicitor activity compared with Pi-DAG A and B. See Supplemental Figure S22 for the procedures of purification of Pi-DAGs and Supplemental Figures S23-26 for their structural analysis. (C) Potato tubers were treated with 0.3% DMSO (Cont.), 30 µg/ml methanol extract of *P. infestans* mycelium (Pi-MEM), or 100 µg/ml Pi-DAGs, and production of phytoalexins was detected by thin-layer chromatography. L, Lubimin; R, Rishitin; O, Oxylubimin. (D) EPA, but not Pi-Cer D, can induce the production of phytoalexins in potato tubers. Potato tubers were treated with 0.3% DMSO (Cont.), 100 µg/ml EPA or 100 µg/ml Pi-Cer D and produced phytoalexins were detected by LC/MS. Data shown are the representative results of at least 3 separate experiments.

Based on the structural difference of active and inactive Pi-DAGs (Fig. 5A and B), EPA was predicted to be the essential structure for the elicitor activity of Pi-DAGs A and B. Consistently, EPA from *P. infestans* has previously been identified as an elicitor molecule (Bostock et al., 1981). We confirmed that EPA, but not Pi-Cer D, can induce the production of sesquiterpenoid phytoalexins (rishitin and its precursors, lubimin and oxylubimin) in potato tubers (Fig. 5C and D). On the contrary, Pi-Cer D, but not EPA, induced ROS production in potato leaves (Fig. S29), thus indicating that the structurally different oomycete MAMPs, EPA and Pi-Cer D, induce distinctive plant immune responses.

### Treatment of EPA enhances the resistance of potato tuber against *P. infestans*

The effect of treatment with EPA on the resistance of potato tubers against *P. infestans* was evaluated (Fig. 6). Potato tuber discs were treated with 100 µg/ml EPA, and inoculated with a zoospore suspension of *P. infestans* immediately or at 24 h after the EPA treatment. Within 4 days after the inoculation, the growth of *P. infestans* hyphae could be detected on inoculated tubers, and fewer hyphae were developed on tubers treated with EPA (Fig. 6). Conidiophores produced on potato tubers were significantly reduced by EPA treatment, and the effect of EPA treatment on enhanced resistance was even more pronounced when EPA was pretreated 24 h before the inoculation.

**Figure 6.**
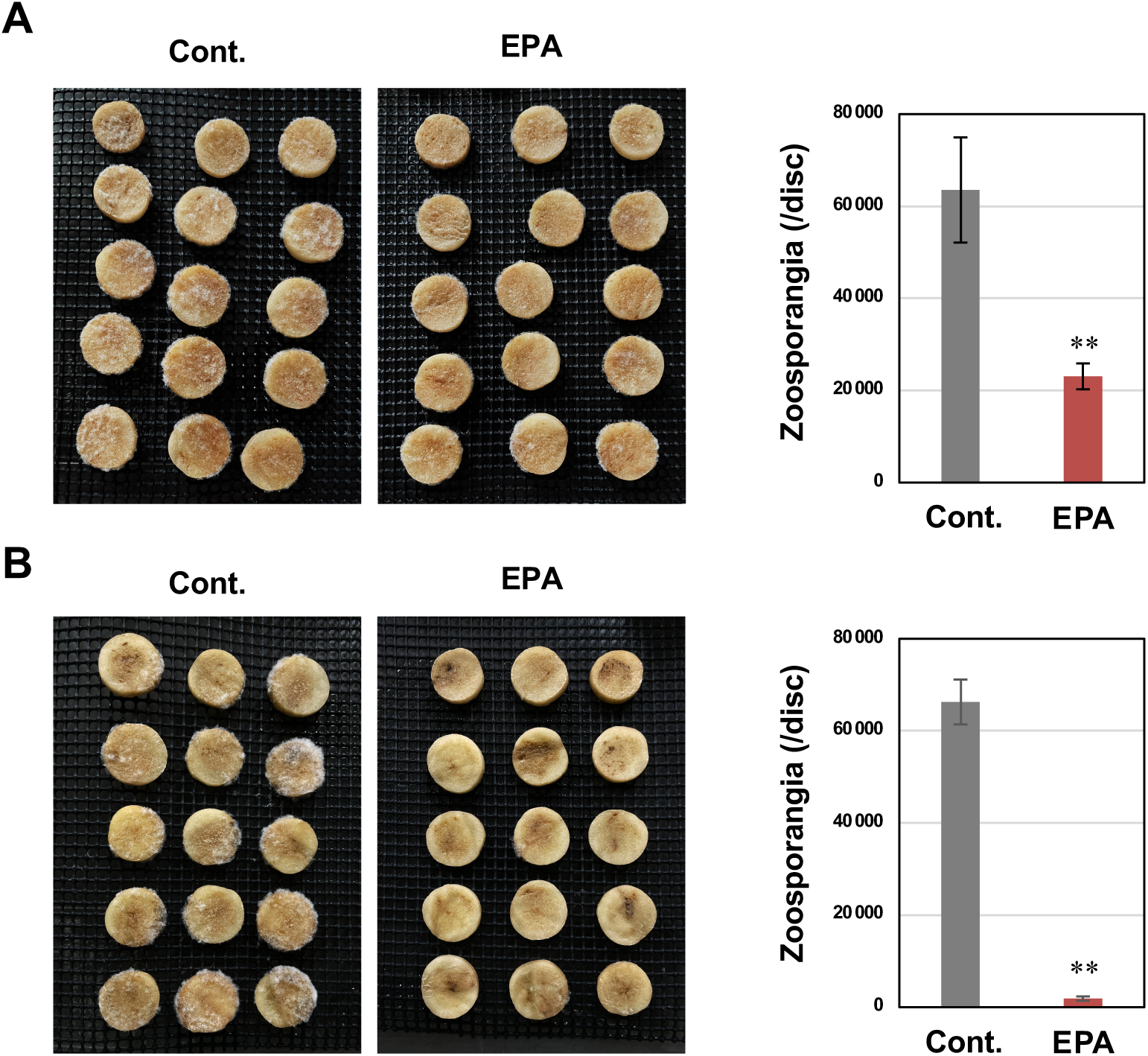
Treatment with EPA enhances the resistance of potato tubers against *P. infestans*. Potato tuber discs (cv. Irish cobbler, 1 cm diameter) were treated with 50 µl of 0.1 % DMSO (Cont.) or 100 µg/ml EPA and inoculated with a spore suspension of *P. infestans* (A) immediately or (B) 1 day after treatment. (left) Growth of *P. infestans* on potato tubers. Photographs were taken at 4 days post inoculation (dpi). (right) No. of conidiophores produced on potato tubers were counted at 4 dpi (n = 16). Data marked with asterisks are significantly different from control as assessed by two-tailed Student’s *t* tests: \*\**P* < 0.01. Data shown are the representative results of 2 separate experiments.

### Identification of the essential structure of unsaturated fatty acid to be recognized as MAMPs in potato

EPA is an omega-3 polyunsaturated fatty acid with 20 carbon chain length and 5 double bonds (20:5, Δ5,8,11,14,17, ω-3) (Fig. 7A). The primary unsaturated fatty acids contained in plants are linoleic acid (18:2, Δ9,12, ω-6) and α-linolenic acid (18:3, Δ9,12,15, ω-3) (Cahoon and Li-Beisson, 2020), thus it is presumed that plants recognize the characteristic structure of unsaturated fatty acids not contained in plants as MAMPs. To identify the essential molecular pattern of unsaturated fatty acid to be recognized as a MAMP, structurally related unsaturated fatty acids were tested for their elicitor activity on potato. Two days after the treatment of various fatty acids at 100 µg/ml on potato tuber, accumulated rishitin was extracted and quantified by GC/MS. As predicted, three unsaturated fatty acids derived from plants, linoleic acid (LA), α-linolenic acid (ALA) and γ-linolenic acid (18:3, Δ6,9,12, ω-6, GLA) had no elicitor activity. Treatment of arachidonic acid (20:4, Δ5,8,11,14, ω-6, ALA), which was previously reported as an elicitor (Bostock et al., 1981), induced rishitin production, but other structurally related unsaturated fatty acid, namely eicosadienoic acid (20:2, Δ11,14, ω-6, EDA), eicosatrienoic acid (20:3, Δ8,11,14, ω-6, ETA), mead acid (20:3, Δ5,8,11, ω-9, MA) and docosahexaenoic acid (22:6, Δ4,7,10,13,16,19, ω-3, DHA) did not show elicitor activity on potato, even though DHA includes conjugated double bonds or MA shares an identical structure to AA and EPA at the end of the carboxylic acid group (Fig. 7). Thus, 5,8,11,14-tetraene-type fatty acid (5,8,11,14-TEFA) is presumed to be the essential structure recognized by potato as a MAMP.

**Figure 7.**
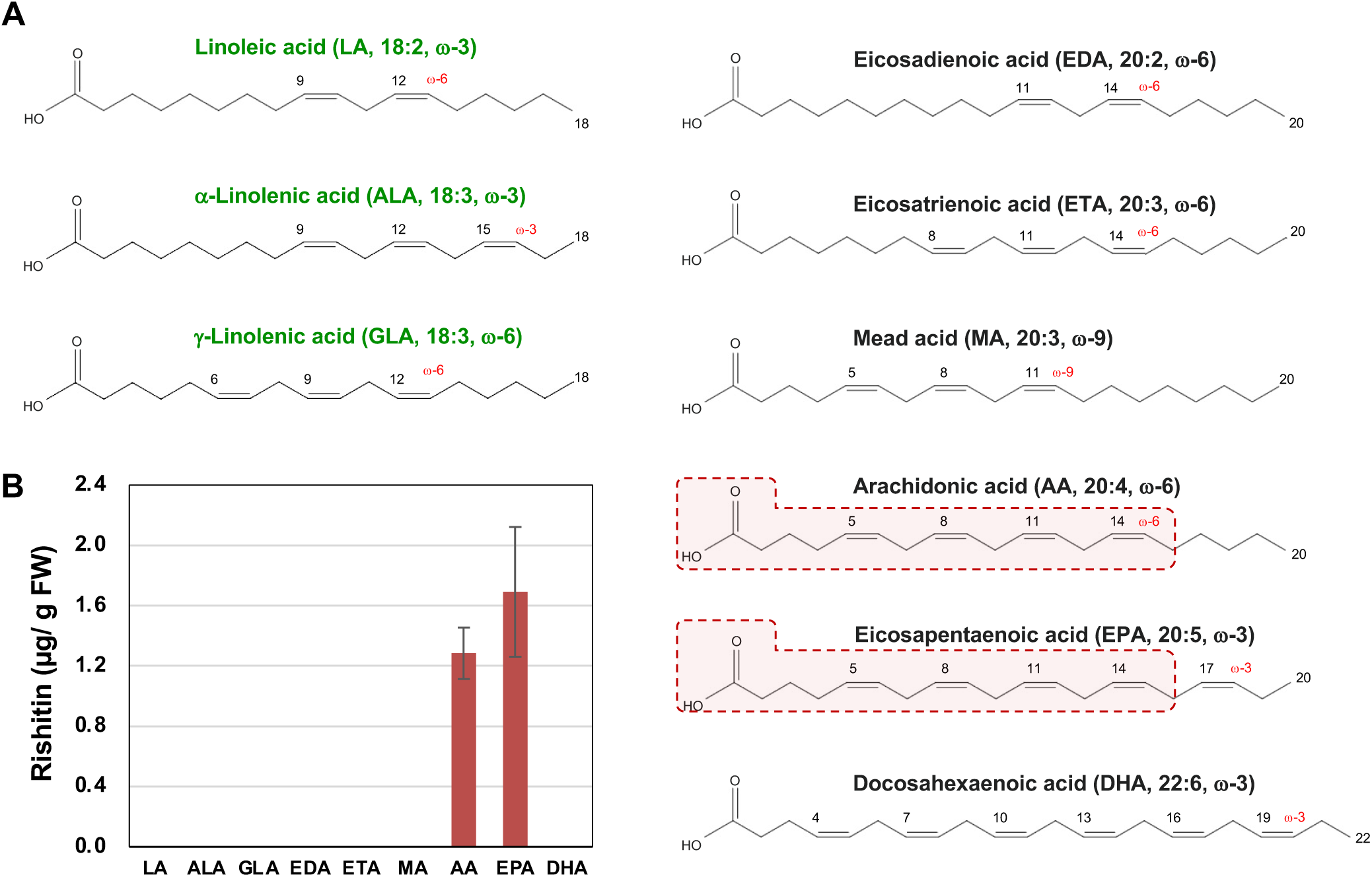
Comparison of elicitor activity of polyunsaturated fatty acids for induction of rishitin production in potato tubers. (A) Structures of polyunsaturated fatty acids used in this study. Fatty acids in green letters indicate those derived from plant. (B) Potato tubers were treated with indicated polyunsaturated fatty acids (100 µg/ml) and rishitin was extracted 2 days after the treatment. Produced rishitin was quantified by GC/MS (n = 6 for AA and EPA, n = 3 for LA, ALA, GLA, EDA, ETA, MA. n = 12 for DHA, biological replicates from 3 separate experiments).

### Distinctive sets of genes were upregulated in Arabidopsis treated with EPA, Pi-Cer D and their mixture

To analyze the differences in plant responses resulting from the recognition of EPA and Pi-Cer D, an Arabidopsis pWRKY33-LUC transformant was treated with 100 µg/ml EPA and Pi-Cer D. While Pi-Cer D can induce the transient activation of the *WRKY33* promoter, only marginal induction of the *WRKY33* promoter was detected by the treatment with EPA (Fig. 8A). Similarly, callose deposition, a form of penetration resistance (Ellinger et al., 2013), was induced by either Pi-Cer D or EPA treatment on leaves of Arabidopsis seedlings, but Pi-Cer D induced significantly greater number of depositions (Fig. S30). RNAseq analysis was performed for Arabidopsis seedlings 12 h after the treatment with 100 µg/ml EPA, Pi-Cer D or a mixture of EPA and Pi-Cer D. When the number of upregulated genes was compared under the same criteria (Log2 fold change ≥2, p ≤0.05), Pi-Cer D had only 211 upregulated genes, compared to 1,422 genes for EPA treatment. Unexpectedly, simultaneous treatment with Pi-Cer D and EPA did not result in an expression pattern that would be expected by the mere combination of either treatment. Instead, a significantly a smaller number of genes (439 genes) was upregulated by the mixture of EPA and Pi-Cer D compared with the single treatment of EPA, and 76.8% of genes upregulated by the mixture of EPA and Pi-Cer D (337 genes) were specifically induced by the co-treatment of both elicitors (Fig. 8B). Similarly, clustering analysis of significantly up- or down-regulated genes further revealed that treatment with EPA or Pi-Cer D leads to distinct gene expression patterns. While some genes respond similarly to either treatment, other genes are induced exclusively by only EPA or Pi-Cer D treatments (Fig. 8C). Expression of some genes was significantly upregulated only by simultaneous treatment with both elicitors (Fig. S31A), whereas other genes induced by EPA were attenuated by the co-treatment with Pi-Cer D (or vice versa) (Fig. S31B and C), resulting in an expression pattern different from either single treatment.

**Figure. 8.**
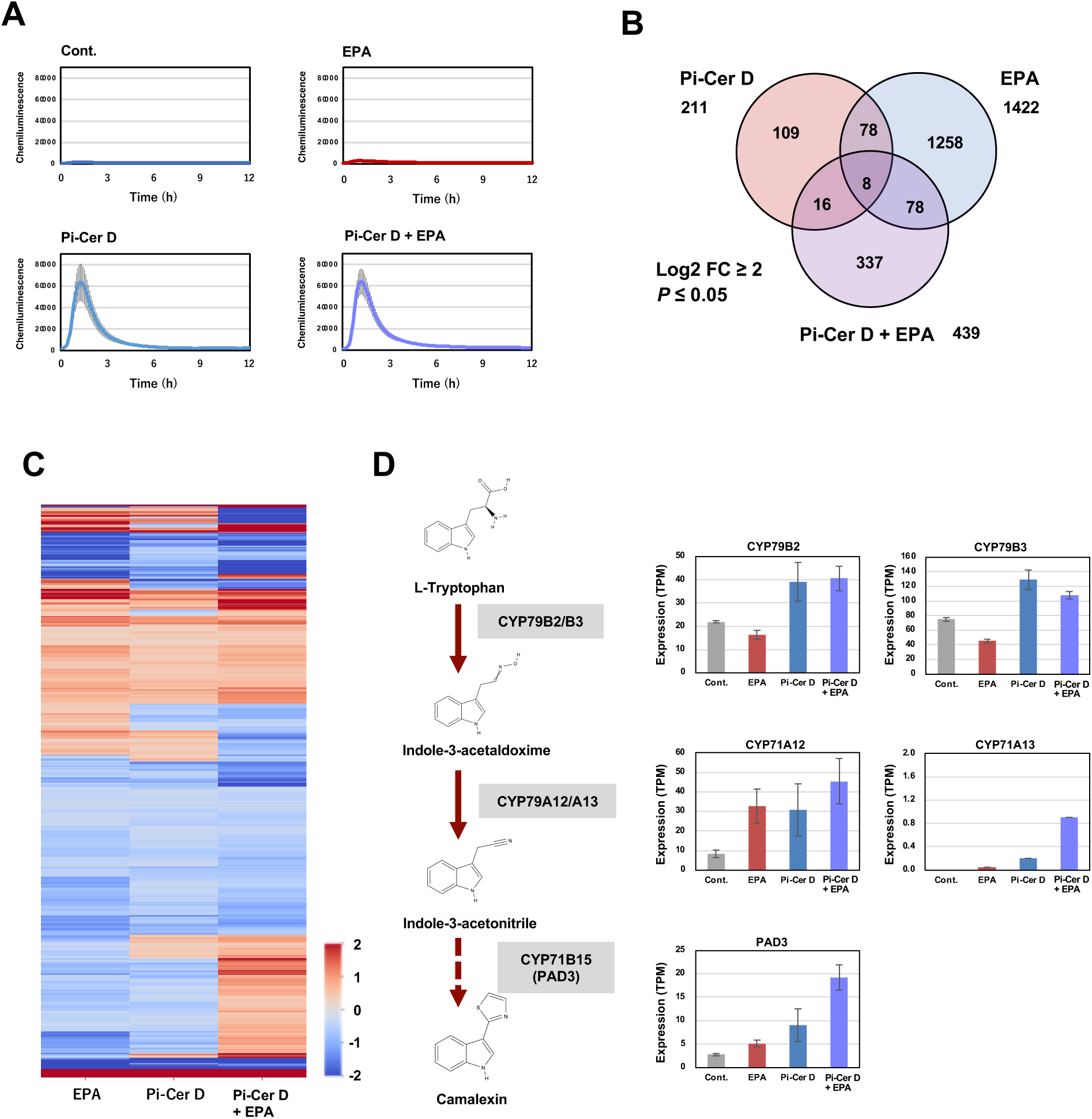
Distinctive sets of Arabidopsis genes were upregulated by the treatment with EPA and Pi-Cer D. (A) Arabidopsis transformant pWRKY33-LUC containing LUC transgene under the control of AtWRKY33 (AT2G38470.1) promoter was treated with 100 µg/ml EPA, Pi-Cer D or a mixture of 100 µg/ml EPA and Pi-Cer D. Chemiluminescence was monitored for 12 h after the treatment. (B) Venn diagram representing the up-regulated differentially expressed genes (DEGs) in Arabidopsis treated with 100 µg/ml EPA, 100 µg/ml Pi-Cer D, or a mixture of 100 µg/ml EPA and 100 µg/ml Pi-Cer D for 12 h, which are selected based on TPM ≥ 1, log2 fold change ≥ 2 and *P* ≤ 0.05. (n = 3). (C) Heatmap analysis of differentially expressed genes for the elicitors treatments. The color bar represents the fold change value (log2) with *P* ≤ 0.05. (D) Expression profile of Arabidopsis genes for camalexin biosynthesis after treatment with 100 µg/ml EPA, Pi-Cer D or a mixture of 100 µg/ml EPA and Pi-Cer D.

To interpret the overall influence of treatment with EPA, Pi-Cer D or their mixture, upregulated genes were assigned to gene ontology (GO) terms using the analysis tool PANTHER (Thomas et al., 2022) for GO enrichment analysis (Supplemental Fig. S32). Biological processes (BP) enhanced by Pi-Cer D treatment include categories related to plant defense against microorganisms such as “Defense response to fungus (GO:0050832)”, “Defense response to bacterium (GO:0042742)”. EPA treatment also enhanced “Defense response to fungus (GO:0050832)”, but also induced distinctive categories such as “Oligopeptide transport (GO:0006857)”, “Detoxification (GO:0098754)” and “Inorganic ion homeostasis (GO:0098771)”. It should be noted that although the number of genes induced by EPA treatment (1422 genes) used for GO analysis was approx. 7-fold compared to that of Pi-Cer D (211 genes), the number of enriched GO terms hit was less than 2.2-fold (15 terms for Pi-Cer D and 33 terms for EPA, Supplemental Fig. S32), indicating that the genes induced by solo treatment of EPA are not focused on a specific pathway or response. More importantly, GO analysis using 432 genes induced by simultaneous treatment with EPA and Pi-Cer D detected the relatively larger enrichment of much more GO terms (145 terms), including GOs involved in disease resistance, such as “Camalexin biosynthetic process (GO:0010120)” (also see Fig. 8D), “Response to chitin (GO:0010200)”, “Innate immune response (GO:0045087)”, “Defense response to fungus (GO:0050832)”, “Defense response to bacterium (GO:0042742)” and so on. This result could indicate that the simultaneous recognition of multiple MAMPs by plants better defines the appropriate choices of the stress responses in the plant cells.

## DISCUSSION

As the major autotroph on land, terrestrial plants are suitable predation targets for many organisms, including bacteria, fungi, oomycetes, insects, animals, or even some parasitic plants, which consume the organic compounds produced by plants. Therefore, since their ancestral phyla, defense against various types of foreign enemies has been of utmost importance. While plants form surface physical barriers and superficially accumulate antimicrobial substances on their surfaces as the first line of defense in general, they also need to activate greater resistance when attacked by microorganisms, and recognition of so-called MAMPs is the key event to activate the next line of plant disease resistance.

Characteristic surface structures of microorganisms are recognized by plant cells as MAMPs. The major components of fungal cell walls (such as chitin and β-glucan) and the secretory proteins (NEP and elicitins), as well as the bacterial flagellin and lipopolysaccharides are typical examples of MAMPs localized at the surface (Jones and Takemoto, 2004; Ngou et al., 2022). Flagellin, lipopolysaccharides, chitin and β-glucan are recognized as MAMPs for induction of innate immunity in animal cells by distinct mechanisms from plants and are the results of convergent evolution (Takeuchi et al., 1999; Hayashi et al., 2001, He et al., 2021). Whereas some MAMPs derived from internal molecules, such as bacterial elongation factor EF-Tu or extracellular DNA and RNA, are released after disruption of the microbial cells, and then recognized by the plant cells (Kunze et al., 2004; Bhat and Ryu, 2016). When plant cell walls are damaged by pathogen attack, fragments of oligosaccharides (such as cello-oligosaccharides and xylo-oligosaccharides) are recognized by cell surface receptors as DAMPs (Martín-Dacal et al., 2023; Pring et al., 2023).

In this study, lipophilic MAMPs derived from microbial cell membrane, Pi-Cers and Pi-DAGs, were purified from a representative phytopathogenic oomycete, *P. infestans*, for their activity in inducing ROS production and phytoalexin accumulation in potato. These substances were also contained in polyxenous oomycete pathogen *Pythium aphanidermatum* (Figs. S19, S28) and other oomycetes (Moreau et al., 1998, Fernandes et al., 2019). Pi-Cers and Pi-DAGs (or EPA) activated resistance responses in potato, Arabidopsis and other plant species, indicating that they are oomycete MAMPs recognized by plants (Fig. 9, Bostock et al., 2011).

**Figure 9.**
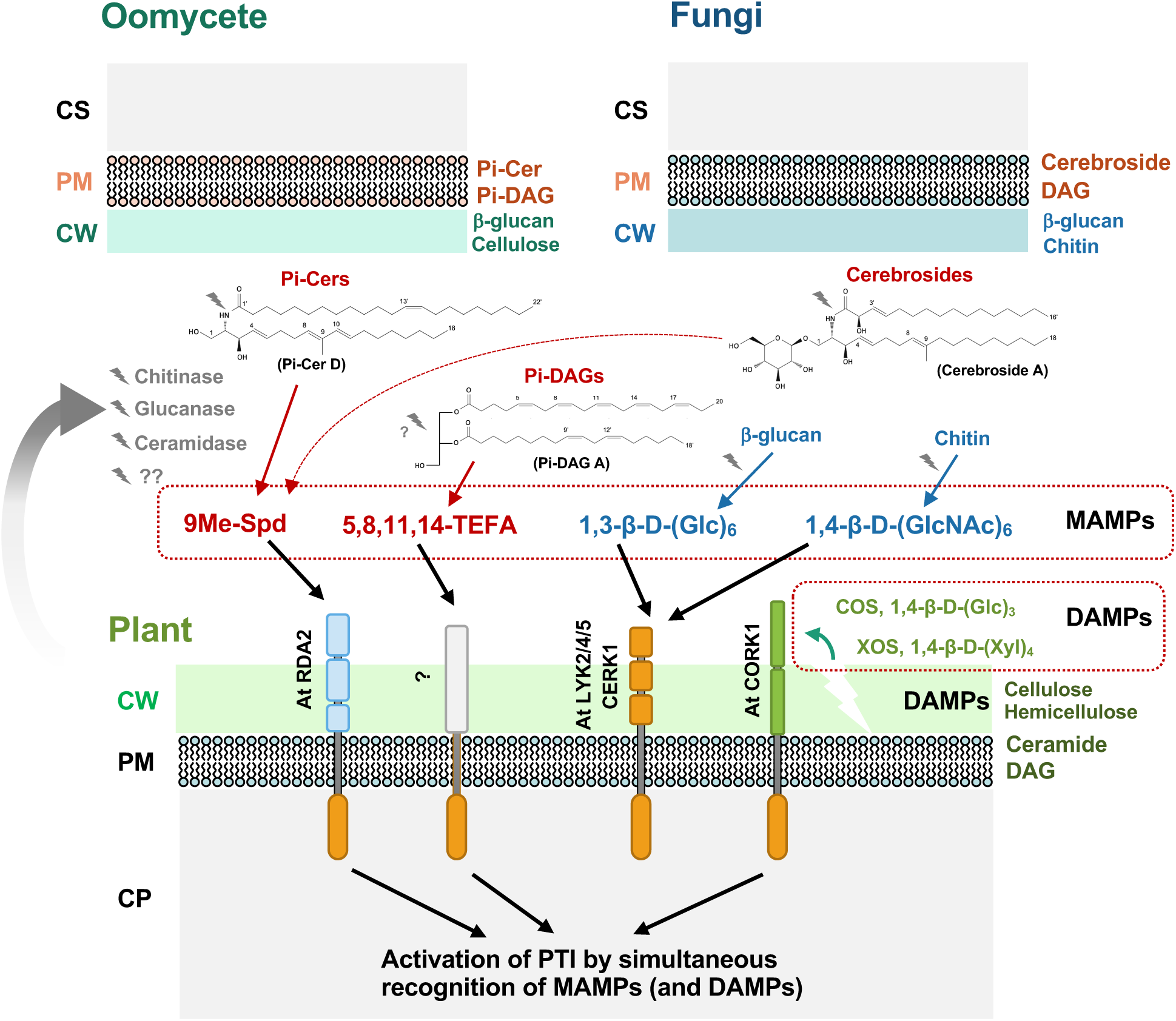
A model of Pattern-triggered immunity (PTI) activation by simultaneous recognition of multiple MAMPs (Microbe-associated molecular patterns) and DAMPs (Damage-associated molecular patterns). Plants secrete enzymes into the apoplast to release MAMPs from the pathogen. 9Me-Spd, 9-methyl-4,8-sphingadienine; 5,8,11,14-TEFA, 5,8,11,14-tetraene-type fatty acid; COS, Cellooligosaccharide; XOS, Xylooligosaccharide; CW, Cell wall; PM, Plasma membrane; CP, Cytoplasm.

### Two characteristic structures of microbial plasma membrane lipids are recognized by plant cells

Ceramides and diacylglycerols are principal structural elements of cell membranes. Ceramides are composed of a sphingoid base and a fatty acid joined via an amide bond. The most evident structural difference between active Pi-Cers and inactive Pi-Cer C is the 8,10-diene and methyl-branching at C-9 in the sphingoid base (Fig. 2). The same structure was found in ceramide-phosphoethanolamines Pi-CerPE A, B and D purified from other fractions (Figs. S1, S17). In a previous report, we indicated that 9-methyl branching and the 4,8-double bond structure of the sphingoid base (9Me-Spd) represent the epitope structure for its elicitor activity in Arabidopsis (Kato et al., 2022). Although Pi-Cers are not detected in fungal pathogens (Fig. S19), cerebrosides have been identified as elicitor molecules from rice blast pathogen *Pyricularia oryzae* (Koga et al., 1998, Fig. 9). Cerebroside elicitors isolated from *P. oryzae* and other fungal pathogens also contain epitope structure 9Me-Spd (Umemura et al., 2000). Given that 9Me-Spd is commonly found in (evolutionally distant) oomycete and fungal species but essentially not found in plants (Moreau et al., 1998; Sperling and Heinz, 2003, Jiang et al., 2021), this structure appears to be a molecular pattern that can be leveraged by the plant to recognize potentially pathogenic microorganisms. Knockout of a gene for sphingolipid C-9 methyltransferase *FgMT2* of *Fusarium graminearum* (which caused an approx. 70% reduction of C-9 methylation in glucosylceramides) showed reduced growth and pathogenicity on wheat and Arabidopsis (Ramamoorthy et al., 2009). In *Cryptococcus neoformans*, Sphingolipid C9 methyltransferase SMT1 was shown to be involved in maintaining membrane lipid bilayer rigidity and pathogenicity in mice (Singh et al., 2012). These reports indicate that 9Me-Spd is an essential structure for the fungal membrane, which fungi may not be able to modify with ease. By mutagenesis of the Arabidopsis pWRKY33-LUC marker line and screening of mutants insensitive to Pi-Cer D, Kato et al. (2022) identified that a lectin receptor kinase RDA2 is the receptor of 9Me-Spd in Arabidopsis. Moreover, cleavage of Pi-Cer D by an apoplastic ceramidase NCER2 is essential for perception of 9-methyl sphingoid base by RDA2 in Arabidopsis, indicating that the excision of microbial components by the plant is an important process for the activation of pattern-triggered immunity (PTI) of plants (Fig. 9).

Diacylglycerols consist of two fatty acid chains linked to a glycerol molecule through covalent ester bonds. Pi-DAGs A and B, which have elicitor activity and induce potato phytoalexin production, are 1,2- and 1,3-diacylglycerols, respectively, and share the same fatty acids they consisted of were the same. Comparison of Pi-DAGs A and B with Pi-DAGs C and D, which had no elicitor activity (Fig. 5), suggested that a structure containing five unsaturated fatty acids is the site recognized by the plant. In most plants, the predominant unsaturated fatty acids are oleic acid (18:1, ω-9), linoleic acid (18:2, ω- 6), and α-linolenic acid (18:3, ω-3) (Harwood, 1988), as well as γ-linolenate (18:3, ω-6), which is found in a few seed oils (Nykiforuk et al., 2012). Natural unsaturated fatty acids in plants contain one to three conjugated double bonds, and these unsaturated fatty acids showed no elicitor activity on potato tuber (Fig. 7). In contrast, EPA, which is a pentaene fatty acid (20:5, ω-3) contained in bacteria, fungi and oomycete (Adarme-Vega et al., 2012), was previously reported as an elicitor of phytoalexin production in potato by Bostock et al. 1981. In this study, we identified the essential and sufficient structure of microbe-derived unsaturated fatty acids that act as MAMP. Although Bostock et al. (1981) also reported that docosahexaenoic acid (DHA, 22:6, ω-3) had elicitor activity, experiments conducted in this study with three different lots of DHA showed that treatment with DHA has no elicitor activity for the induction of phytoalexin production in potato (Fig. 7), suggesting that four or more conjugated double bonds alone are not sufficient to be recognized as a MAMP by plants. Given that arachidonic acid (20:4, ω-6), but not mead acid (20:3, ω-9), showed elicitor activity on potato, 5,8,11,14-tetraene-type fatty acid (5,8,11,14-TEFA) is presumed to be the essential structure recognized by plants as a MAMP. Collectively, plants activate PTI by recognizing characteristic microbial characteristic structures in the cell membrane. Since plants secrete enzymes that partially degrade MAMPs such as Pi-Cers, chitin, and β-glucans, such secreted enzyme (e.g. phospholipase or esterase) might also be involved in the degradation of Pi-DAGs via the exposure of the 5,8,11,14-TEFA structure (Fig. 9).

### Simultaneous recognition of two lipophilic MAMPs induces distinctive plant immune responses

Over the course of infection by plant pathogens, plants are exposed to a series of MAMPs, at different times of the infection process. While patterns on the microbial cell wall and cell membrane are expected to be recognized first, intracellular components and DAMPs (e.g. oligosaccharides derived from plant cell walls, Pring et al., 2023) should be released at later stages of the infection. It is presumed that plants are able to utilize these temporal cues to activate a response better suited to the disease progression. Since we demonstrated that co-exposure to two MAMPs can result in a different defense response, plants may leverage this mechanism to infer information of the disease progression and fine-tune their response.

Two MAMPs derived from the oomycete membrane were isolated in this study, each activating different resistance responses in potato, namely ROS generation and phytoalexin accumulation respectively, indicating that distinct plant signaling is activated by these two MAMPs. In Arabidopsis, Pi-Cer D treatment activates the promoter of *AtWRKY33*, a central activator of disease resistance (Zheng et al., 2006), while EPA hardly activates *AtWRKY33*. Consistently, distinctive sets of genes are induced by the treatment with these two elicitors, though GO terms for up-regulated genes after the treatment with Pi-Cer D or EPA are both includes terms categorized under plant disease resistance. Importantly, simultaneous treatment with these two MAMPs was shown to activate GO categories related to disease resistance that were not triggered by the single treatment (Fig. S32). These results seem to indicate that plants may combine the signals from simultaneously recognized MAMPs to specifically adapt the defense response to a particular type of pathogen. Insight into how these elicitor signals are processed, as well as how the response pathways of both elicitors affect one another, will require further analysis. If additional MAMPs and DAMPs activating different signaling systems were to be determined, we might be able to devise effective combinations that effectively activate resistance. Such MAMP/DAMP recipes might be tailored to act as efficient and stable resistance inducers for use in agricultural production (Pring et al., 2023).

## MATERIALS AND METHODS

### Biological materials, growth conditions and inoculation

Potato suspension-cultured cells were prepared from calluses induced from potato tuber discs (cv. Sayaka carrying the *R1* and *R3* genes) in callus-inducing medium [1 ml/l B5-vitamin (100 g/l Myo-inositol, 2.5 g/l glycine, 500 mg/l thiamine-HCl, 500 mg/l pyridoxine-HCl, 5 g/l nicotinic acid, 50 mg/l biotin, 50 mg/l folic acid), 2 mg/l 1-naphthaleneacetic acid (NAA), 0.5 mg/l kinetin, 3% sucrose, 0.2% phytagel and Murashige-Skoog basal medium], under dark condition. Suspension-cultured potato cells were grown by agitation at 130 rpm, 23°C in 95 ml of Murashige-Skoog medium supplemented with 30 mg/ml sucrose, 1 mg/l thiamine, 100 mg/l myo-inositol, 200 mg/l KH_2_PO_4_, and 0.2 mg/l 2,4-dichlorophenoxyacetic acid. Cells were sub-cultured every week and used for experiments 4-5 days after the subculturing. Potato, rice and Arabidopsis (Col-0) plants were grown at 23-25°C under a 16 h photoperiod and an 8 h dark period in an environmentally controlled growth room. Tubers of potato (cv. Rishiri) carrying the *R1* gene were stored at 4°C until use. *P. infestans* isolate 08YD1 and PI1234-1 (Shibata et al., 2010, 2011) was maintained on rye-media at 10°C. Inoculation of potato leaves with a zoospore suspension of *P. infestans* was performed as described previously (Shibata et al., 2016). Inoculation of potato tubers with *P. infestans* was performed as follows. Potato tuber discs (cv. Irish cobbler, 1 cm diameter, 3 mm thick) were treated with 50 µl of 0.1 % DMSO (Cont.) or 100 µg/ml EPA and inoculated with a spore suspension (2 x 10^5^ zoospores/ml) of *P. infestans* immediately or 1 day after EPA treatment. Potato tubers at 4 days after inoculation were washed in 1 ml water and numbers of conidiophores in 2 µl aliquots were counted. *Pythium aphanidermatum* strain Pyaph isolated from cucumber, *Botrytis cinerea* strain AI18 isolated from strawberry leaves (Kuroyanagi et al., 2022), *Fusarium oxysporum* f. sp. *melonis* strain Mel02010 (Namiki et al., 1998) and *Colletotrichum orbiculare* strain 104-T (Ishida and Akai 1969) were grown on potato dextrose agar (PDA) at 25°C and maintained at -80°C in 20% glycerol. *N. benthamiana* line SNPB-A5 (Shibata et al., 2016) was grown in a growth room at 23°C with 16 h of light per day.

### Preparation of methanol extract of *Phytophthora infestans* mycelia (Pi-MEM)

The *Phytophthora infestans* isolates PI1234-1 were grown on rye-seed extract agar medium in test tubes at 18°C in dark condition for 2 weeks. Parts of the growing mycelia were placed in 100 ml flasks containing 20 ml of rye liquid nutrient medium of rye seed-extract (60 g rye seed), 20 g of sucrose and 2 g of yeast extract per 1 liter and incubated in the dark for 2 weeks at 18°C to allow the growth of the mycelia. The mycelial mats grown in the liquid medium were washed thoroughly with water. To remove excess water, water was filtered through filter paper (Toyoroshi No. 2) under reduced pressure and the tissue then frozen at -20°C. The collected mycelia of *P. infestans* were ground in liquid nitrogen by mortar and pestle. The ground mycelia were transferred to a 50 ml tube containing methanol (10 ml/g mycelia). The mycelia suspension was finely grounded using a polytron type homogenizer (HG30, Hitachi Koki, Japan) for 2 min. After centrifugation at 4°C, 3000 × g for 30 min, the supernatant was collected and dried using an evaporator and used as Pi-MEM elicitor. Further procedures for the purification of Pi-Cers and Pi-DAGs from Pi-MEM and their structural analyses with spectral data are described in the supplemental document. Mycelia of *Pythium aphanidermatum*, *B. cinerea*, *F. oxysporum* f. sp. *melonis* and *C. orbiculare* were grown in 50 ml of potato dextrose broth (PDB) in 100 ml flasks at 25°C with gentle shaking.

### Measurement of ROS production in potato suspension-cultured cells

The relative intensity of ROS generation in potato suspension-cultured cells was measured by counting photons from L-012-mediated chemiluminescence. The potato suspension-cultured cells (50 mg/ml) were washed with the assay buffer (175 mM mannitol, 50 mM MES-KOH, 0.5 mM CaCl_2_ and 0.5 mM K_2_SO_4_, pH 5.7) twice for removal of the liquid culture medium. For the detection of ROS produced in cultured cells, the cells were resuspended in assay buffer, equilibrated for 1 h at 100 rpm at 23°C. Cells were then treated with elicitors and incubated under the same condition for 3 h. After the incubation, chemiluminescence was measured using 20 mM L-012 (Wako Pure Chemical, Osaka, Japan) in a multimode microplate reader Mithras LB940 (Berthold Technologies, Bad Wildbad, Germany).

### Measurement of ROS production in plant leaves

The relative intensity of ROS production was determined by counting photons from L-012-mediated chemiluminescence. To detect the ROS production in potato, *N. benthamiana,* Arabidopsis and rice leaves, 0.5 mM L-012 in 10 mM MOPS-KOH (pH 7.4) was allowed to infiltrate to the intercellular space of leaves. For potato, *N. benthamiana* and Arabidopsis, a syringe without a needle was used for leaf infiltration, whereas vacuum infiltration was used for rice. Chemiluminescence was monitored using a photon image processor equipped with a sensitive CCD camera in the dark chamber at 20°C (Aquacosmos 2.5; Hamamatsu Photonics, Shizuoka, Japan) and quantified using the U7501 program (Hamamatsu Photonics). For the data shown in Supplemental Fig. S21 and S29, chemiluminescence was monitored using Lumino Graph II EM (ATTO, Tokyo, Japan).

### Detection of potato phytoalexins by thin-layer chromatography (TLC)

Potato phytoalexins, exuded from potato tissue, were extracted with ethyl acetate as described previously (Noritake et al., 1996). The extract was separated on TLC plates (TLC aluminum sheet of silica gel 60, Merck, Whitehouse Station, NJ, USA), which were developed with cyclohexane:ethyl acetate (1:1, v/v) and visualized by spraying with sulfuric acid containing 0.5% vanillin followed by heating at 120°C.

### Detection of potato phytoalexins by liquid chromatography/mass spectrometry (LC/MS) and as chromatography/mass spectrometer (GC/MS)

Potato tuber discs (2 cm in diameter and approx. 4 mm thick) were prepared and incubated in a humidified chamber in dark at 23° C for 24 h before treatment with elicitors. The upper side of incubated potato tubers was then treated with 100 µl of elicitor solution and further incubated at 23°C in the dark for 48 h before the extraction of phytoalexins. Four tuber disks/sample were immersed in 5 ml of ethyl acetate and shaken for 1 h, then the organic solvent was collected and evaporated. The residual material was redissolved in 100 µl of 50% (v/v) acetonitrile and 2 µl was injected for the analysis with LC/MS (6520 Accurate-Mass Q-TOF connected to Agilent 1100 Series HPLC, Agilent Technologies, Santa Clara, CA, USA) with a Cadenza CD-C18 column (i. d. 3 x 50 mm, Imtakt, Kyoto, Japan) as previously described (Imano et al., 2022). Detection and quantification of rishitin produced in potato tuber by GC/MS using an Agilent Technologies 7890A GC System with a DuraBond Ultra Inert column (length 30 m; diameter 0.25 mm; film 0.25 µm, Agilent Technologies, Santa Clara, CA, USA) as previously described (Camanga et al., 2020).

### Measuring the activation of *AtWRKY33* promoter

Seeds of Arabidopsis containing *Luciferase* marker gene under the control of *AtWRKY33* (AT2G38470) promoter (Kato et al., 2022) were surface sterilized with 3% hydrogen peroxide and 50 % ethanol for 1 min with gentle shaking, washed with sterilized water, and then individual sterilized seeds were placed in separate wells of a 96 microwell plate containing 150 µl of Murashige and Skoog (MS) liquid medium (1/2 MS salts, 0.05% [w/v] MES, 0.5% [w/v] Sucrose, adjusted to pH 5.8 with NaOH) with 50 µM D-Luciferin potassium salt (Biosynth Carbosynth, Compton, UK), covered with a clear plastic cover. Plates containing Arabidopsis seeds were then incubated in a growth chamber for approx. 12 days at 23°C with 24 h light. After the treatment of elicitors, chemiluminescence intensity derived from expressed luciferase was measured using Mithras LB940 (Berthold) for 12 h.

### RNA-seq and gene ontology enrichment analysis

Total RNA was extracted from 6 seedlings/sample of 10-days old Arabidopsis 24 h after the treatment with elicitors using the RNeasy Plant Mini Kit (QIAGEN, Hilden, Germany). Libraries were constructed using KAPA mRNA Capture Kit (Roche, Basel, Switzerland) and MGIEasy RNA Directional Library Prep Set (MGI, Shenzhen, China), and sequenced on DNBSEQ-G400RS (MGI) with 150 bp paired-end protocol. The RNA-seq reads were filtered using trim-galore v.0.6.6 (Martin, 2011, bioinformatics.babraham.ac.uk) and mapped to the Arabidopsis genome (TAIR 10.1; RefSeq: GCF_000001735.4) using HISAT2 v.2.2.1 (Kim et al., 2019) and assembled via StringTie v.2.1.7 (Kovaka et al., 2019). Significant differential expression was determined using DESeq2 v.1.32.0 (Love et al., 2014). All software used during RNA-seq analysis was run with default settings. The expression profile was calculated from the log2-fold expressions using the clustermap function from seaborn v. 0.11.1 (Waskom, 2021). Gene ontology (GO) enrichment analysis was performed using the PANTHER statistical overrepresentation test (http://pantherdb.org; Version 17.0, Thomas et al., 2022) with default settings (Fisher’s exact test, False discovery rate (FDR) < 0.05). The heatmap was plotted using SRplot (https://www.bioinformatics.com.cn/en). RNA-seq data reported in this work are available in GenBank under the accession numbers DRA017453.

## ACKNOWLEDGMENTS

We thank Ms. Kayo Shirai (Hokkaido Central Agricultural Experiment Station, Japan) and Dr. Seishi Akino (Hokkaido University, Japan) for providing *P. infestans* isolate, Dr. Kenji Asano, Dr. Kotaro Akai and Mr. Seiji Tamiya (National Agricultural Research Center for Hokkaido Region, Japan) and Mr. Yasuki Tahara (Nagoya University, Japan) for providing tubers of potato cultivars, and Mr. Masamu Yamashita and Mr. Kosuke Tsuda (Nagoya University) for their contribution to the project. We also thank Dr. Masaharu Kubota (National Agriculture and Food Research Organization, Japan) for providing *Pythium aphanidermatum* strain, Prof. Takashi Tsuge (Chubu University, Japan) for providing *Fusarium oxysporum* strain and Prof. Yoshitaka Takano (Kyoto University, Japan) for providing *Colletotrichum orbiculare* strain. This work was supported by a Grant-in-Aid for Scientific Research (B) (17H03771, 20H02985, 23H02212) to DT and (17H03963) to KK from the Japan Society for the Promotion of Science (JSPS), and by Japan Science and Technology Agency (JST), PRESTO (JPMJPR22D2) to HK.

## SUPPLEMENTAL DATA

**Supplemental Figure S1.** Purification procedure of *Phytophthora infestans* ceramide (Pi-Cer) and ceramide phosphoethanolamine (Pi-CerPE) elicitors.

**Supplemental Figure S2.** NMR spectra of Pi-Cer A (CDCl_3_, 400 MHz for ^1^H, 100 MHz for ^13^C).

**Supplemental Figure S3.** NMR spectra of Pi-Cer B (CDCl_3_, 600 MHz for ^1^H, 150 MHz for ^13^C).

**Supplemental Figure S4.** NMR spectra of Pi-Cer C (CDCl_3_, 400 MHz for ^1^H, 100 MHz for ^13^C).

**Supplemental Figure S5.** NMR spectra of Pi-Cer D (CDCl_3_, 600 MHz for ^1^H, 150 MHz for ^13^C).

**Supplemental Figure S6.** NMR spectra of Pi-CerPE A (CDCl_3_-CD_3_OD 4:1, 400 MHz for ^1^H, 100 MHz for ^13^C).

**Supplemental Figure S7.** NMR spectra of Pi-CerPE B (CDCl_3_-CD_3_OD 4:1, 600 MHz for ^1^H, 150 MHz for ^13^C).

**Supplemental Figure S8.** NMR spectra of Pi-CerPE D (CDCl_3_-CD_3_OD 4:1, 600 MHz for ^1^H, 150 MHz for ^13^C).

**Supplemental Figure S9.** NMR spectra of Pi-CerPE D (CDCl_3_-CD_3_OD 4:1, 600 MHz for ^1^H, 150 MHz for ^13^C).

**Supplemental Figure S10.** MS/MS fragmentation of Pi-Cer A (precursor ion: [M + Na]^+^).

**Supplemental Figure S11.** Two-dimensional NMR correlations of Pi-Cer B (thick bonds: DQF-COSY, curved arrows: HMBC, l + m =15).

**Supplemental Figure S12.** MS/MS fragmentation of Pi-Cer B (precursor ion: [M + Na]^+^).

**Supplemental Figure S13.** Two-dimensional NMR correlations of Pi-Cer B (thick bonds: DQF-COSY, curved arrows: HMBC, l + m + n = 19).

**Supplemental Figure S14.** DQF-COSY (thick bonds), HMBC (curved arrows), and NOESY (dashed curves) correlations of Pi-Cer D.

**Supplemental Figure S15.** Linked-scan FAB MS/MS (negative ion mode) of the fatty acid derived from Pi-Cer D by acid hydrolysis.

**Supplemental Figure S16.** DQF-COSY (thick bonds) and HMBC (curved arrows) correlations of Pi-CerPE D.

**Supplemental Figure S17.** Ceramide phosphoethanolamine (Pi-CerPE) elicitors purified from methanol extract of *P. infestans* mycelia (Pi-MEM), which can induce the production of reactive oxygen species (ROS) in potato suspension-cultured cells.

**Supplemental Figure S18.** Activation of *A. thaliana WRKY33* promoter by Pi-Cer and Pi-CerPE elicitors.

**Supplemental Figure S19.** Pi-Cer contents in mycelia of oomycete and fungal plant pathogens.

**Supplemental Figure S20.** Induction of reactive oxygen species (ROS) production by Pi-Cer D and Pi-CerPE D treatment in potato suspension-culture cells.

**Supplemental Figure S21.** Pi-Cer D, but not Pi-Cer C, can induce the production of reactive oxygen species (ROS) in *Nicotiana benthamiana*.

**Supplemental Figure S22.** Production of phytoalexins in potato tuber is not induced by Pi-Cers and Pi-CerPEs.

**Supplemental Figure S23.** Purification procedure of *Phytophthora infestans* diacylglycerol (Pi-DAG) elicitors.

**Supplemental Figure S24.** ESI-TOF MS of Pi-DAG A. NMR spectra of Pi-DAG A (CDCl_3_, 400 MHz for ^1^H and 100 MHz for ^13^C). Two-dimensional NMR of Pi-DAG A (CDCl_3_, 400 MHz). Determination of fatty acids of Pi-DAG A by MS/MS.

**Supplemental Figure S25.** ^1^H NMR spectrum of Pi-DAG B (CDCl_3_, 400 MHz). Determination of fatty acids in Pi-DAG B by negative ion FAB MS

**Supplemental Figure S26.** NMR spectra of Pi-DAG C (CDCl_3_, 400 MHz for ^1^H, 100 MHz for ^13^C). Determination of fatty acids in Pi-DAG C by positive ion ESI MS/MS. Confirmation of fatty acid linkage in Pi-DAG C.

**Supplemental Figure S27.** NMR spectra of Pi-DAG D (MeOD, 400 MHz for ^1^H, 100 MHz for ^13^C). Determination of fatty acids in Pi-DAG D by hydrolysis followed by negative ion FAB MS and MS/MS analysis of two fatty acid products.

**Supplemental Figure S28.** Pi-DAG contents in mycelia of oomycete plant pathogens.

**Supplemental Figure S29.** Eicosapentaenoic acid (EPA) does not induce the production of reactive oxygen species (ROS) in potato leaves.

**Supplemental Figure S30.** Callose deposition of Arabidopsis seedlings treated with EPA or Pi-Cer D.

**Supplemental Figure S31.** Expression profiles of Arabidopsis genes upregulated by co-treatment of EPA and Pi-Cer D, EPA or Pi-Cer D.

**Supplemental Figure S32.** Gene ontology (GO) enrichment analysis of up-regulated genes in Arabidopsis treated with eicosapentaenoic acid (EPA), Pi-Cer D or their mixture for 12 h.

