## Supplemental Figures for "Two structurally different oomycete lipophilic MAMPs induce distinctive plant immune responses"

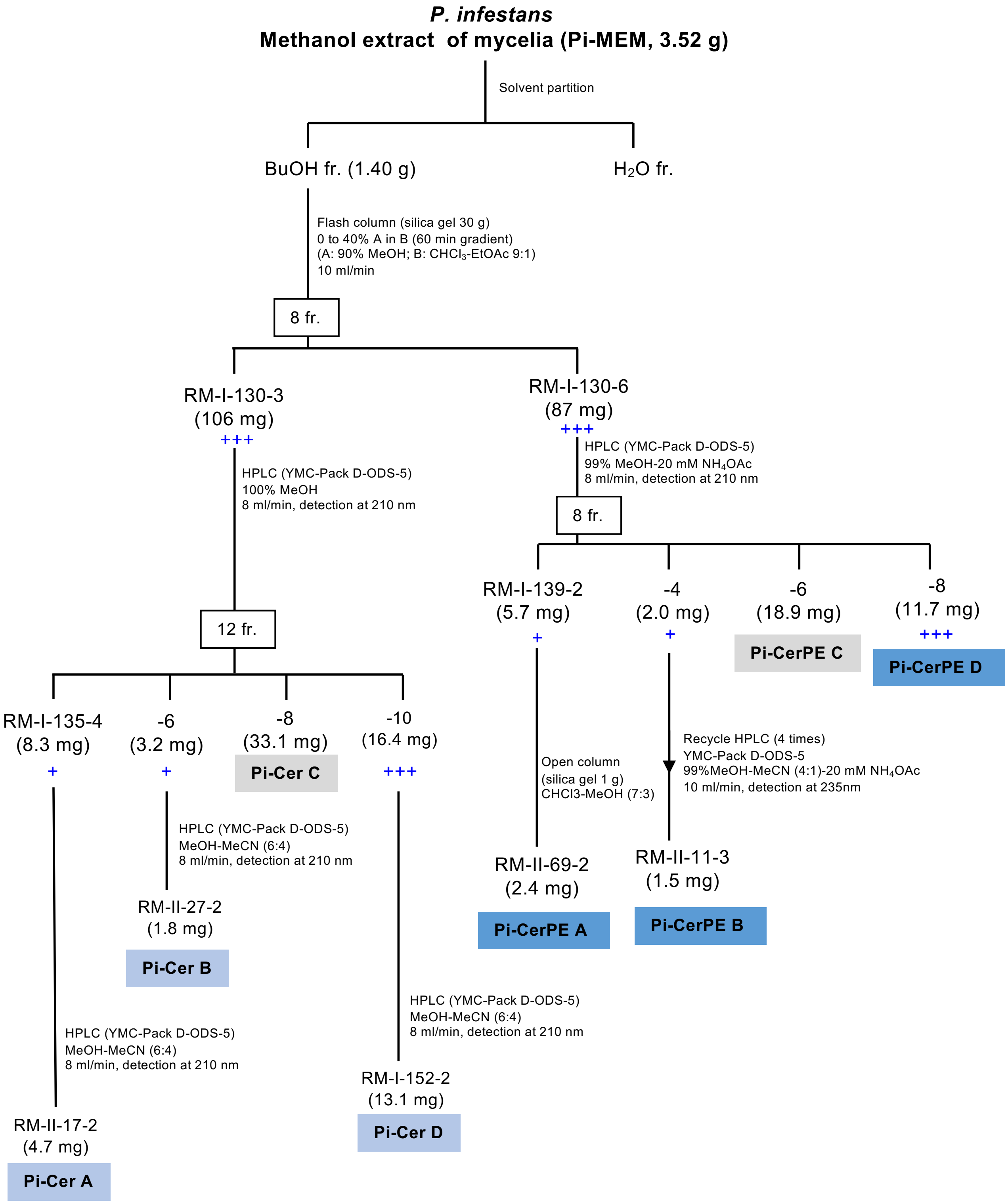


**Supplemental Figure S1.** Purification procedure of *Phytophthora infestans* ceramide (Pi-Cer) and ceramide phosphoethanolamine (Pi-CerPE) elicitors. Methanol extract of *P. infestans* mycelia (Pi-MEM) was dried and fractionated with butanol (BuOH) and water. BuOH-soluble fraction was then partitioned into fractions (fr.) by a series of column chromatography as indicated. Reactive oxygen species (ROS) producing activity (+) was detected in potato suspension-cultured cells as L-012-mediated chemiluminescence after elicitor treatment. See supplemental document for the detail of the procedure.

#### Pi-Cer A

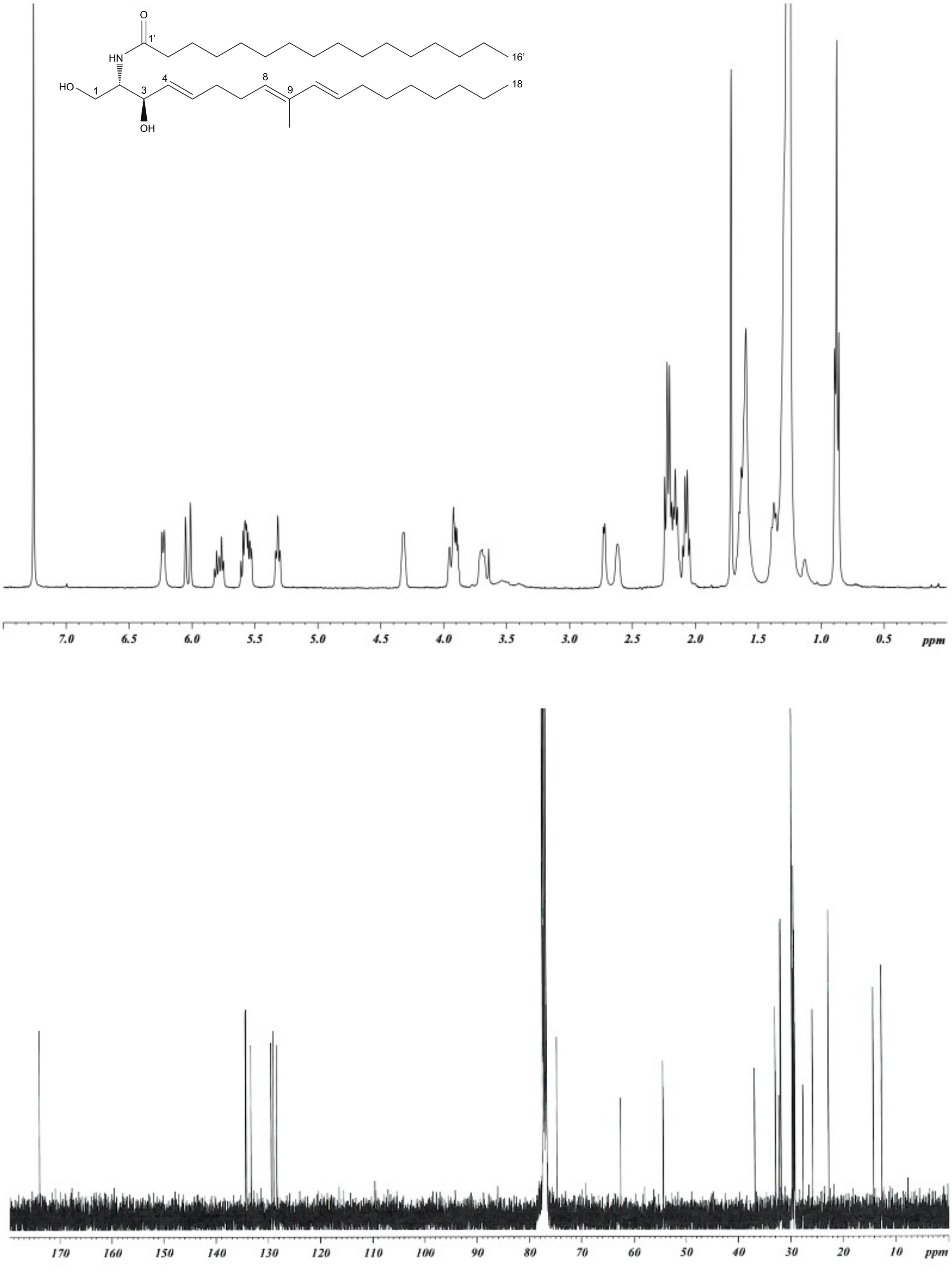


**Supplemental Figure S2.** NMR spectra of Pi-Cer A ($CDCl_3$, 400 MHz for $^1H$, 100 MHz for $^{13}C$).

**Pi-Cer B**

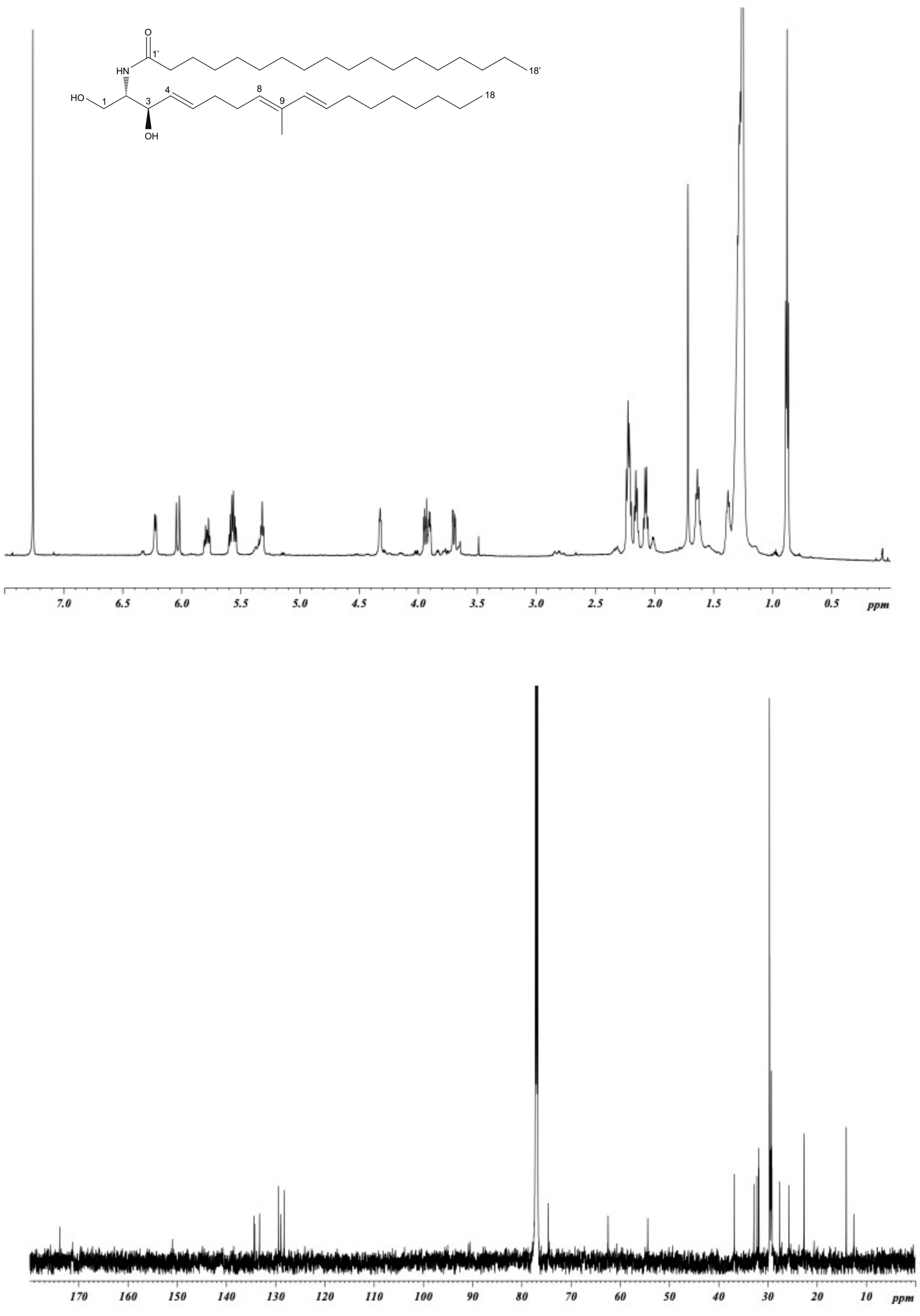


**Supplemental Figure S3.** NMR spectra of Pi-Cer B ($CDCl_3$, 600 MHz for $^{1}H$, 150 MHz for $^{13}C$).

#### Pi-Cer C

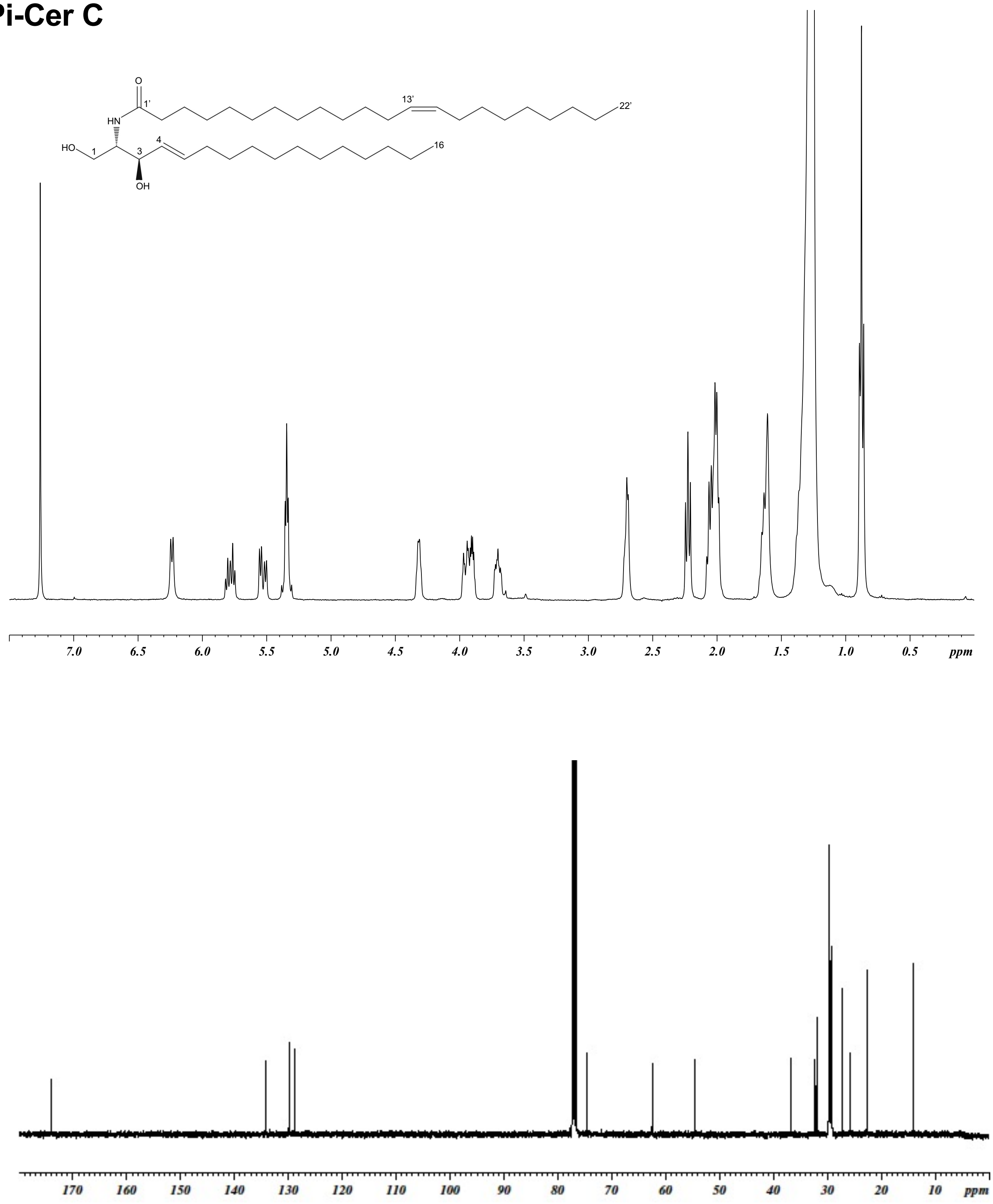


**Supplemental Figure S4.** NMR spectra of Pi-Cer C ($CDCl_3$, 400 MHz for $^1H$, 100 MHz for $^{13}C$).

### Pi-Cer D

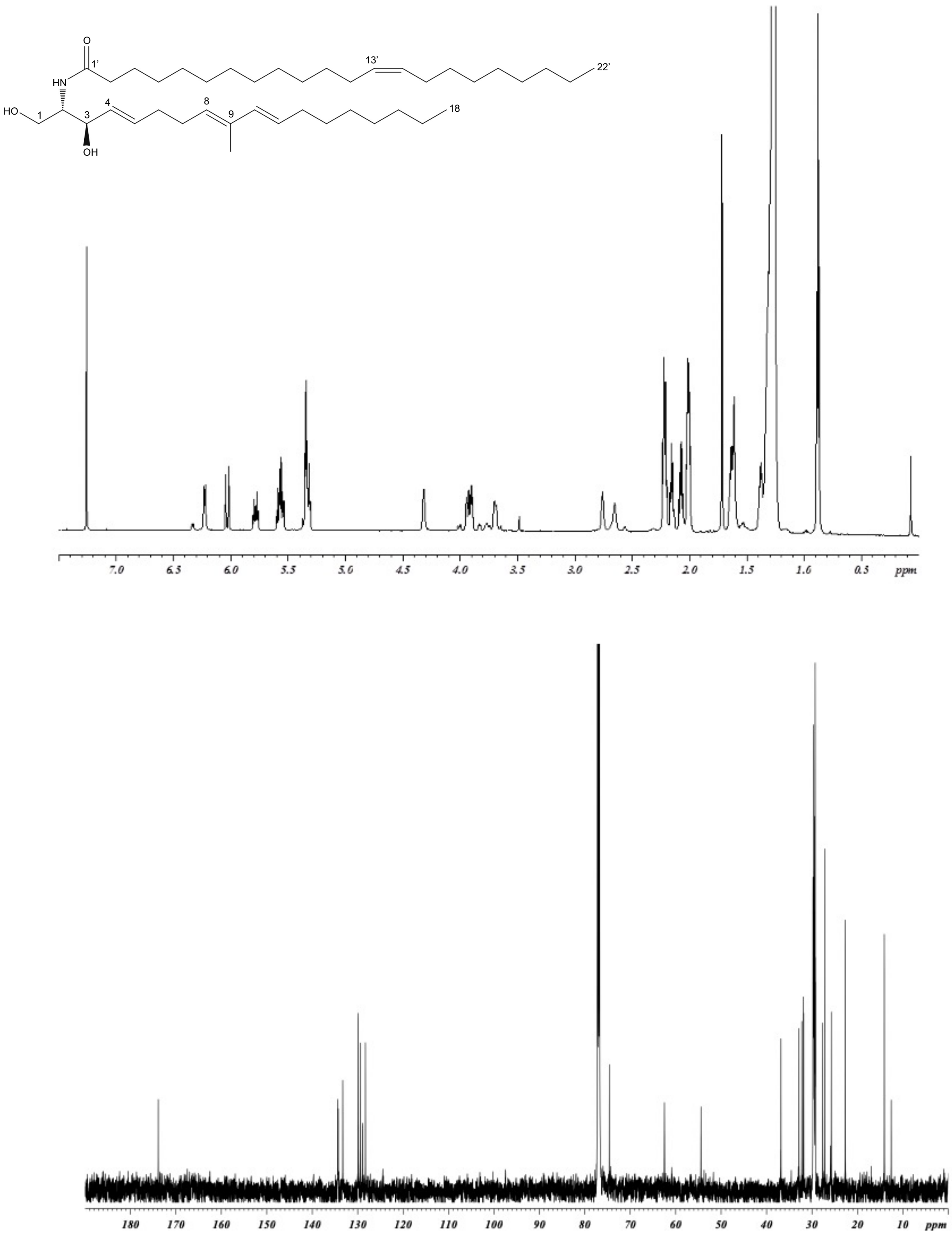


**Supplemental Figure S5.** NMR spectra of Pi-Cer D ($CDCl_3$, 600 MHz for $^1H$, 150 MHz for $^{13}C$).

### Pi-CerPE A

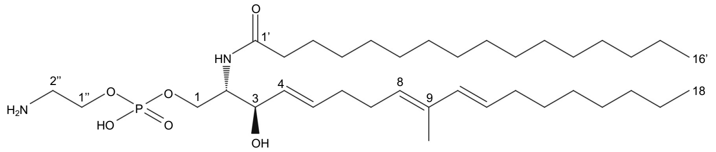


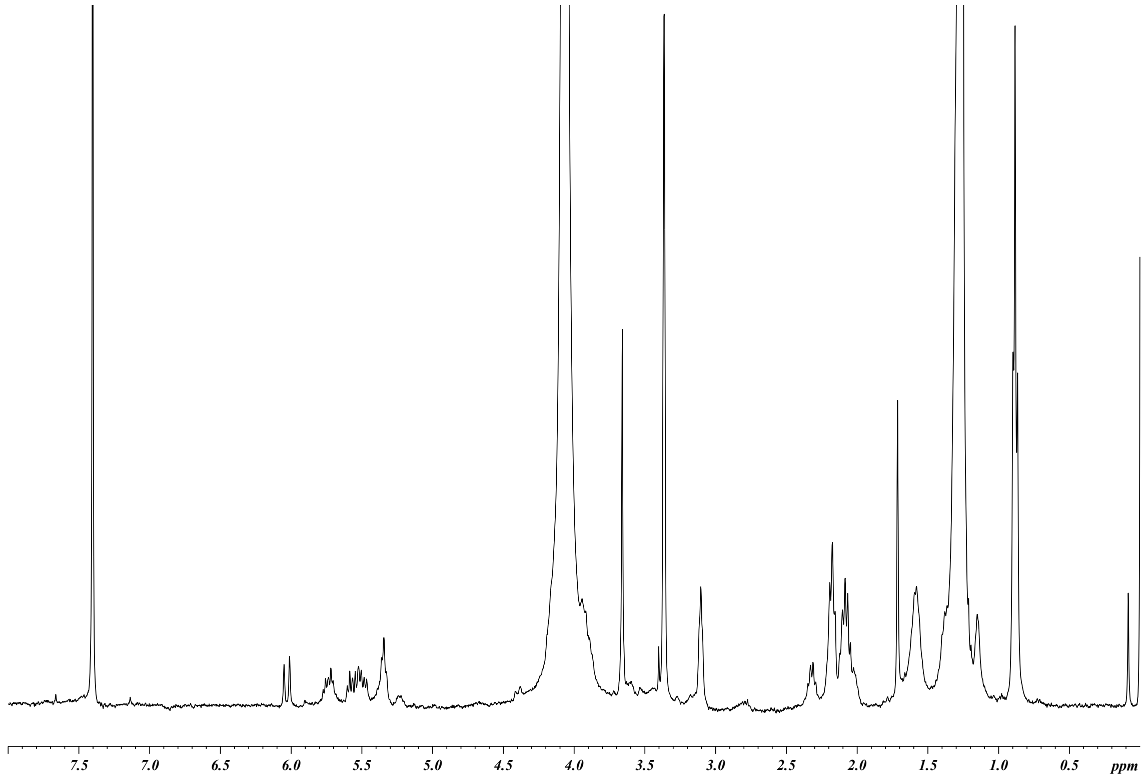


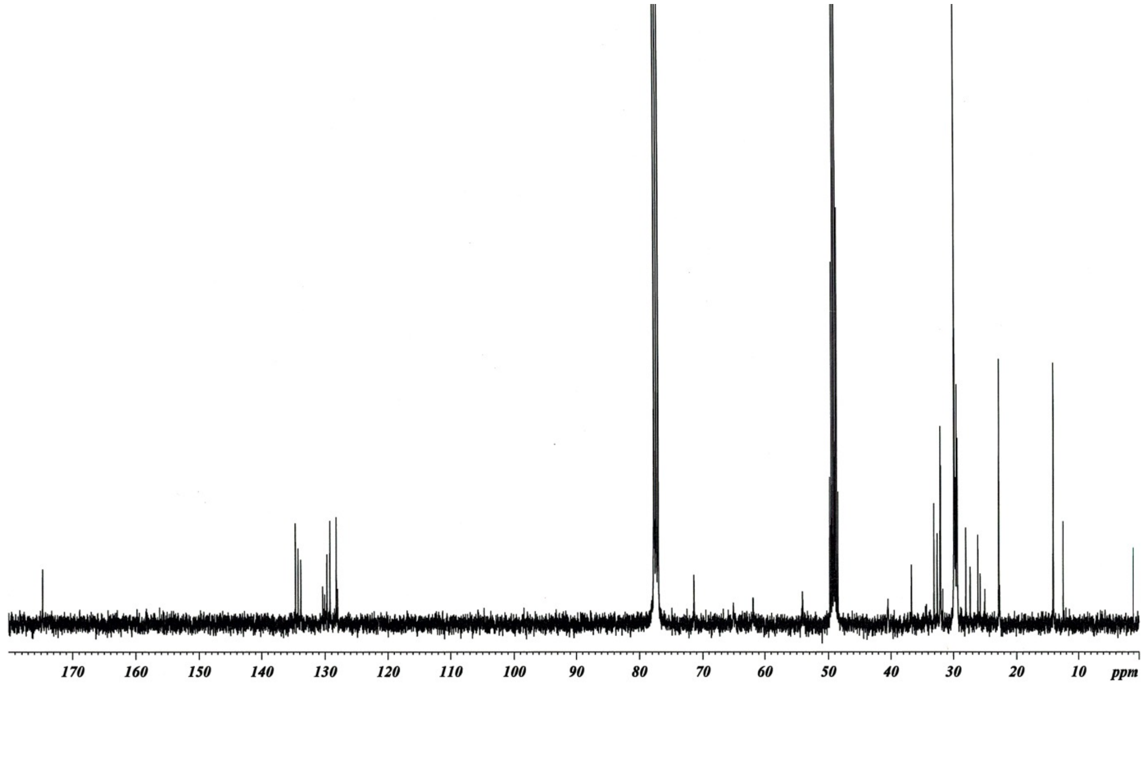


**Supplemental Figure S6.** NMR spectra of Pi-CerPE A ($CDCl_3$-$CD_3OD$ 4:1, 400 MHz for $^1H$, 100 MHz for $^{13}C$).

### Pi-CerPE B

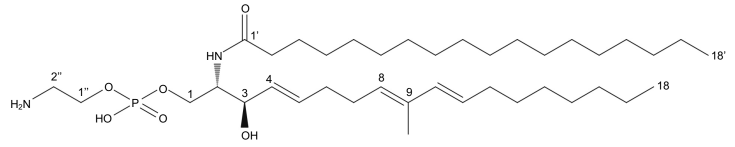


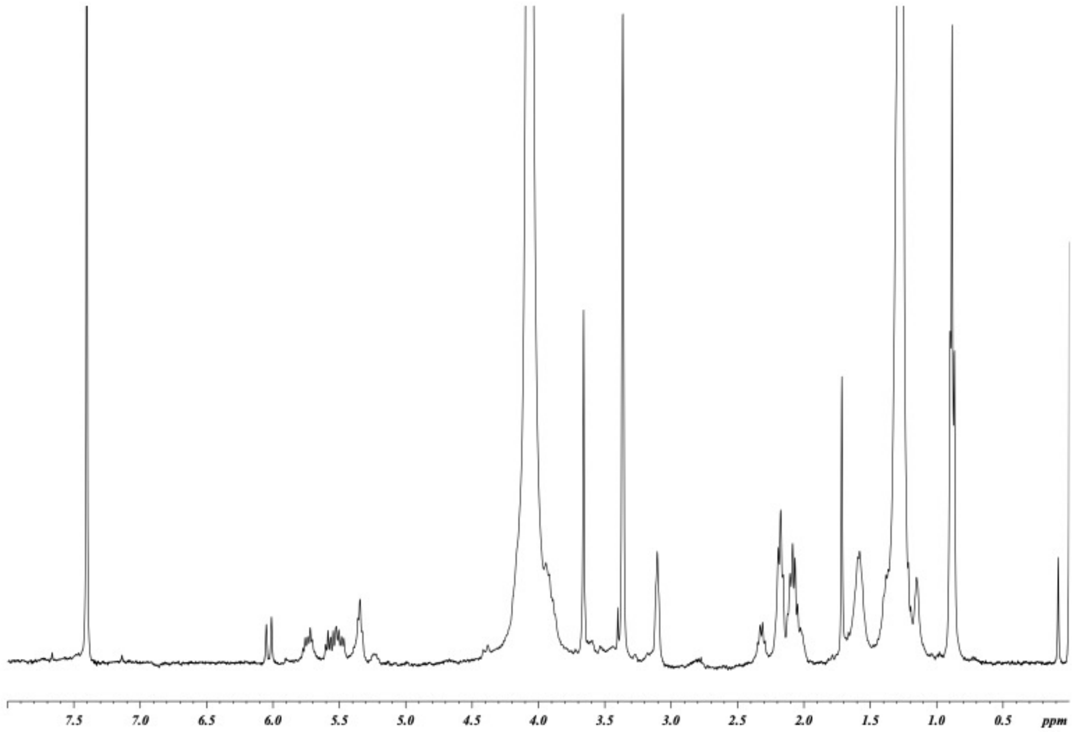


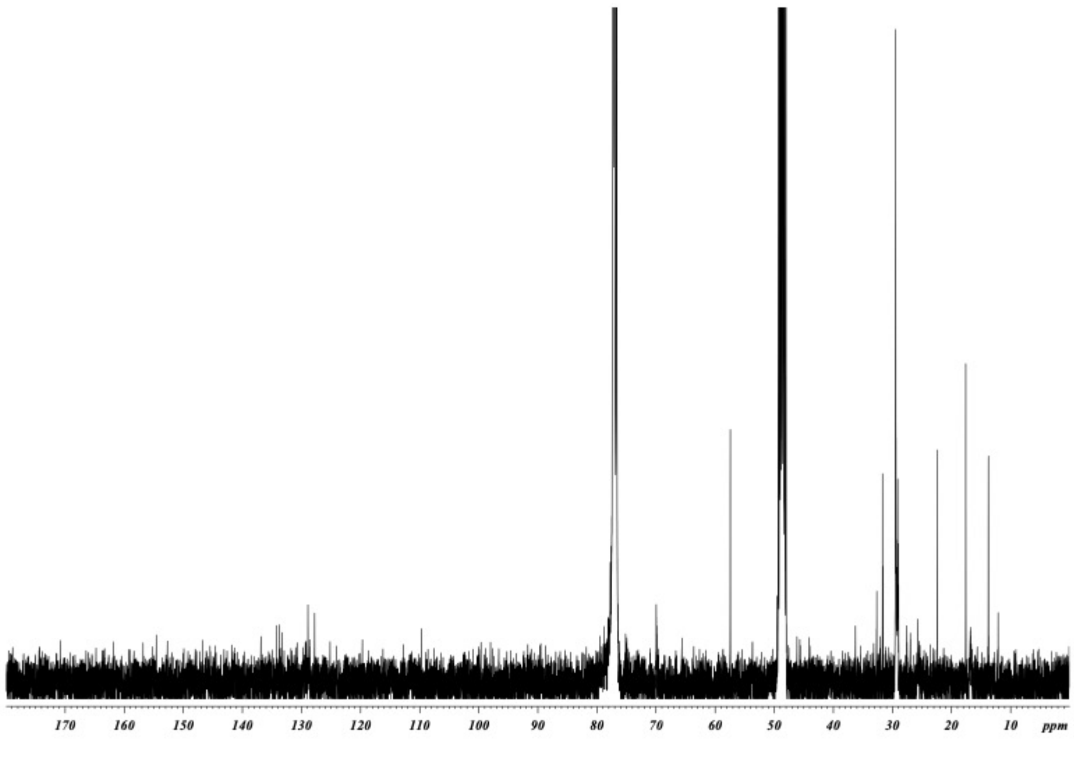


**Supplemental Figure S7.** NMR spectra of Pi-CerPE B ($CDCl_3$-$CD_3OD$ 4:1, 600 MHz for $^1H$, 150 MHz for $^{13}C$).

### Pi-CerPE C

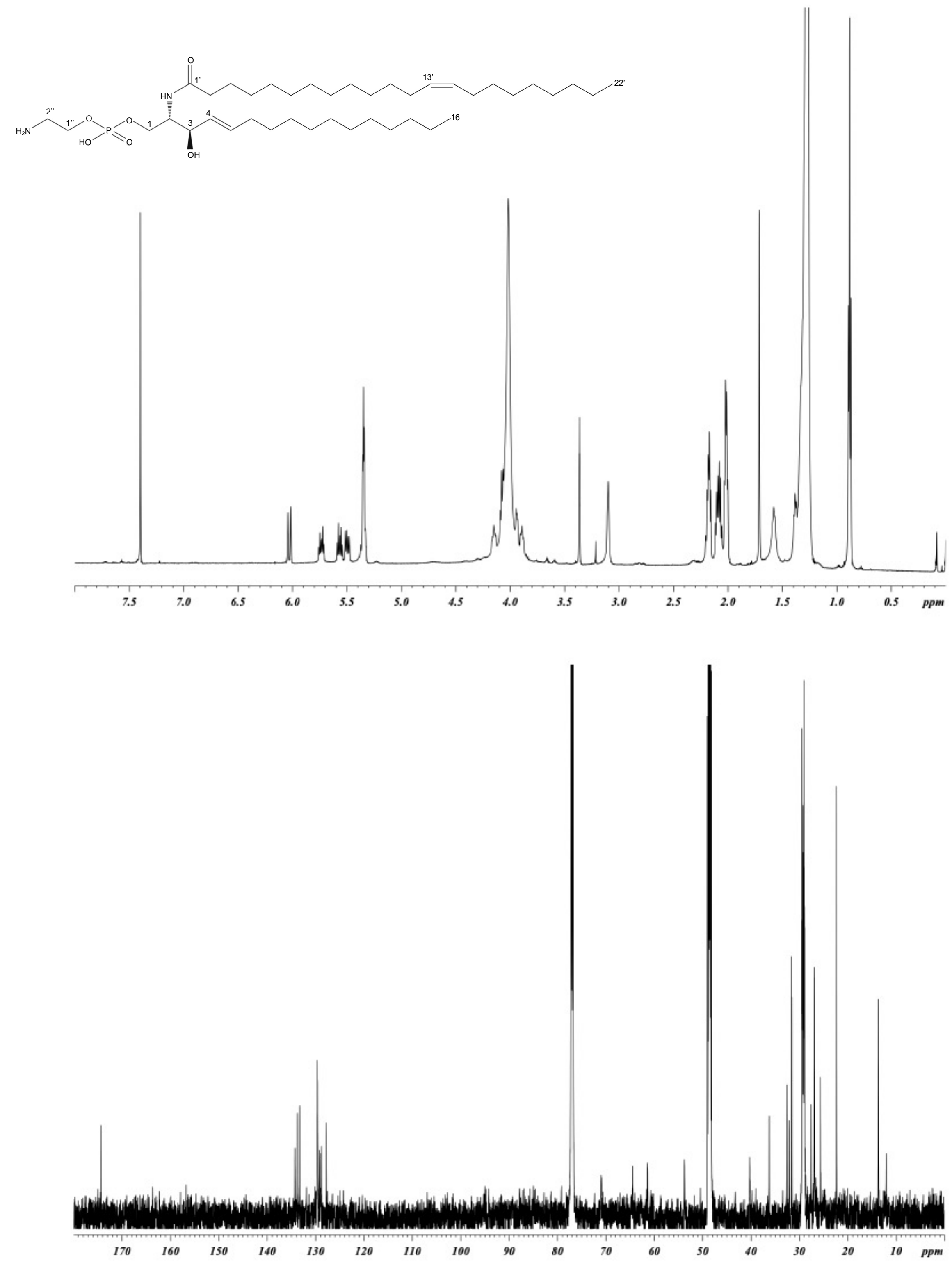


**Supplemental Figure S8.** NMR spectra of Pi-CerPE D ($CDCl_3$-$CD_3OD$ 4:1, 600 MHz for $^1H$, 150 MHz for $^{13}C$).

### Pi-CerPE D

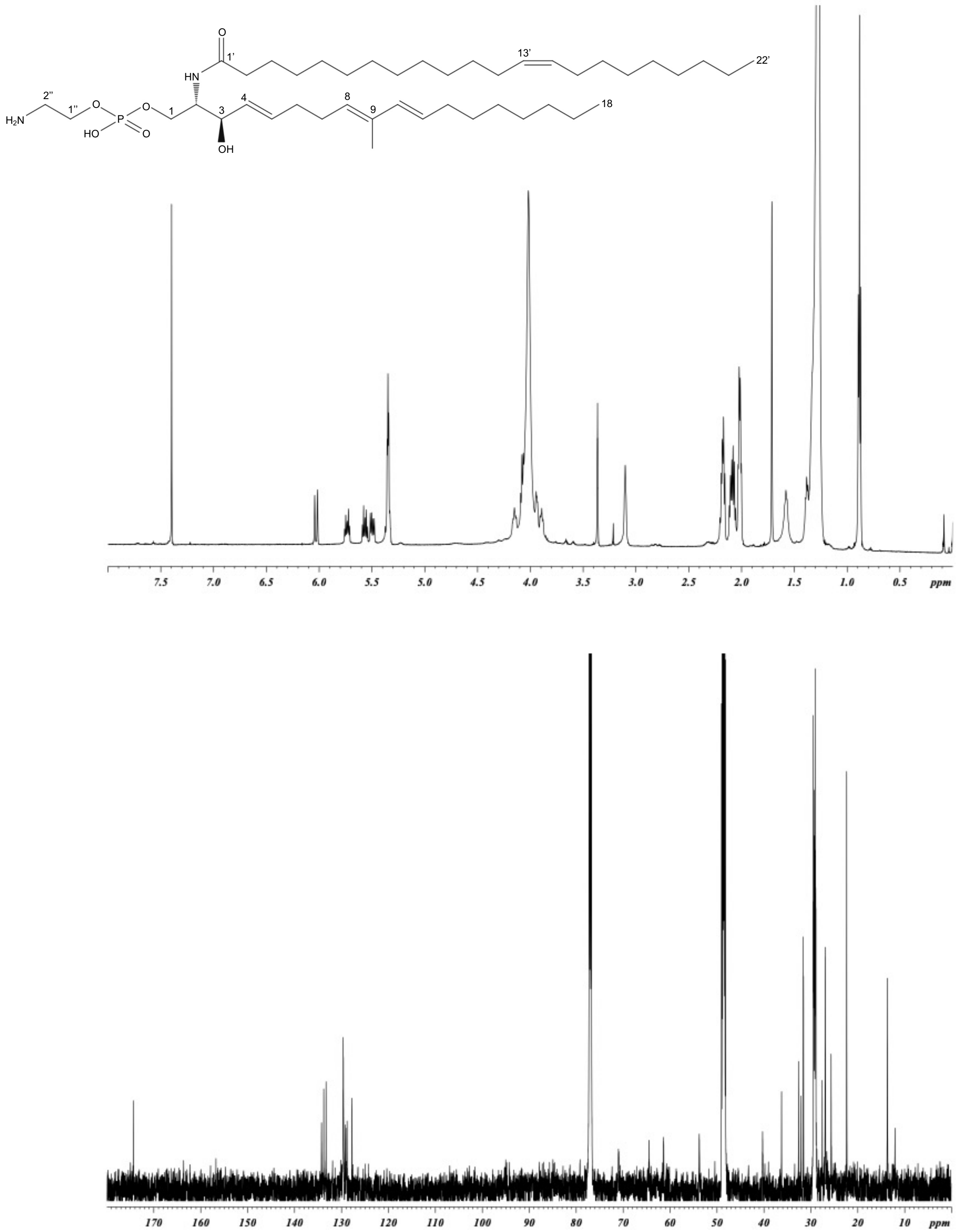


**Supplemental Figure S9.** NMR spectra of Pi-CerPE D ($CDCl_3$-$CD_3OD$ 4:1, 600 MHz for $^1H$, 150 MHz for $^{13}C$).

**Supplemental Figures S10-16 and Supplemental document**

### Contents

**General procedure**

**Purification of ceramide elicitors**

**Spectral data**

Pi-Cer A, B, C and D

Pi-CerPE A, B, C and D

**Structural analysis**

Pi-Cer A, B, C and D

Pi-CerPE A, B, C and D

**References**

#### General procedure

Specific rotations were recorded on a DIP-370 spectrometer (Jasco, Tokyo, Japan). IR spectra were measured using an FT-IR-7000S spectrometer (Jasco). UV spectra were obtained using a V-530 spectrometer (Jasco). High-resolution ESI-TOF mass spectra (MS) were recorded on a Mariner Biospectrometry Workstation (Applied Biosystems, CA, USA) in the positive ESI mode using 50–80% MeOH–0.1% HCOOH as the infusion solvent. FAB MS/MS was performed on an Mstation JMS-700 mass spectrometer (Jeol, Tokyo, Japan) in the linked-scan mode using a *m*-nitrobenzyl alcohol–thioglycerol mixture as a matrix. NMR spectra were recorded on an AVANCE ARX400 (400 MHz for $^{1}$H) or AMX2 600 (600 MHz for $^{1}$H) spectrometer (Bruker Biospin, Yokohama, Japan). The chemical shifts (ppm) were referenced to the TMS peak. Flash column chromatography with a packed column was performed on an 880-PU pump (Jasco) equipped with an 880-02 gradient unit (Jasco). Analytical HPLC was performed on a high-pressure gradient system (Jasco) composed of a PU-1580 pump, a DG-2080-54 degasser, and an MD-2010 plus photodiode array detector. Preparative HPLC was performed with a high-pressure gradient system (Jasco) composed of a PU-1586 pump, DG-1580-53 degasser, and UV 1570 detector.

#### Purification of ceramide elicitors

The mycelia obtained from *P. infestans* culture broths (37 flasks, 3.7 L in total) were homogenized in MeOH (10 mL/1g mycelia), and the insoluble materials were removed by centrifugation (3000 x g, 30 min). The supernatant was concentrated under reduced pressure to give a crude elicitor (3.52 g). The crude elicitor was suspended in water (300 mL) and extracted twice with butanol (300 mL). The butanol layers were combined and concentrated under reduced pressure to give an oily material (1.40 g), which was subjected to flash column chromatography [Hi-Flash™ column, size L, silica gel 30 g (Yamazen Co., Osaka, Japan), 0 to 40% A solvent in B solvent (60 min linear gradient), A=90% MeOH, B=$CHCl_3$-EtOAc (9:1), flow 10 mL/min]. The third fraction (106 mg) eluted with 9-12% A solvent was separated by HPLC [YMC-Pack D-ODS-5 (20 mm i.d. x 250 mm, YMC Co., Kyoto, Japan), MeOH, flow 8 mL/min, detection at 210 nm] to give crude Pi-Cer A (8.3 mg, retention time ($t_R$) = 24.5 min), crude Pi-Cer B (3.2 mg, $t_R$ = 32.2 min), Pi-Cer C (33.1 mg, $t_R$ = 36.6 min), and crude Pi-Cer D (16.4 mg, $t_R$ = 41.9 min). The crude Pi-Cer A, B, and D were purified by HPLC [YMC-Pack D-ODS-5 (20 mm i.d. x 250 mm), MeOH-MeCN (6:4), flow 8 mL/min, detection at 210 nm] to give Pi-Cer A (4.7 mg, $t_R$ = 46.4 min), Pi-Cer B (1.8 mg, $t_R$ = 64.2 min), and Pi-Cer D (13.1 mg, $t_R$ = 69.9 min), respectively. The sixth fraction (87.0 mg) eluted with 28-31% A solvent in the flash column chromatograph was separated by HPLC [YMC-Pack D-ODS-5 (20 mm i.d. x 250 mm), 99% MeOH-20 mM $NH_4OAc$, flow 8 mL/min, detection at 210 nm] to give crude Pi-CerPE A (5.7 mg, $t_R$ = 28.4 min), crude Pi-CerPE B (2.0 mg, $t_R$ = 37.5 min), Pi-CerPE C (18.9 mg, $t_R$ = 47.0 min), and Pi-CerPE D (11.7 mg, $t_R$ = 52.9 min). The crude Pi-CerPE A was further purified using silica gel (Wakogel C-300, 1 g, Osaka, Japan) column chromatography [$CHCl_3$-MeOH (7:3)] to give Pi-CerPE A (2.4 mg). The crude Pi-CerPE B was purified by recycled HPLC [YMC-Pack D-ODS-5 (20 mm i.d. x 250 mm), 99% MeOH-MeCN (80:20)-20 mM $NH_4OAc$, flow 10 mL/min, detection at 235 nm, four recycling times] to give Pi-CerPE B (1.5 mg, $t_R$ = 40.5 min).

**Spectral data for ceramide elicitors**

***Pi-Cer A***: colorless power, $[\alpha]^{29}_D$ −1.5 (*c* 0.21, $CHCl_3$), $^1$H NMR (400 MHz, $CDCl_3$) δ 0.88 (t, J=6.6 Hz, 6H, H-18, 16'), 1.26-1.40 (m, 32H, H-14 to 17, H-4' to 15'), 1.38 (m, 2H, H-13), 1.63 (m, 2H, H-3'), 1.72 (s, 3H, 9-Me), 2.08 (m, 2H, H-12), 2.16 (m, 2H, H-6), 2.21 (m, 2H, H-7), 2.22 (m, 2H, H-2'), 2.59 (brs, 1-OH), 2.71 (brd, J=4.8 Hz, 3-OH), 3.70 (m, 1H, H-1), 3.90 (m, 1H, H-2), 3.94 (m, 1H, H-1), 4.32 (m, 1H, H-4), 5.32 (t, J=6.6 Hz, 1H, H-8), 5.55 (m, 1H, H-4), 5.58 (m, 1H, H-11), 5.78 (dt, J=15.5, 7.1 Hz, 1H, H-5), 6.03 (d, J=15.6 Hz, 1H, H-10), 6.23 (d, J=7.2 Hz, 1H, NH), $^{13}$C NMR (100 MHz, $CDCl_3$) δ 12.6 (9-Me), 14.1 (2C, C-18, 16’), 22.7 (2C, C-17, 15’), 25.7 (C-3'), 27.6 (C-7), 29.2-29.7 (13C, C-13 to 15, C-4' to 13'), 31.8 (C-14'), 31.9 (C-16), 32.2 (C-6), 32.9 (C-12), 36.8 (C-2'), 54.3 (C-2), 62.5 (C-1), 74.7 (C-3), 128.3 (C-

11), 129.0 (C-8), 129.4 (C-4), 133.3 (C-5), 134.3 (C-9), 134.4 (C-10), 173.9 (C-1'), high-resolution (HR) MS (ESI-TOF) *m/z* 548.5051 $[M+H]^+$, calcd for $C_{35}H_{66}NO_3$, 548.5037.

***Pi-Cer B***: colorless power, $[\alpha]^{25}_D$ –3.2 (*c* 0.15, $CHCl_3$), UV (MeOH) $\lambda_{max}$ 235 nm ($\varepsilon$ 18,000); IR (film) 3299, 1642, 1550, 1260, 1076, 962, 759 $cm^{-1}$; $^1$H NMR (600 MHz, $CDCl_3$) δ 0.88 (t, J=7.0 Hz, 3H, H-18), 0.88 (t, J=7.0 Hz, 3H, H-18'), 1.26-1.40 (m, 36H, H-14 to 17, H-4' to 17'), 1.38 (m, 2H, H-13), 1.64 (m, 2H, H-3'), 1.72 (s, 3H, 9-Me), 2.08 (m, 2H, H-12), 2.15 (m, 2H, H-6), 2.21 (m, 2H, H-7), 2.23 (m, 2H, H-2'), 3.70 (dd, J=11.2, 3.2 Hz, 1H, H-1), 3.90 (m, 1H, H-2), 3.94 (dd, J=11.2, 3.7 Hz, 1H, H-1), 4.32 (m, 1H, H-3), 5.32 (t, J=7.0 Hz, H-8), 5.55 (m, 1H, H-4), 5.58 (m, 1H, H-11), 5.79 (dt, J=15.5, 7.1 Hz, 1H, H-5), 6.03 (d, J=15.5 Hz, 1H, H-10), 6.22 (d, J=7.3 Hz, 1H, NH), $^{13}$C NMR (150 MHz, $CDCl_3$) δ 12.6 (9-Me), 14.1 (2C, C-18, 18'), 22.7 (2C, C-17, 17'), 25.8 (C-3'), 27.6 (C-7), 29.2-29.7 (15C, C-13 to 15, C-4' to 15'), 31.9 (2C, C-16, 16'), 32.2 (C-6), 32.9 (C-12), 36.8 (C-2'), 54.4 (C-2), 62.5 (C-1), 74.6 (C-3), 128.3 (C-11), 129.0 (C-8), 129.4 (C-4), 133.3 (C-5), 134.3 (C-9), 134.4 (C-10), 173.9 (C-1'), HRMS (ESI-TOF) *m/z* 576.5374 $[M+H]^+$, calcd for $C_{37}H_{70}NO_3$, 576.5350.

***Pi-Cer C***: colorless power, $[\alpha]^{30}_D$ –3.2 (*c* 0.23, $CHCl_3$), $^1$H NMR (600 MHz, $CDCl_3$) δ 0.88 (t, J=7.0 Hz, 3H, H-16), 0.88 (t, J=7.0 Hz, 3H, H-22'), 1.26-1.40 (m, 46H, H-7 to 15, H-4' to 11', H-16' to 21'), 1.64 (quint, J=7.2 Hz, 2H, H-3'), 2.01 (m, 2H, H-12'), 2.01 (m, 2H, H-15'), 2.05 (m, 2H, H-6), 2.22 (t, J=7.2 Hz, 2H, H-2'), 2.76 (m, 1H, 1-OH), 2.76 (m, 1H, 3-OH), 3.70 (m, 1H, H-1), 3.90 (m, 1H, H-2), 3.96 (brd, J=11.2 Hz, 1H, H-1), 4.31 (m, 1H, H-3), 5.35 (m, 1H, H-13'), 5.35 (m, 1H, H-14'), 5.53 (dd, J=15.2, 6.6 Hz, 1H, H-4), 5.78 (dt, J=15.2, 7.2 Hz, 1H, H-5), 6.24 (d, J=7.6 Hz, 1H, NH), $^{13}$C NMR (150 MHz, $CDCl_3$) δ 14.1 (2C, C-16, 22'), 22.7 (2C, C-15, 21'), 25.7 (C-3'), 27.2 (2C, C-12', 15'), 29.1-29.8 (19C, C-7 to 13, C-4' to 11', C-16' to 19'), 31.9 (2C, C-14, 20'), 32.3 (C-6), 36.8 (C-2'), 54.5 (C-2), 62.5 (C-1), 74.7 (C-3), 128.9 (C-4), 129.9 (2C, C-13', 14'), 134.3 (C-5), 173.9 (C-1'), HRMS (ESI-TOF) *m/z* 592.5670 $[M+H]^+$, calcd for $C_{38}H_{74}NO_3$, 592.5663.

***Pi-Cer D***: colorless powder, $[\alpha]^{29}_D$ -1.7 (*c* 0.23, $CHCl_3$), UV (MeOH) $\lambda_{max}$ 235 nm (ε 27000), IR (film) 3303, 1644, 1548, 1284, 1046, 962, 721 $cm^{-1}$, $^1$H NMR (600 MHz, $CDCl_3$) δ 0.88 (t, J=6.9 Hz, 6H, H-18, 22'), 1.26-1.40 (m, 38H, H-13 to 17, 4' to 11', 16' to 21'), 1.61 (m, 2H, H-3'), 1.72 (s, 3H, 9-Me), 2.01 (m, 4H, H-12', 15'), 2.07 (m, 2H, H-12), 2.15 (m, 2H, H-6), 2.21 (m, 2H, H-7), 2.22 (m, 2H, H-2'), 2.65 (brs, 1-OH), 2.76 (brs, 3-OH), 3.69 (brd, J=11.2 Hz, 1H, H-1), 3.90 (m, 1H, H-2), 3.94 (br d, J=11.2 Hz, 1H, H-1), 4.32 (brs, 1H, H-3), 5.32 (t, J=7.1 Hz,

1H, H-8), 5.35 (m, 2H, H-13', 14'), 5.55 (m, 1H, H-4), 5.57 (m, 1H, H-11), 5.78 (dt, J=15.5, 7.1 Hz, 1H, H-5), 6.03 (d, J=15.6 HZ, 1H, H-10), 6.23 (d, J=7.4 Hz, 1H, NH). $^{13}$C NMR (150 MHz, $CDCl_3$) δ 12.6 (9-Me), 14.1 (2C, C-18, 22'), 22.7 (2C, C-17, 21'), 25.7 (C-3'), 27.2 (2C, C-12', 15'), 27.7 (C-7), 29.2-29.8 (15C, C-13 to 15, 4' to 11', 16' to 19'), 31.9 (2C, C-16, 20'), 32.2 (C-6), 32.9 (C-12), 36.8 (C-2'), 54.4 (C-2), 62.5 (C-1), 74.6 (C-3), 128.3 (C-11), 128.9 (C-8), 129.4 (C-4), 129.9 (C-13', 14'), 133.3 (C-5), 134.3 (C-9), 134.4 (C-10), 173.9 (C-1'). HR ESI-TOF-MS *m/z* 630.5845 $[M+H]^+$, calcd for $C_{41}H_{76}NO_3$ 630.5820.

***Pi-CerPE A***: colorless powder, $[\alpha]^{30}_D$ +7.8 (*c* 0.16, $CHCl_3$-MeOH (4:1)), $^1$H NMR (400 MHz, $CDCl_3$-$CD_3OD$ (4:1)) δ 0.88 (t, J=6.4 Hz, 6H, H-18, 16'), 1.20-1.35 (m, 32H, H-14 to 17, H-4' to 15'), 1.38 (m, 2H, H-13), 1.58 (m, 2H, H-3'), 1.71 (s, 3H, 9-Me), 2.08 (m, 4H, H-6, 12), 2.17 (m, 4H, H-7, 2'), 3.10 (brs, 2H, H-2"), 3.88 (m, 1H, H-1), 3.94 (m, 1H, H-2), 4.05 (m, 2H, H-1"), 4.07 (m, 1H, H-3), 4.15 (m, 1H, H-1), 5.34 (m, 1H, H-8), 5.49 (dd, J=15.2, 7.2 Hz, H-4), 5.57 (dt, J=15.6, 7.2 Hz, 1H, H-11), 5.73 (dt, J=15.2, 6.2 Hz, 1H, H-5), 6.03 (d, J=15.6 Hz, 1H, H-10), $^{13}$C NMR (100 MHz, $CDCl_3$-$CD_3OD$ (4:1)) δ 12.5 (9-Me), 14.2 (2C, C-16, 18'), 22.8 (2C, C-15, 17'), 26.1 (C-3'), 28.0 (C-7), 29.4-30.0 (13C, C-13, C-4' to 15'), 32.1 (2C, C-14, 16'), 32.5 (C-6), 33.1 (C-12), 36.7 (C-2'), 40.5 (C-2"), 54.1 (C-2), 61.9 (C-1"), 65.1 (C-1), 71.4 (C-3), 128.2 (C-11), 129.2 (C-8), 129.7 (C-4), 133.8 (C-5), 134.2 (C-9), 134.6 (C-10), 174.7 (C-1'), HRMS (ESI-TOF) *m/z* 671.5109 $[M+H]^+$, $C_{37}H_{72}N_2O_6P$ 671.5123.

***Pi-CerPE B***: colorless powder, $[\alpha]^{30}_D$ +3.3 (*c* 0.075, $CHCl_3$-MeOH (4:1)), UV (MeOH) $\lambda_{max}$ 235 nm ($\varepsilon$ 14,000); IR (film) 3276, 1645, 1550, 1225, 1079, 962, 840, 721 $cm^{-1}$, $^1$H NMR (600 MHz, $CDCl_3$-$CD_3OD$ (4:1)) δ 0.88 (t, J=6.4 Hz, 6H, H-18, 18'), 1.20-1.35 (m, 36H, H-14 to 17, H-4' to 17'), 1.38 (m, 2H, H-13), 1.58 (m, 2H, H-3'), 1.71 (s, 3H, 9-Me), 2.08 (m, 4H, H-6, 12), 2.17 (m, 4H, H-7, 2'), 3.11 (brs, 2H, H-2"), 3.88 (m, 1H, H-1), 3.94 (m, 1H, H-2), 4.05 (m, 2H, H-1"), 4.08 (m, 1H, H-3), 4.15 (m, 1H, H-1), 5.35 (m, 1H, H-8), 5.50 (dd, J=7.4, 15.4 Hz, 1H, H-4), 5.57 (m, 1H, H-11), 5.73 (m, 1H, H-5), 6.03 (d, J=15.6 Hz, 1H, H-10), $^{13}$C NMR (150 MHz, $CDCl_3$-$CD_3OD$ (4:1)) δ 12.6 (9-Me), 14.2 (2C, C-18, 18'), 22.9 (2C, C-17, 17'), 26.1 (C-3'), 28.0 (C-7), 29.4-30.0 (15C, C-13 to 15, C-4' to 15'), 32.0 (C-16'), 32.1 (C-16), 32.6 (C-6), 33.1 (C-12), 36.7 (C-2'), 40.7 (C-2"), 54.1 (C-2), 61.8 (C-1"), 65.0 (C-1), 71.4 (C-3), 128.2 (C-11), 129.3 (C-8), 129.7 (C-4), 133.8 (C-5), 134.3 (C-9), 134.7 (C-10), 174.8 (C-1'), HRMS (ESI-TOF) *m/z* 699.5422 $[M+H]^+$, $C_{39}H_{76}N_2O_6P$ 699.5436.

***Pi-CerPE C***: colorless powder, $[\alpha]^{30}_{D}$ +8.6 (*c* 0.26, $CHCl_3$-MeOH (4:1)), $^1H$ NMR (600 MHz, $CDCl_3$-$CD_3OD$ (4:1)) δ 0.88 (t, J=7.0 Hz, 6H, H-16, 22'), 1.20-1.35 (m, 46H, H-7 to 15, H-4' to 11', H-16' to 21'), 1.58 (m, 2H, H-3'), 2.02 (m, 6H, H-6, 12', 15'), 2.17 (m, 2H, H-2'), 3.11 (brs, 2H, H-2''), 3.89 (m, 1H, H-1), 3.94 (m, 1H, H-2), 4.05 (m, 2H, H-1''), 4.08 (m, 1H, H-3), 4.15 (m, 1H, H-1), 5.35 (m, 2H, H-13', 14'), 5.44 (dd, J=15.6, 7.8 Hz, 1H, H-4), 5.71 (dt, J=15.6, 6.6 Hz, 1H, H-5), $^{13}C$ NMR (150 MHz, $CDCl_3$-$CD_3OD$ (4:1)) δ 14.2 (2C, C-16, 22'), 22.9 (2C, C-15, 21'), 26.2 (C-3'), 27.4 (2C, C-12', 15'), 29.4-30.0 (20C, C-6 to 13, C-4' to 11', C-16' to 19'), 32.1 (2C, C-14, 20'), 32.6 (C-6), 36.7 (C-2'), 40.9 (C-2''), 54.3 (C-2), 62.2 (C-1''), 64.9 (C-1), 71.5 (C-3), 129.2 (C-4), 130.1 (2C, C-13', 14'), 134.7 (C-5), 174.8 (C-1'), HRMS (ESI-TOF) *m/z* 715.5730 $[M+H]^+$, $C_{40}H_{80}N_2O_6P$ 715.5749.

***Pi-CerPE D***: colorless glass, $[\alpha]^{28}_{D}$ +9.2 (*c* 0.14, $CHCl_3$-MeOH (4:1)), UV (MeOH) $\lambda_{max}$ 235 nm (ε 14000), IR (film) 3279, 3006, 1644, 1550, 1222, 1078, 1031, 962, 840, 722 $cm^{-1}$, $^1H$ NMR (600 MHz, $CDCl_3$-$CD_3OD$ (4:1)) δ 0.88 (t, J=6.9 Hz, 6H, H-18, 22'), 1.26-1.40 (m, 38H, H-13 to 17, 4' to 11', 16' to 21'), 1.58 (m, 2H, H-3'), 1.71 (s, 3H, 9-Me), 2.02 (m, 4H, H-12', 15'), 2.07 (m, 2H, H-12), 2.09 (m, 2H, H-6), 2.17 (m, 2H, H-2'), 2.18 (m, 2H, H-7), 3.10 (brs, 2H, H-2''), 3.89 (m, 1H, H-1), 3.94 (m, 1H, H-2), 4.05 (m, 2H, H-1''), 4.08 (m, 1H, H-3), 4.15 (m, 1H, H-1), 5.34 (m, 1H, H-8), 5.35 (m. 2H, H-13', 14'), 5.50 (dd, J=15.3, 7.3 Hz, 1H, H-4), 5.57 (dt, J=15.5, 7.4 Hz, H-11), 5.73 (dt, J=15.3, 7.1 Hz, 1H, H-5), 6.03 (d, J=15.5 Hz, 1H, H-10), $^{13}C$ NMR (150 MHz, $CDCl_3$-$CD_3OD$ (4:1)) δ 12.1 (9-Me), 13.7 (2C, C-18, 22'), 22.4 (2C, C-17, 21'), 25.7 (C-3'), 26.9 (C-12', 15'), 27.6 (C-7), 28.9-29.5 (15C, C-13 to 15, 4' to 11', 16' to 19'), 31.6 (C-16, 20'), 32.1 (C-6), 32.6 (C-12), 36.2 (C-2'), 40.3 (C-2''), 53.8 (C-2), 61.4 (C-1''), 64.5 (C-1), 71.0 (C-3), 127.8 (C-11), 128.8 (C-8), 129.2 (C-4), 129.6 (2C, C-13', 14'), 133.3 (C-5), 133.8 (C-9), 134.3 (C-10), 174.3 (C-1'). HR ESI-TOF-MS *m/z* 753.5932 $[M+H]^+$, calcd for $C_{43}H_{82}N_2O_6P$ 753.5905.

**Structural analysis**

***Pi-Cer A*.** Pi-Cer A was found to be a congener of Pi-Cer B based on the high similarity of $^1H$ NMR (see the following section for the structure analysis of Pi-Cer B). The molecular formulae of Pi-Cer A, $C_{35}H_{65}NO_3$, determined by high-resolution MS is smaller than that of Pi-Cer B by $C_2H_4$, suggesting the difference of the carbon chain length. The carbon chain length was then determined by MS/MS generated from $[M + Na]^+$ ion, indicating that the product ions including the carboxy amide moiety, *m/z* 304.4, 320.5, and 484.7, were smaller than those of Pi-Cer B by $C_2H_4$ (Supplemental Figure S10). The absolute configuration is identical to that of Pi-Cer B by the comparison of those specific rotation. This compound was found to be the known compound.[1]

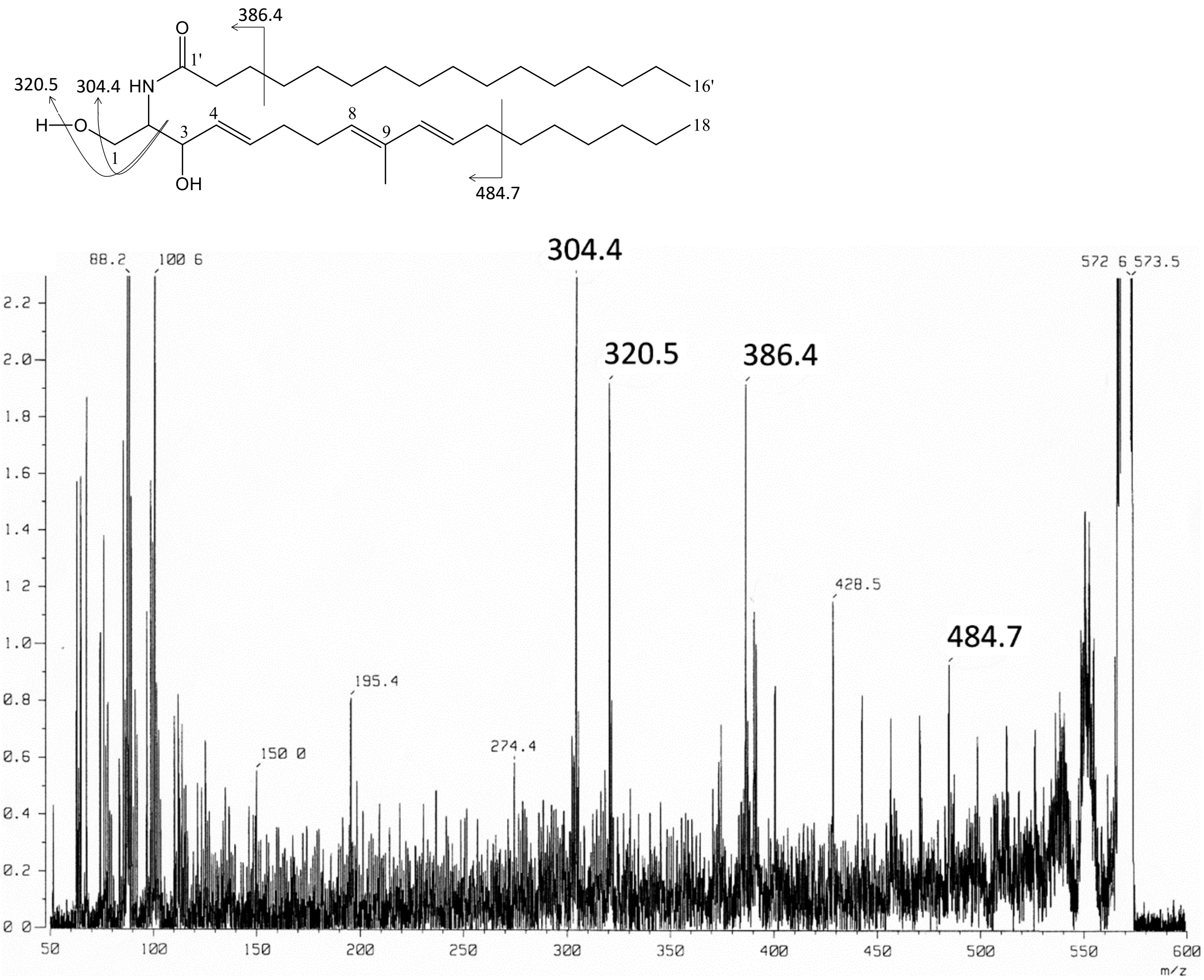


**Supplemental Figure S10.** MS/MS fragmentation of Pi-Cer A (precursor ion: $[M + Na]^+$).

***Pi-Cer B***. The molecular formula $C_{37}H_{69}NO_3$ was determined by HR ESI-TOF-MS. The absorption bands at 3299, 1642, 1550, 1260 $cm^{-1}$ suggested the presence of an amide function. The UV absorption at 235 nm suggested the presence of a conjugated diene. The two-dimensional NMR (HMQC, DQF-COSY) revealed the amide bond and the carbon skeletons except for 15 methylenes (Supplemental Figure S11).

The length of two carbon chains was then determined by MS/MS of [M + Na]+ ion. The product ions at *m/z* 333.7, 349.2, 417.5, and 511.3 containing the fatty acyl moiety indicated the 18-carbon length of the fatty acyl moiety, whereas the product ion at *m/z* 386.2 indicated the presence of a18-carbon sphingosine moiety (Supplemental Figure S12). The geometry at C-4, 8, 10 was determined by the coupling constants and chemical shifts as described in the analysis of Pi-Cer D. The absolute configuration was determined to be 2*S*,3*R* by the same method as that for Pi-Cer D.

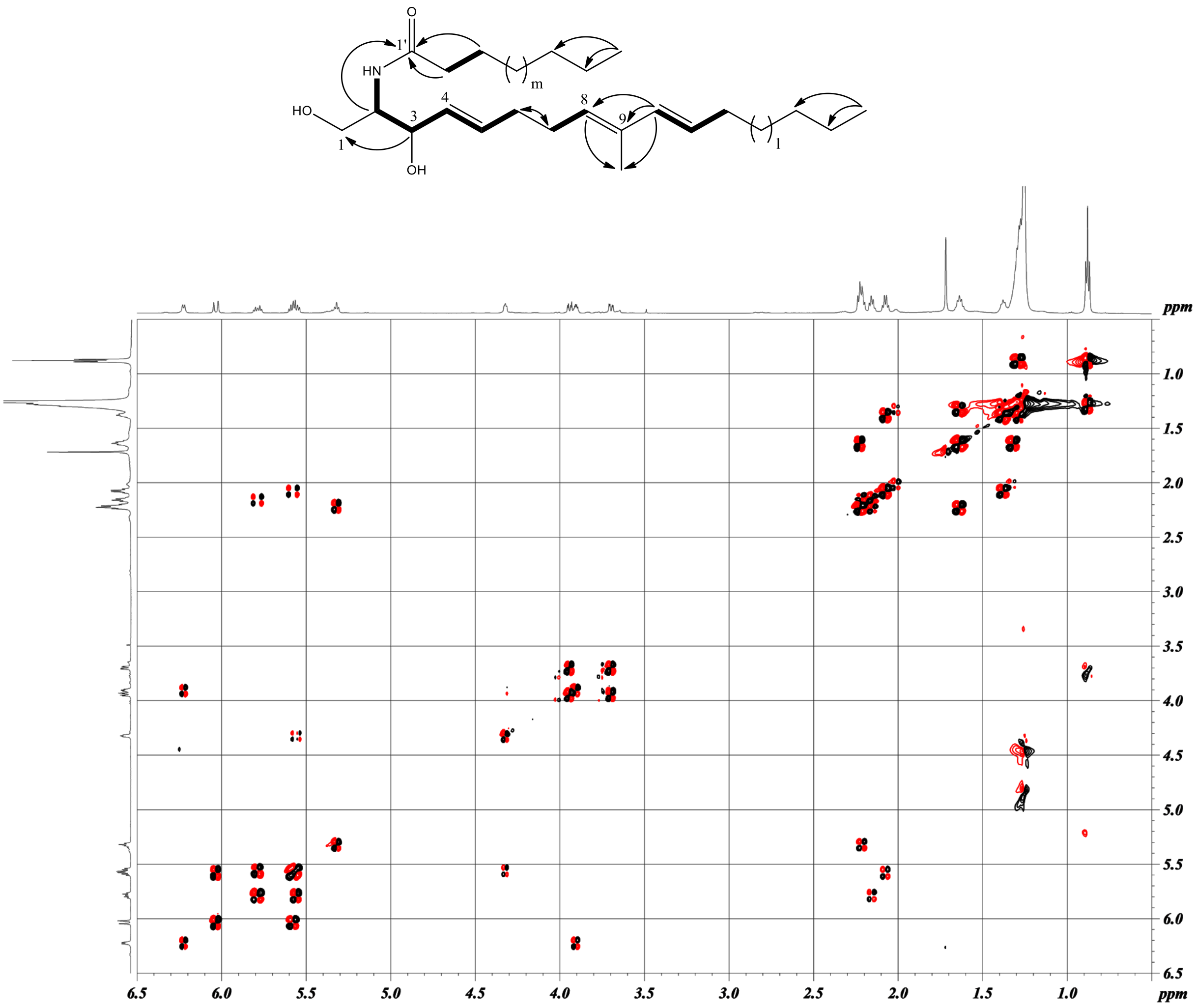

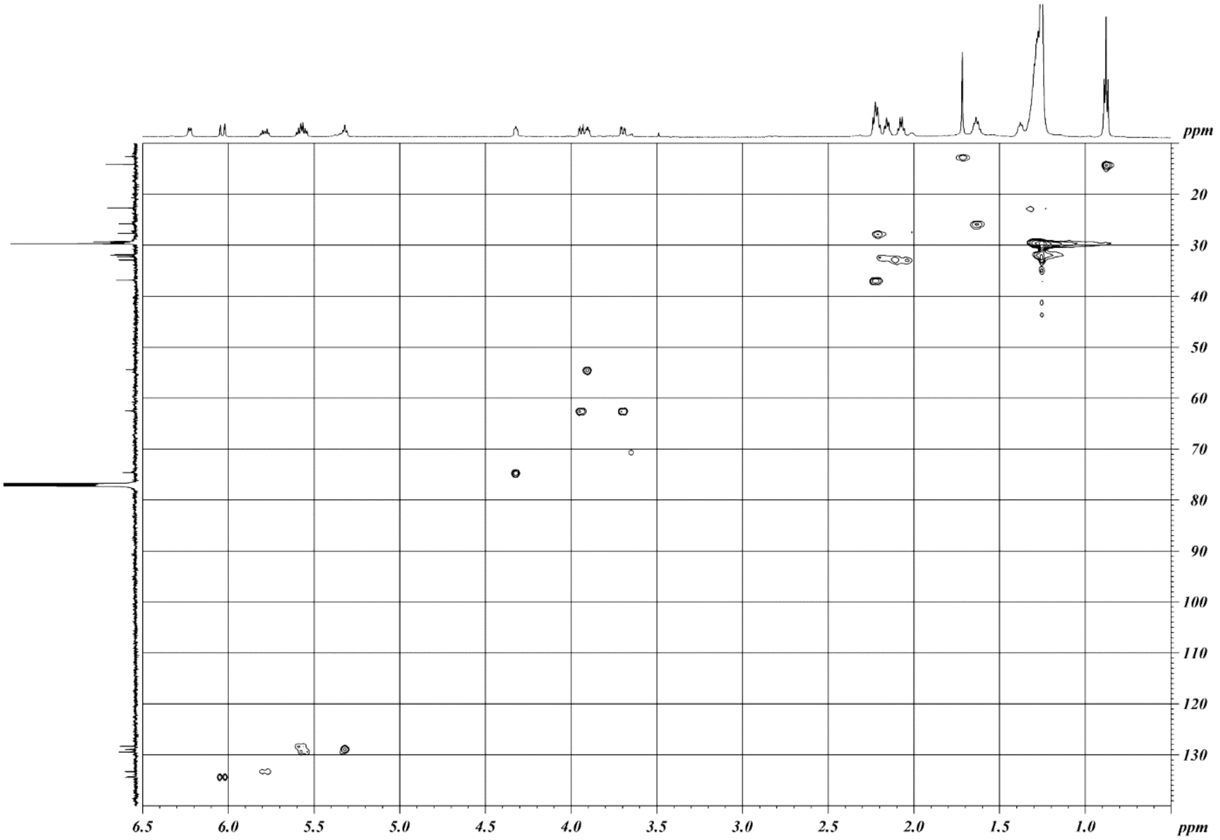

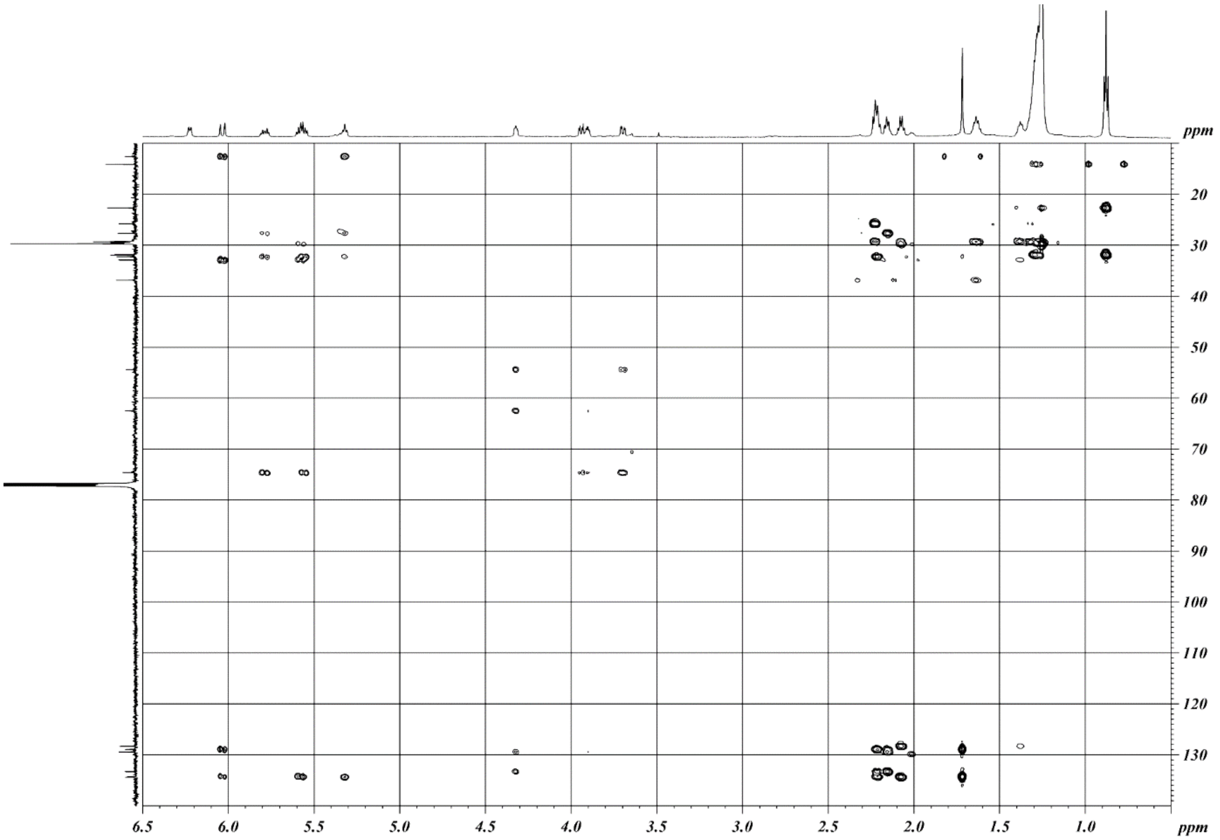

**Supplemental Figure S11.** Two-dimensional NMR correlations of Pi-Cer B (thick bonds: DQF-COSY, curved arrows: HMBC, l + m =15). Spectra ($CDCl_3$, 600 MHz) are DQF-COSY, HMQC, and HMBC (from top).

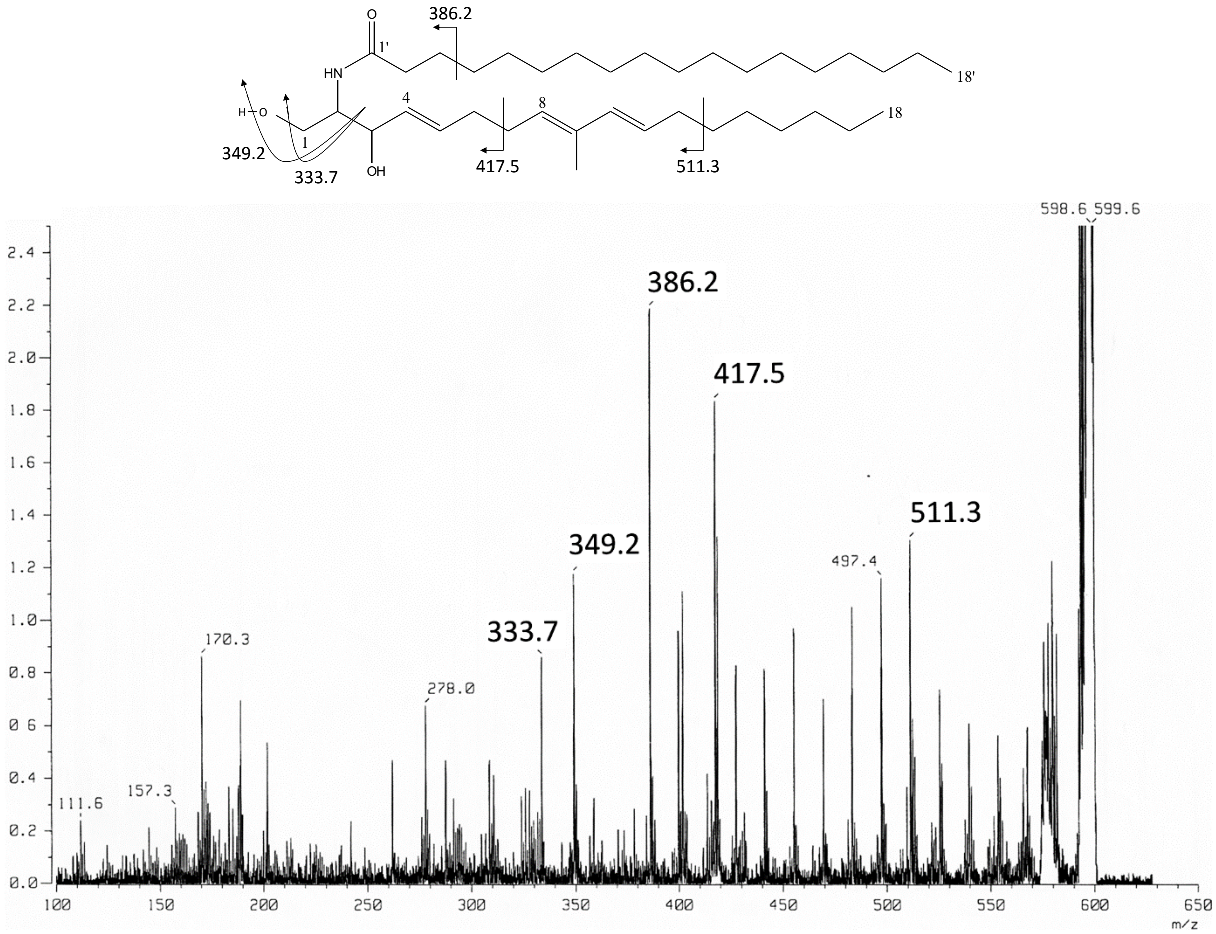


**Supplemental Figure S12.** MS/MS fragmentation of Pi-Cer B (precursor ion: $[M + Na]^+$).

***Pi-Cer C.*** The molecular formula was determined as $C_{38}H_{73}NO_3$ by HR ESI-TOF-MS. The two-dimensional NMR (HMBC, DQF-COSY) revealed the amide bond and the carbon skeletons except for 19 methylenes (Supplemental Figure S13).

The fatty acyl moiety was determined as (*Z*)-docos-13-enoic acid (m = 8 and n = 4 in Supplemental Figure S13) by the same method used for the analysis of Pi-Cer D as described later. The *E* geometry at C-4 was determined by the coupling constant of 15.2 Hz. The absolute configuration was determined to be 2*S*,3*R* by the same method as that for Pi-Cer D. This compound was found to be the known compound.[1]

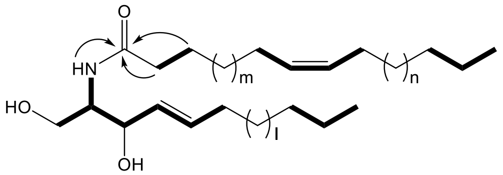
O
HN
m
n
HO
OH
l

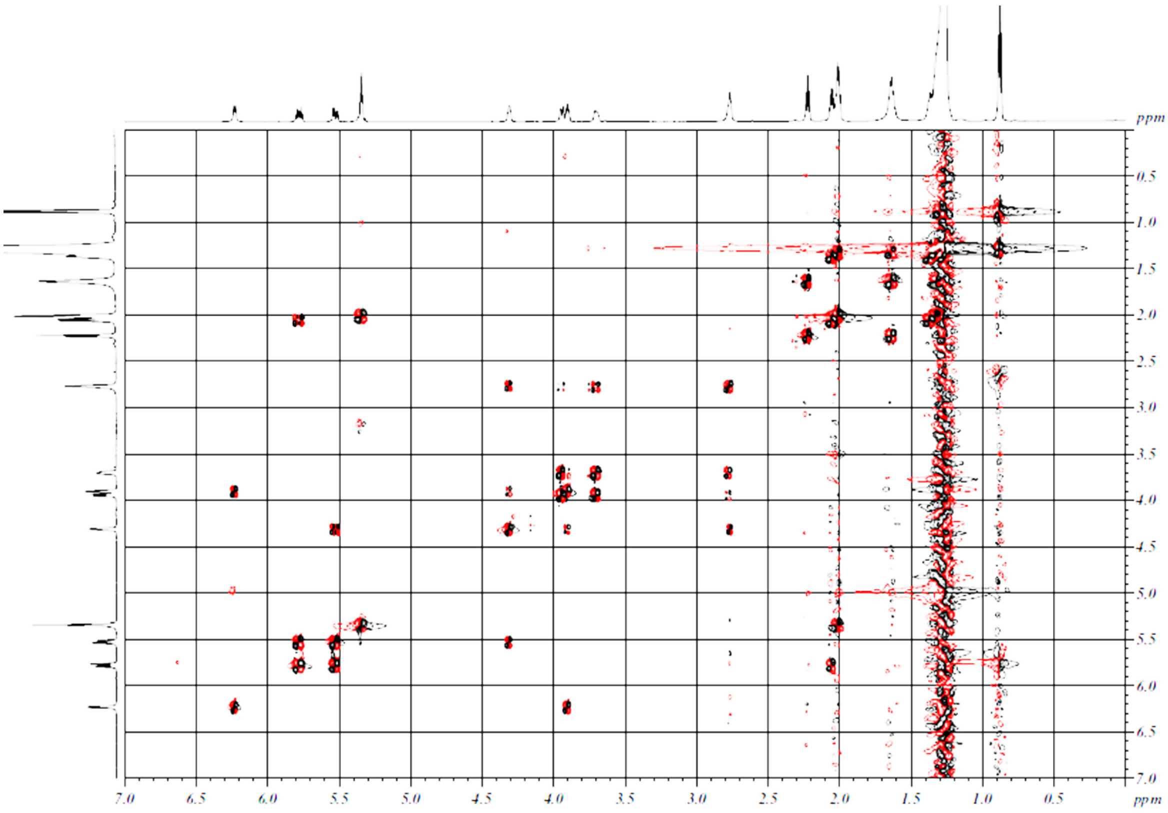
ppm
7.0
6.5
6.0
5.5
5.0
4.5
4.0
3.5
3.0
2.5
2.0
1.5
1.0
0.5
ppm

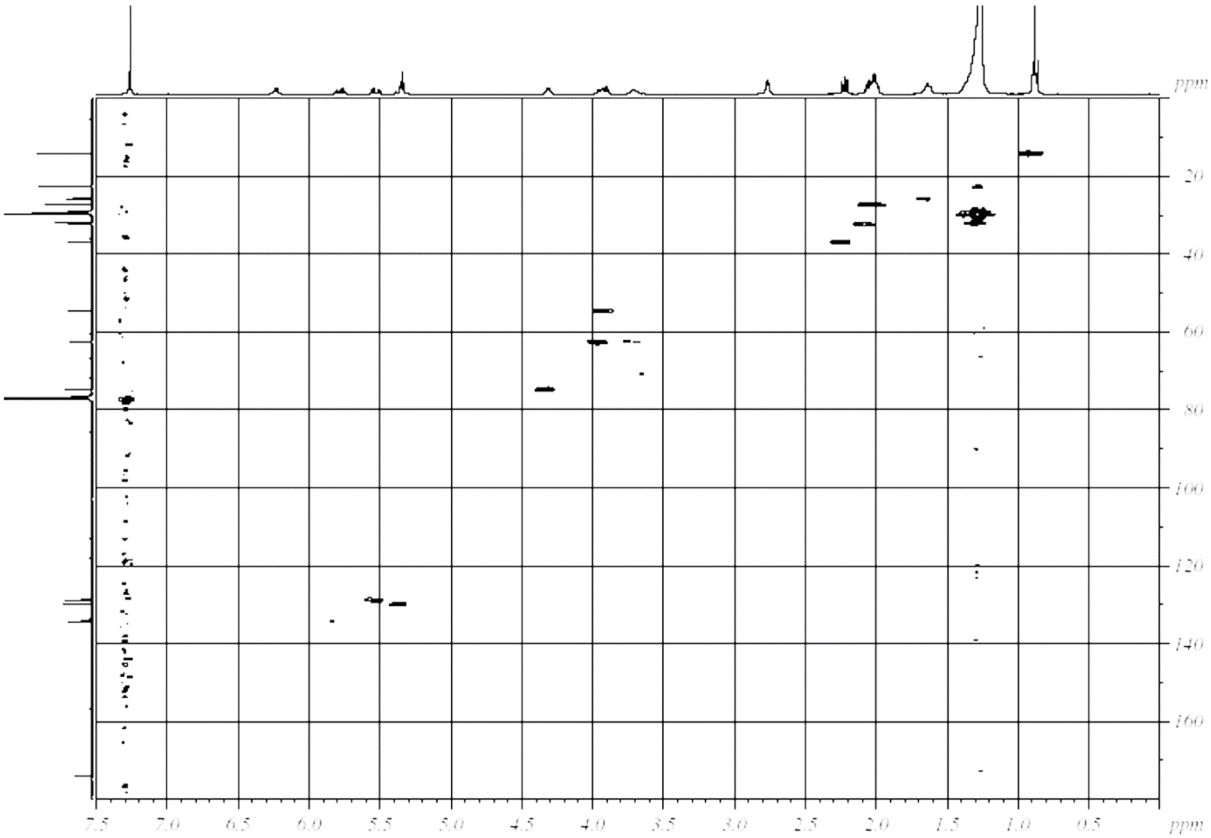
ppm
20
40
60
80
100
120
140
160
7.5
7.0
6.5
6.0
5.5
5.0
4.5
4.0
3.5
3.0
2.5
2.0
1.5
1.0
0.5
ppm

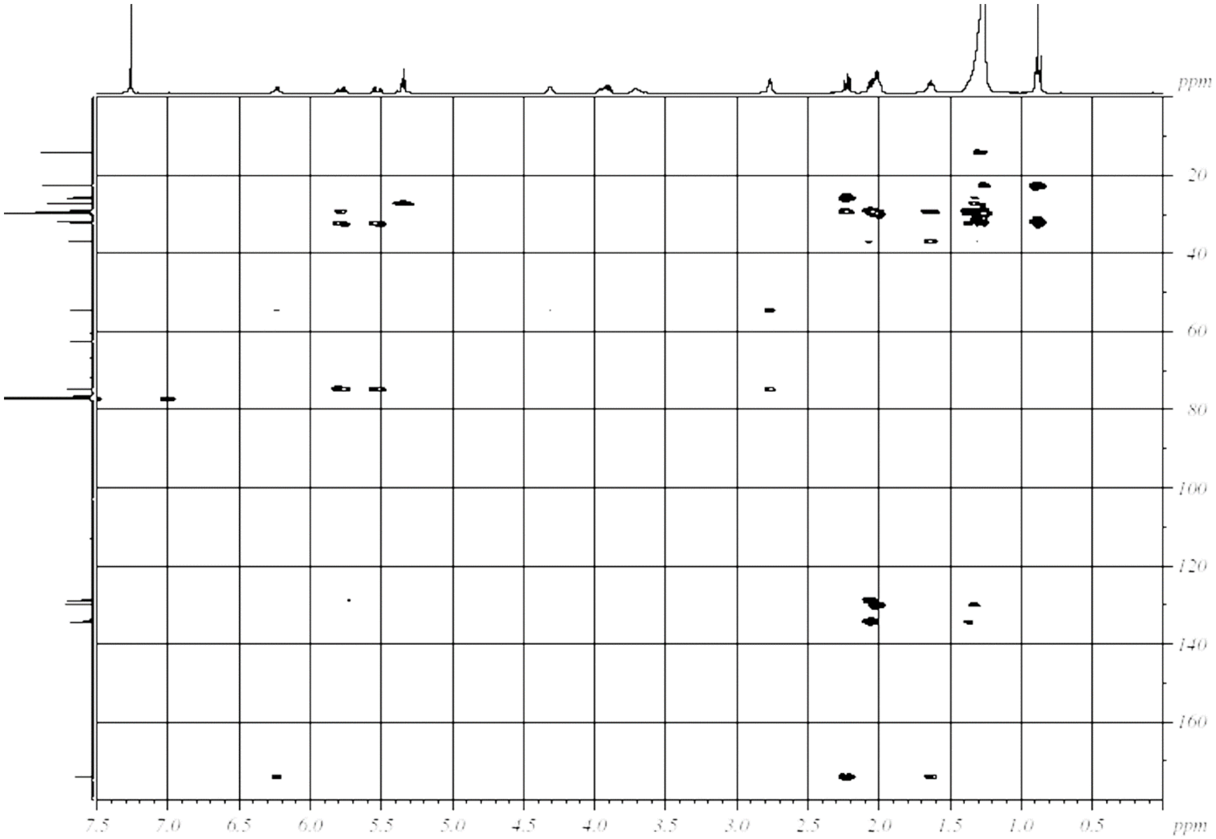


**Supplemental Figure S13.** Two-dimensional NMR correlations of Pi-Cer C (thick bonds: DQF-COSY, curved arrows: HMBC, l + m + n = 19). Spectra (in $CDCl_3$) are DQF-COSY (600 MHz), HMQC (400 MHz), and HMBC (400 MHz) (from top).

***Pi-Cer D***. The molecular formula of Pi-Cer D was found to be $C_{41}H_{75}NO_3$ by using its molecular ion peak at *m/z* 630.5845 $[M + H]^+$ of high-resolution MS. The IR spectrum suggested the presence of hydroxyl (3303 $cm^{-1}$), amide (1644, 1548 $cm^{-1}$), and olefinic groups (962, 721 $cm^{-1}$). The absorption maximum at 235 nm suggested the presence of a conjugated diene. A DQF-COSY spectrum revealed the spin systems indicated by bold bonds in Supplemental Figure S14. An HMBC experiment helped to determine additional connectivity as shown in Supplemental Figure S14. The remaining structural components, which include 15 methylene units, must be inserted between the defined substructures mentioned above. The 8*E*,10*E* geometry was determined by the NOESY correlations shown in Supplemental Figure S14 and the 13’*Z* geometry was indicated by the relatively high-field shift (δ 27.2) of the allylic carbons (C12’ and C15’). The methylene-chain length was determined by negative MS/MS analysis of fatty acid that was obtained by acid hydrolysis of Pi-Cer D (6 M HCl, 110 °C, 4 h). The distinct product ion peaks at *m/z* 251, 237, and 183 in the MS/MS data of the resulting fatty acid (Supplemental Figure S15) clearly indicated the methylene numbers in the carbon chains. The relative 2*S**,3*R** stereochemistry was determined by comparison of the $^1$H NMR data of this compound and some known ceramide compounds; the typical signal at H-1 shows a single one for 2*S**,3*R** or a splitting one for 2*R**,3*R**.[2,3] The absolute configuration of 2*S*,3*R* was then determined by comparison of the sign of specific rotations of this compound ($[\alpha]^{29}_{D}$ -1.7) and some known ceramide compounds; the rotations of 2*S*,3*R* isomers are all negative.[2,4]

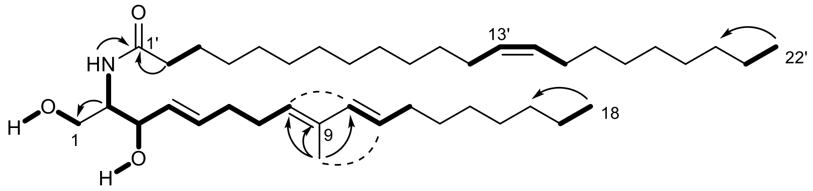


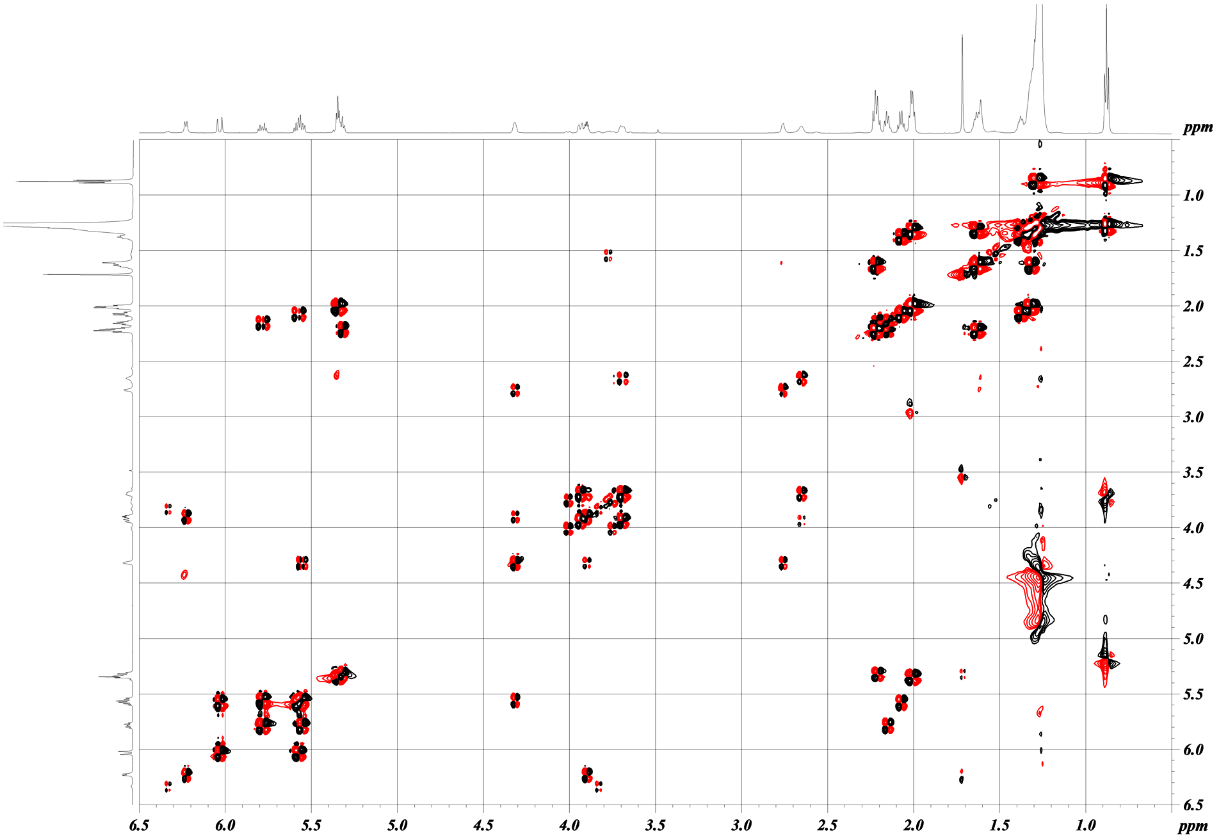

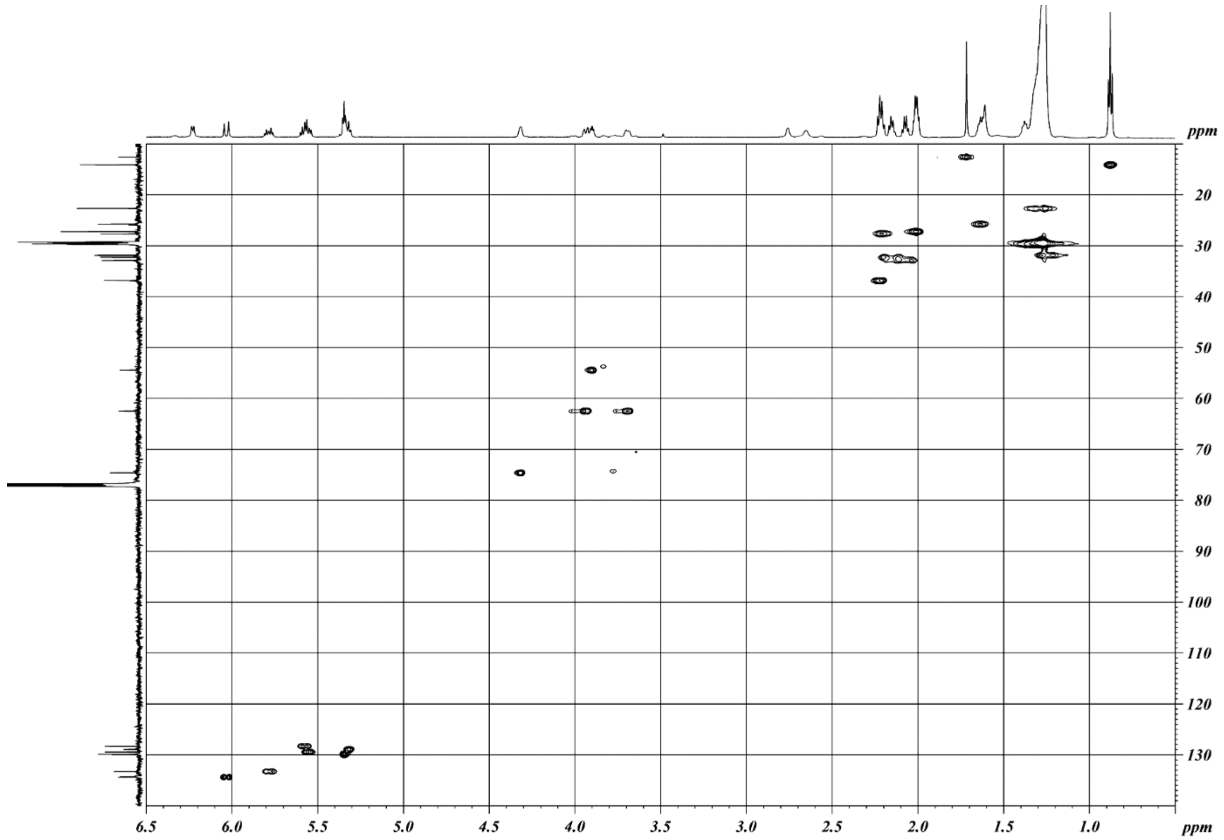
ppm
20
30
40
50
60
70
80
90
100
110
120
130
6.5
6.0
5.5
5.0
4.5
4.0
3.5
3.0
2.5
2.0
1.5
1.0
ppm

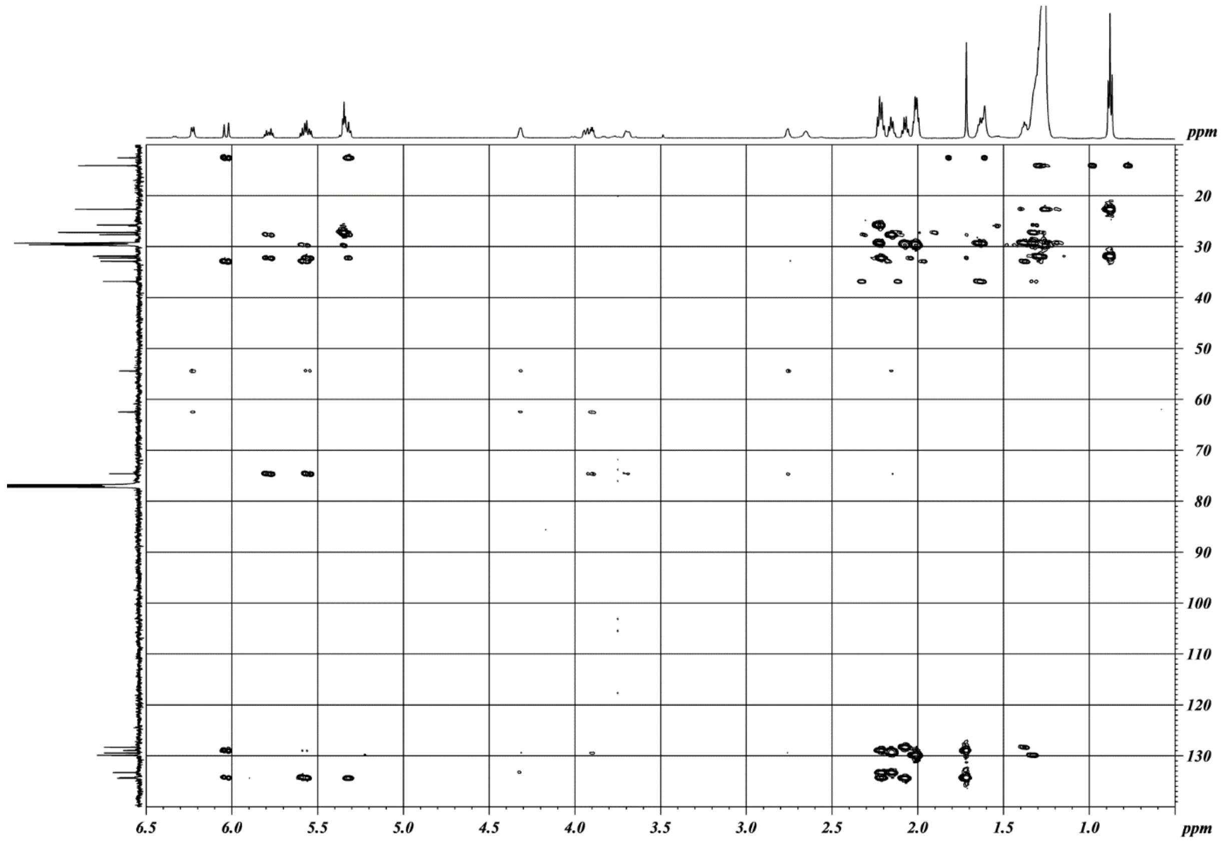
ppm
20
30
40
50
60
70
80
90
100
110
120
130
6.5
6.0
5.5
5.0
4.5
4.0
3.5
3.0
2.5
2.0
1.5
1.0
ppm

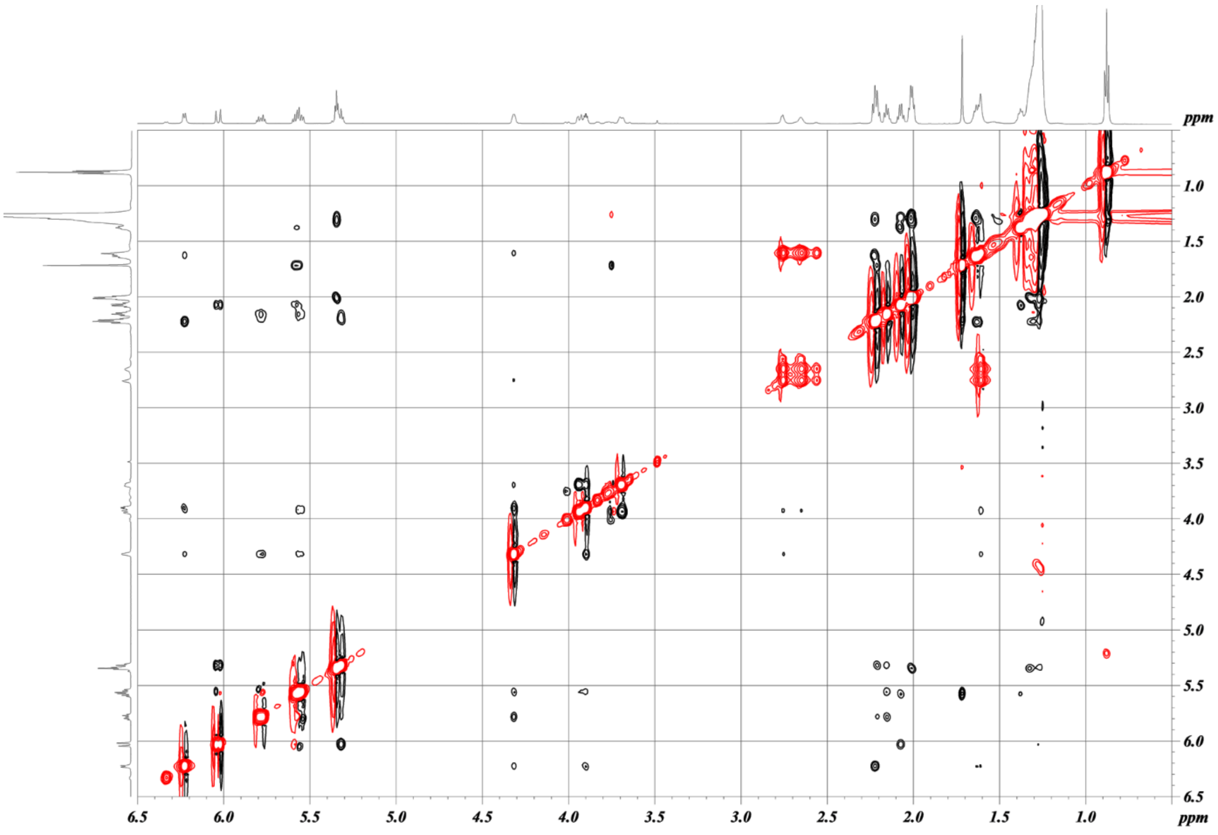

**Supplemental Figure S14**. DQF-COSY (thick bonds), HMBC (curved arrows), and NOESY (dashed curves) correlations of Pi-Cer D. Spectra ($CDCl_3$, 600 MHz) are DQF-COSY, HMQC, HMBC, and NOESY (from top).

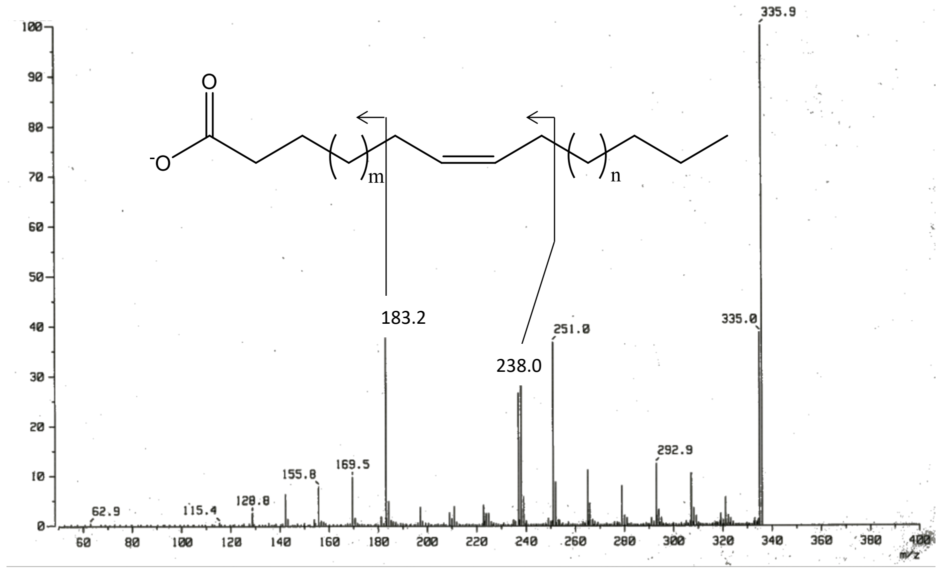


**Supplemental Figure S15**. Linked-scan FAB MS/MS (negative ion mode) of the fatty acid derived from Pi-Cer D by acid hydrolysis.

***Pi-CerPE A***. The molecular formula was determined as $C_{37}H_{71}N_2O_6P$ by HR ESI-TOF-MS. NMR data were quite similar to Pi-CerPE B described in the following section, and the difference is only the lack of $C_2H_4$ in a carbon chain. Since the structural relationship between Pi-Cer A and Pi-CerPE A is the same as that between Pi-Cer B and Pi-CerPE B, this compound was easily identified as the phosphoethanolamine derivative of Pi-Cer A. This compound was found to be the known compound.[5]

***Pi-CerPE B***. The molecular formula was determined as $C_{37}H_{71}N_2O_6P$ by HR ESI-TOF-MS. NMR data was similar to that of Pi-Cer B except for an additional ethylene group [-$CH_2$-$CH_2$-, δ 4.05 (m, 2H, H-1") and 3.11 (br s, 2H, H-2")]. Another difference is that the molecular formula was larger than Pi-Cer B by $C_2H_2NO_3P$. This chemical relationship was the same as that between Pi-Cer D and Pi-CerPE D. Therefore, it was concluded that Pi-CerPE B was the phosphoethanolamine derivative of Pi-Cer B.

***Pi-CerPE C***. The difference of the molecular formula, $C_{40}H_{79}N2O_6P$, and NMR data between Pi-Cer C and Pi-CerPE C was exactly the same as that between other Pi-Cers and Pi-CerPEs. The 2*S*,3*R* configuration was determined by comparison of the specific rotations between the Cer product from Pi-CerPE C treated with sphingomyelinase (from *Bacillus cereus*, Sigma) and Pi-Cer C. Therefore, this compound was determined to be the phosphoethanolamine derivative of Pi-Cer C and found to be the known compound.[5]

***Pi-CerPE D***. The molecular formula $C_{43}H_{81}N_2O_6P$ of Pi-CerPE D was determined on the basis of its molecular ion peak at *m/z* 753.5932 $[M+H]^+$. The NMR spectra were similar to those of Pi-Cer D. The only difference was the presence of an additional ethylene group [-$CH_2$-$CH_2$-, δ 4.05 (m, 2H, H-1") and 3.10 (br s, 2H, H-2")], which was confirmed by DQF-COSY (Supplemental Figure S16). The other NMR signals and 2D NMR correlations were superimposable to those of Pi-Cer D (Supplemental Figure S16). Considering the molecular formula of this compound and the chemical shifts of the ethylene group, the remaining functionality could be $NH_2$ and $PO_3H$. The IR absorption at 1222 $cm^{-1}$ suggested the presence of a phosphate group. Therefore, the structure of Pi-CerPE D was concluded as shown. The carbon-chain length was determined by the same method as those for Pi-Cer D. The absolute configuration of 2*S*,3*R* was determined by comparison of the $^1H$ NMR and specific rotation of this compound ($[\alpha]_D$ +9.2) and the known compound Pi-CerPE C ($[\alpha]_D$ +9.4).

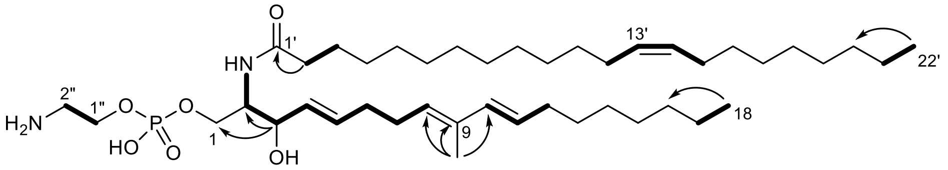
O
1'
13'
22'
HN
2"
1"
H2N
O
P
O
HO
O
1
OH
9
18

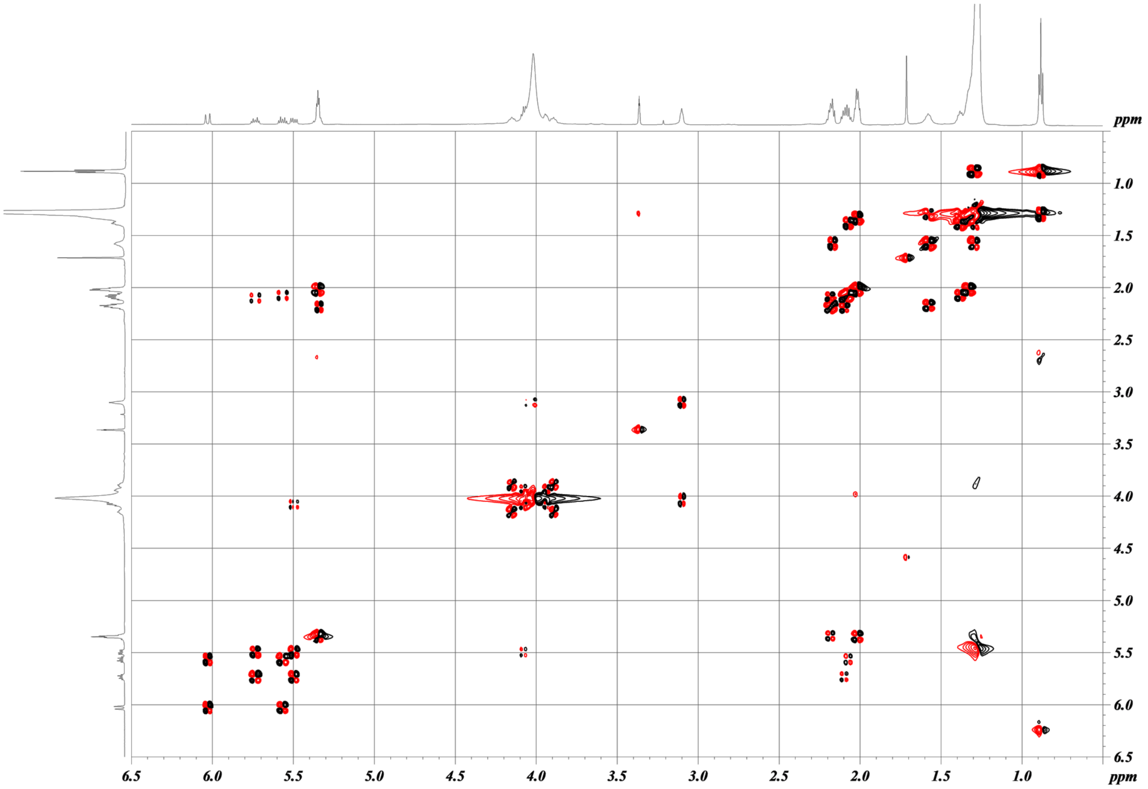
ppm
ppm

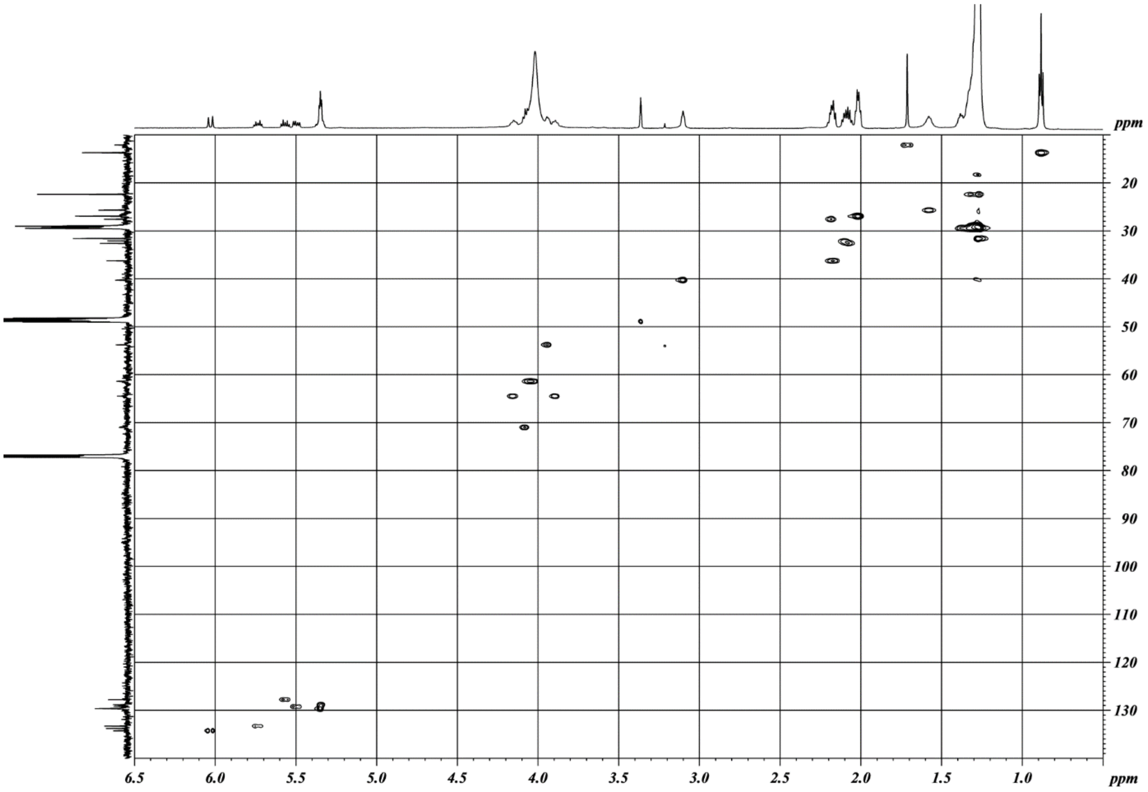
ppm
ppm

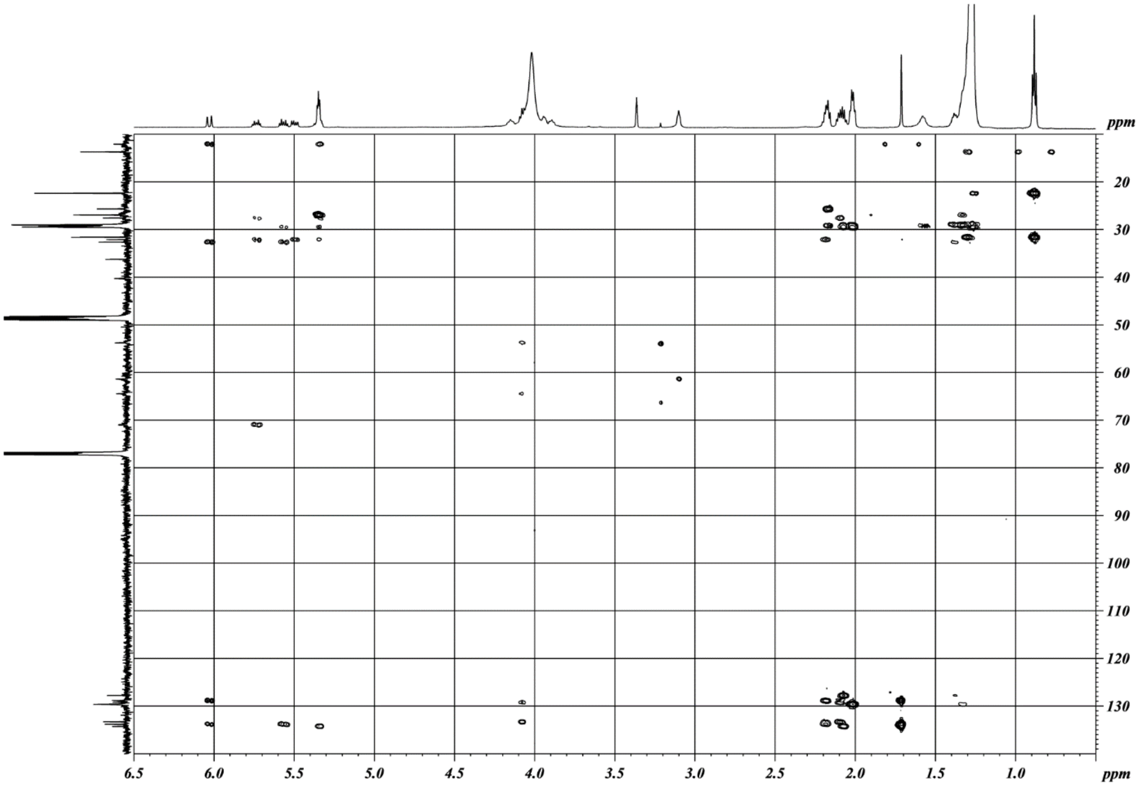


**Supplemental Figure S16**. DQF-COSY (thick bonds) and HMBC (curved arrows) correlations of Pi-CerPE D. Spectra ($CDCl_3$:$CD_3OD$=4:1, 600 MHz) are DQF-COSY, HMQC, and HMBC (from top).

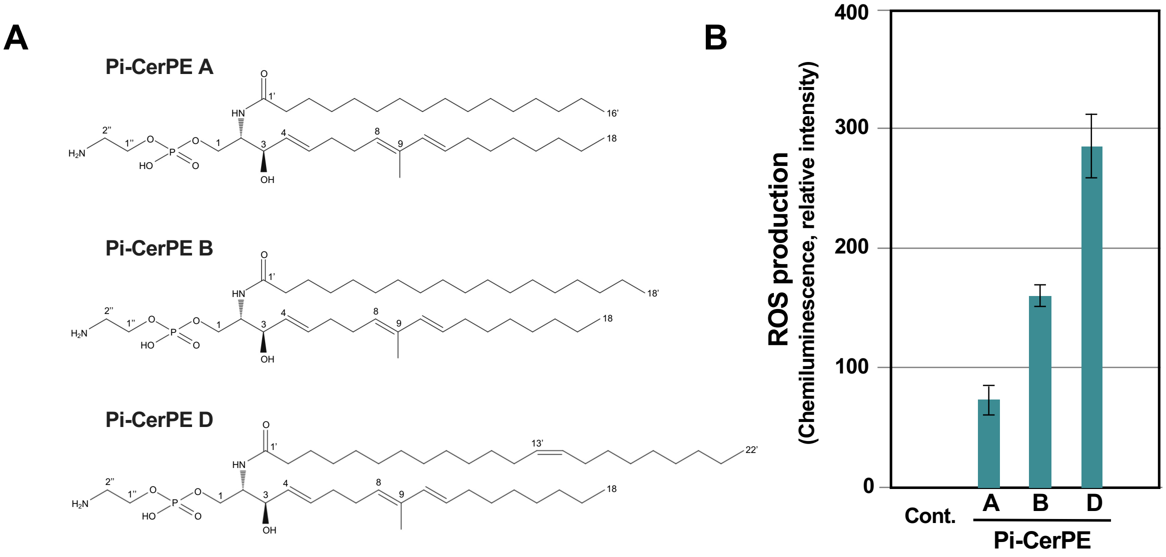


**Supplemental Figure S17.** Ceramide phosphoethanolamine (Pi-CerPE) elicitors purified from methanol extract of *P. infestans* mycelia (Pi-MEM), which can induce the production of reactive oxygen species (ROS) in potato suspension cultured cells. **(A)** Structures of Pi-CerPE A, B and D. See Supplemental Figure S1 for the procedures of purification of elicitors and Supplemental Figures S6, 7, 9, 16 and Supplemental document for details of their structural analysis. **(B)** Potato suspension cultured cells were treated with 0.3% DMSO (Cont.) or 3 µg/ml Pi-CerPE A, B or D and production of ROS was detected as L-012-mediated chemiluminescence 3 h after the treatment. Scores shown are chemiluminescence intensities relative to that of Pi-MEM-treated cells in Figure 2.

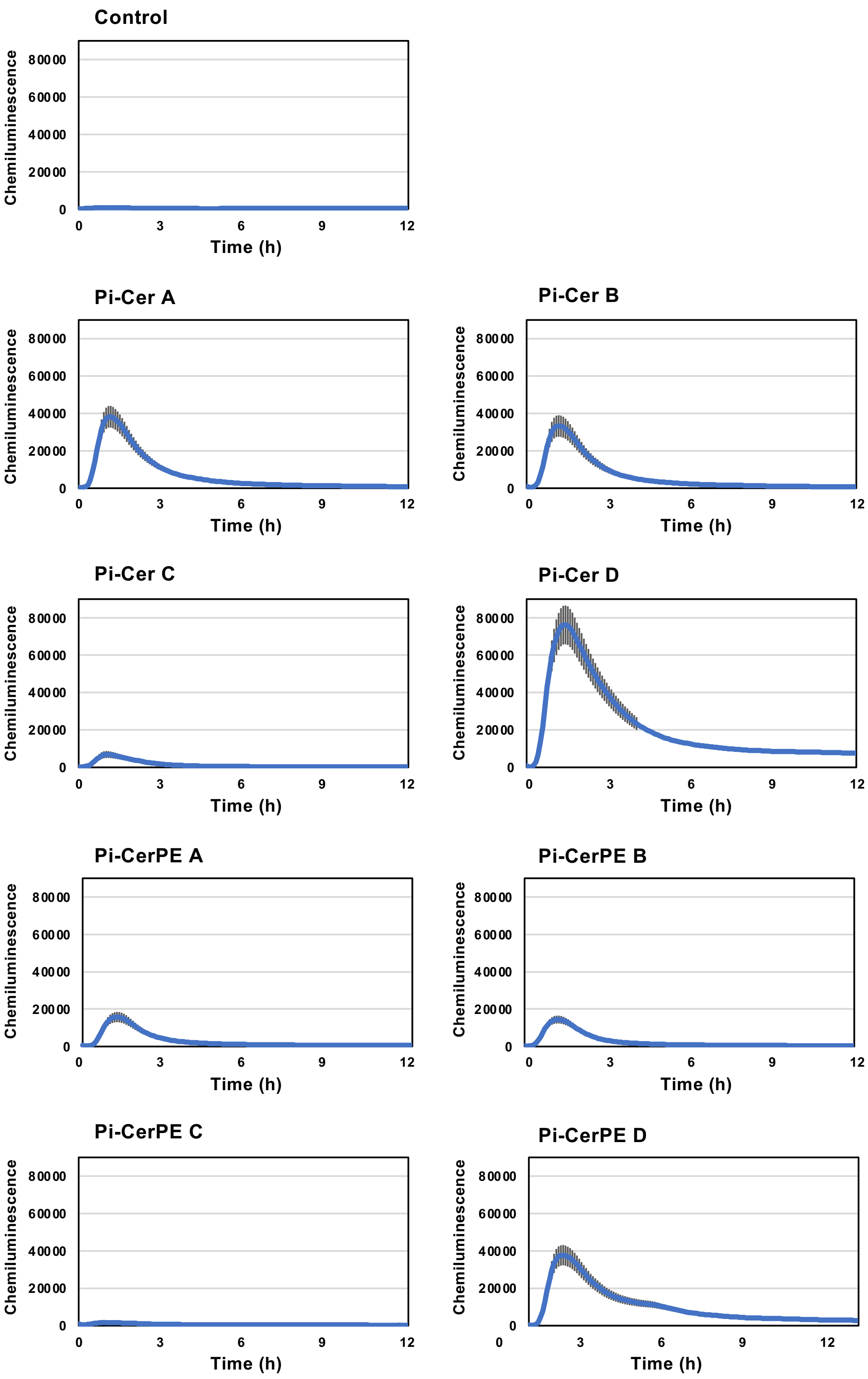


**Supplemental Figure S18.** Activation of *A. thaliana WRKY33* promoter by Pi-Cer and Pi-CerPE elicitors. Arabidopsis transformant pWRKY33-LUC containing *LUC* transgene under the control of *AtWRKY33* (AT2G38470.1) promoter was treated with 20 µg/ml Pi-Cer A, B, C or D and chemiluminescence was monitored for 12 h after the treatment. Data are means ± SE (n = 8).

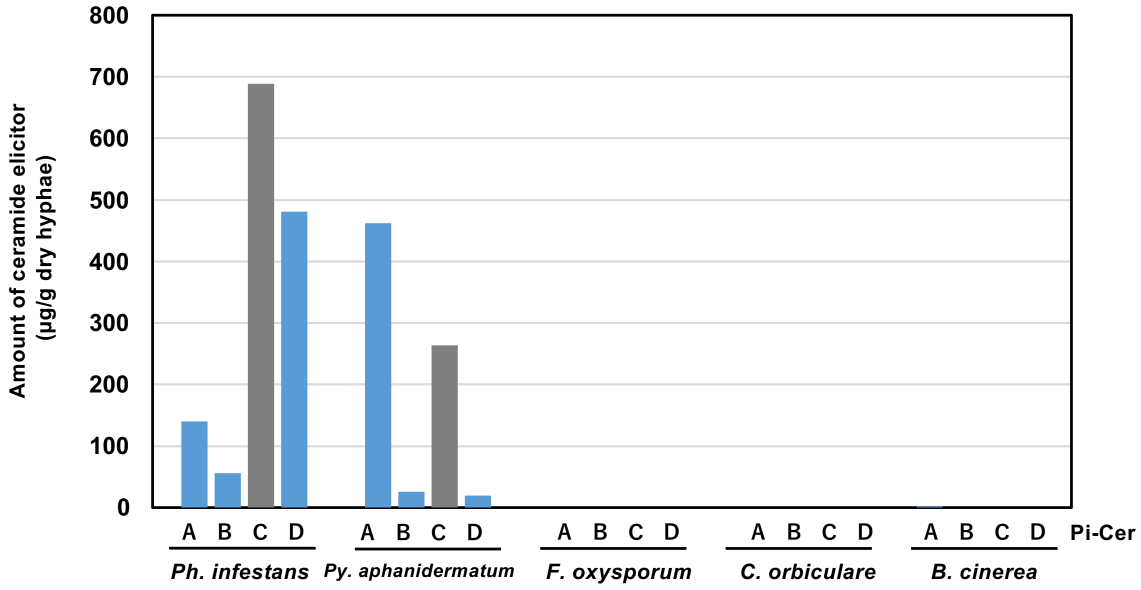


**Supplemental Figure S19.** Pi-Cer contents in mycelia of oomycete and fungal plant pathogens. Pi-Cers were partially purified from mycelia of oomycete (*Phytophthora infestans* and *Pythium aphanidermatum*) and fungal (*Fusarium oxysporum* f. sp. *melonis*, *Colletotrichum orbiculare* and *Botrytis cinerea*) pathogens, and Pi-Cers contents in these pathogens were estimated by LC/MS.

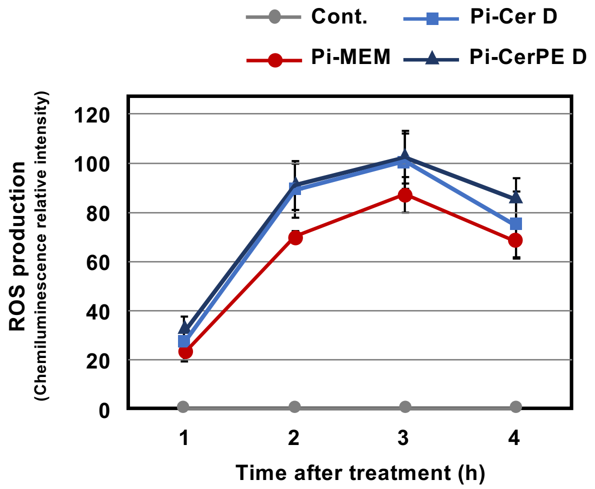


**Supplemental Figure S20.** Induction of reactive oxygen species (ROS) production by Pi-Cer D and Pi-CerPE D treatment in potato suspension-culture cells. Potato cells were treated with 300 ng/ml Pi-Cer D, Pi-CerPE D or 3 μg/ml Pi-MEM and ROS production was detected as L-012 mediated chemiluminescence. Data are means ± SE (n = 3).

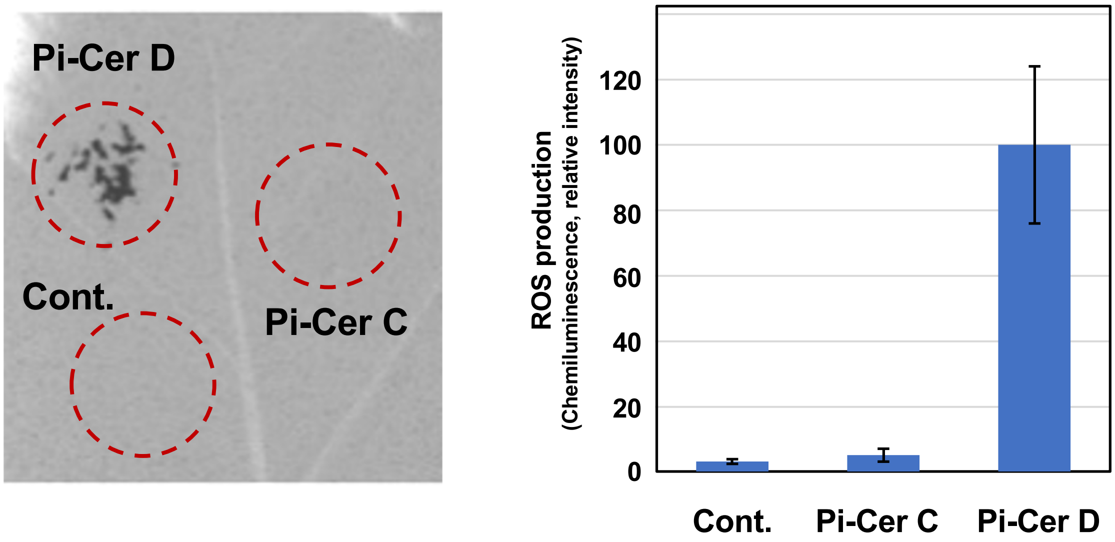


**Supplemental Figure S21.** Pi-Cer D, but not Pi-Cer C, can induce the production of reactive oxygen species (ROS) in *Nicotiana benthamiana*. Leaves of *N. benthamiana* were treated with 1% DMSO (Cont.), 100 µg/ml Pi-Cer C or Pi-Cer D and production of ROS was detected as L-012-mediated chemiluminescence 12 hrs after treatment. Data are means ± SE (n = 10). Scores are chemiluminescence intensities relative to that of Pi-Cer D-treated leaves. Data marked with asterisks are significantly different from control as assessed by two-tailed Student's *t* tests: $^{**}P < 0.01$.

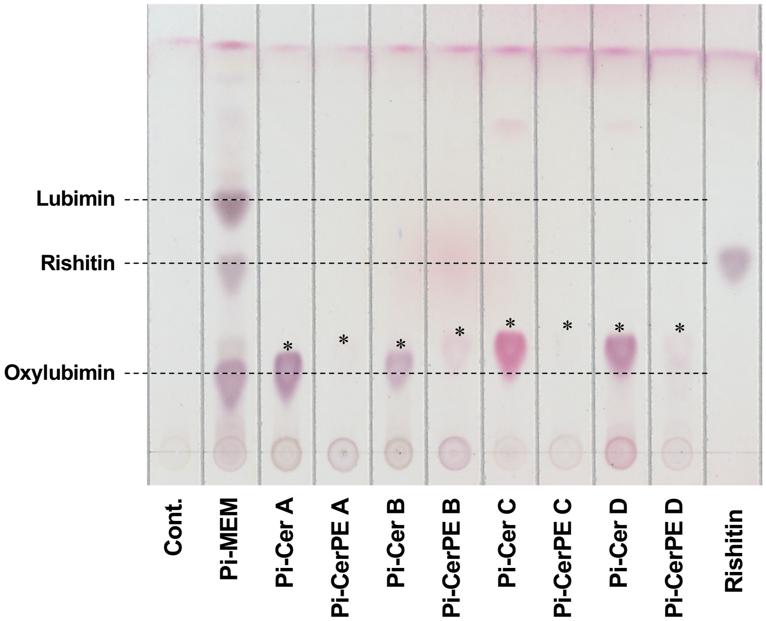


**Supplemental Figure S22.** Production of phytoalexins in potato tuber is not induced by Pi-Cers and Pi-CerPEs. Potato tubers were treated with 10 µg/ml Pi-Cers or Pi-CerPEs and phytoalexins were extracted 48 h after treatment. 0.5 % DMSO and 500 µg/ml Pi-MEM were used as negative and positive controls, respectively. Note that spots marked by asterisks are treated Pi-Cers or Pi-CerPEs.

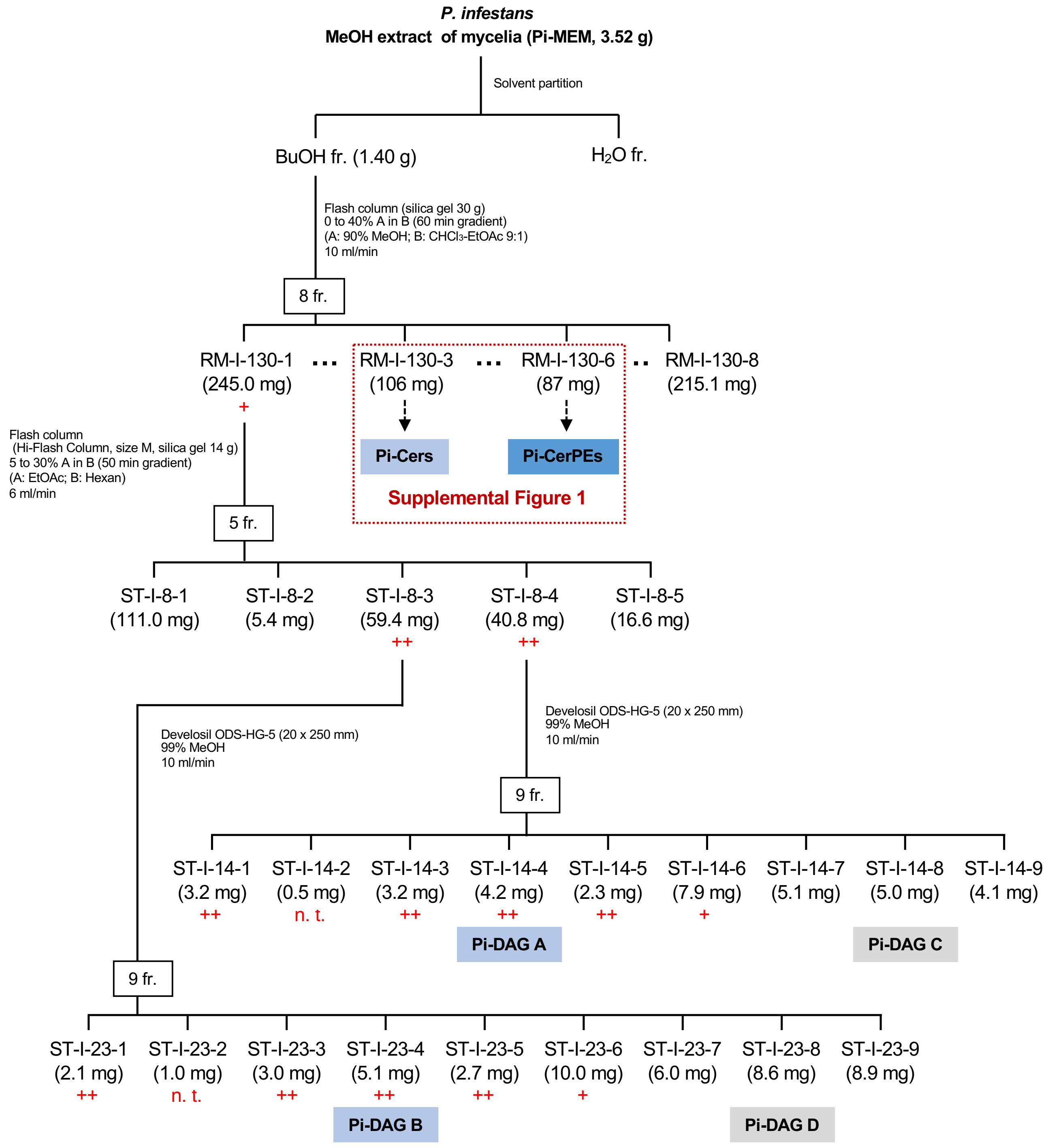


**Supplemental Figure S23.** Purification procedure of *Phytophthora infestans* diacylglycerol (Pi-DAG) elicitors. Methanol extract of *P. infestans* mycelia (Pi-MEM) was dried, and sequentially fractionated with butanol (BuOH) and water. BuOH-soluble fractions were then separated in fractions (fr.) by a series of column chromatography as indicated. Elicitor (phytoalexin-producing) activity (+) was determined by detecting the phytoalexins produced in potato tubers 2 days after treatment by thin-layer chromatography. n. t., not tested.

### Pi-DAG A

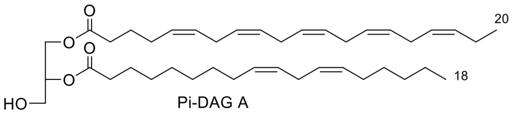


**A**

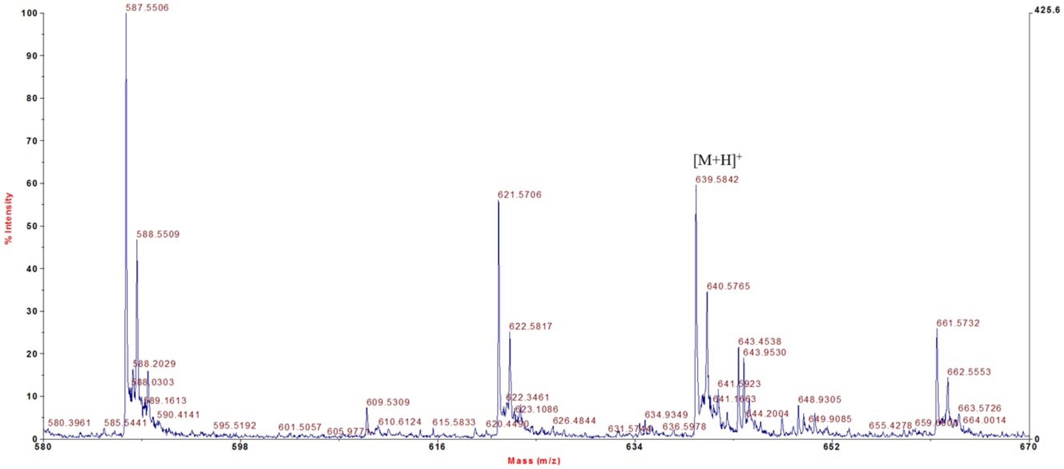


**B**

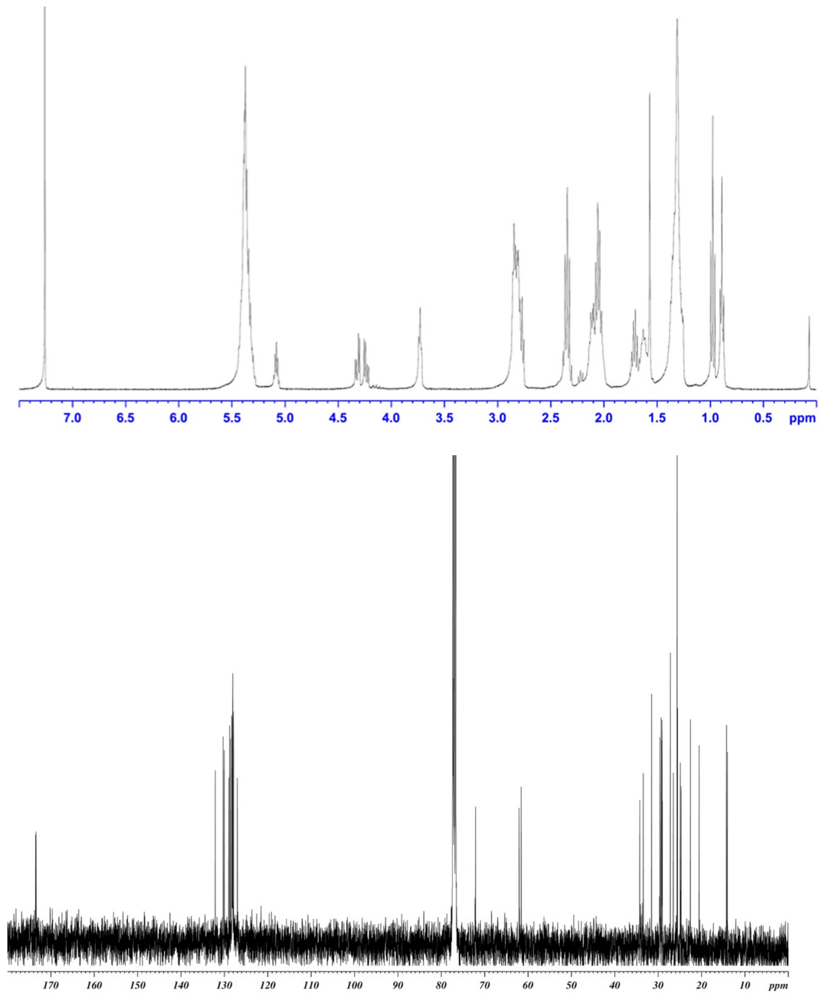


**Supplemental Figure S24.**
**(A)** ESI-TOF MS of Pi-DAG A.
**(B)** NMR spectra of Pi-DAG A ($CDCl_3$, 400 MHz for $^1H$ and 100 MHz for $^{13}C$).

C

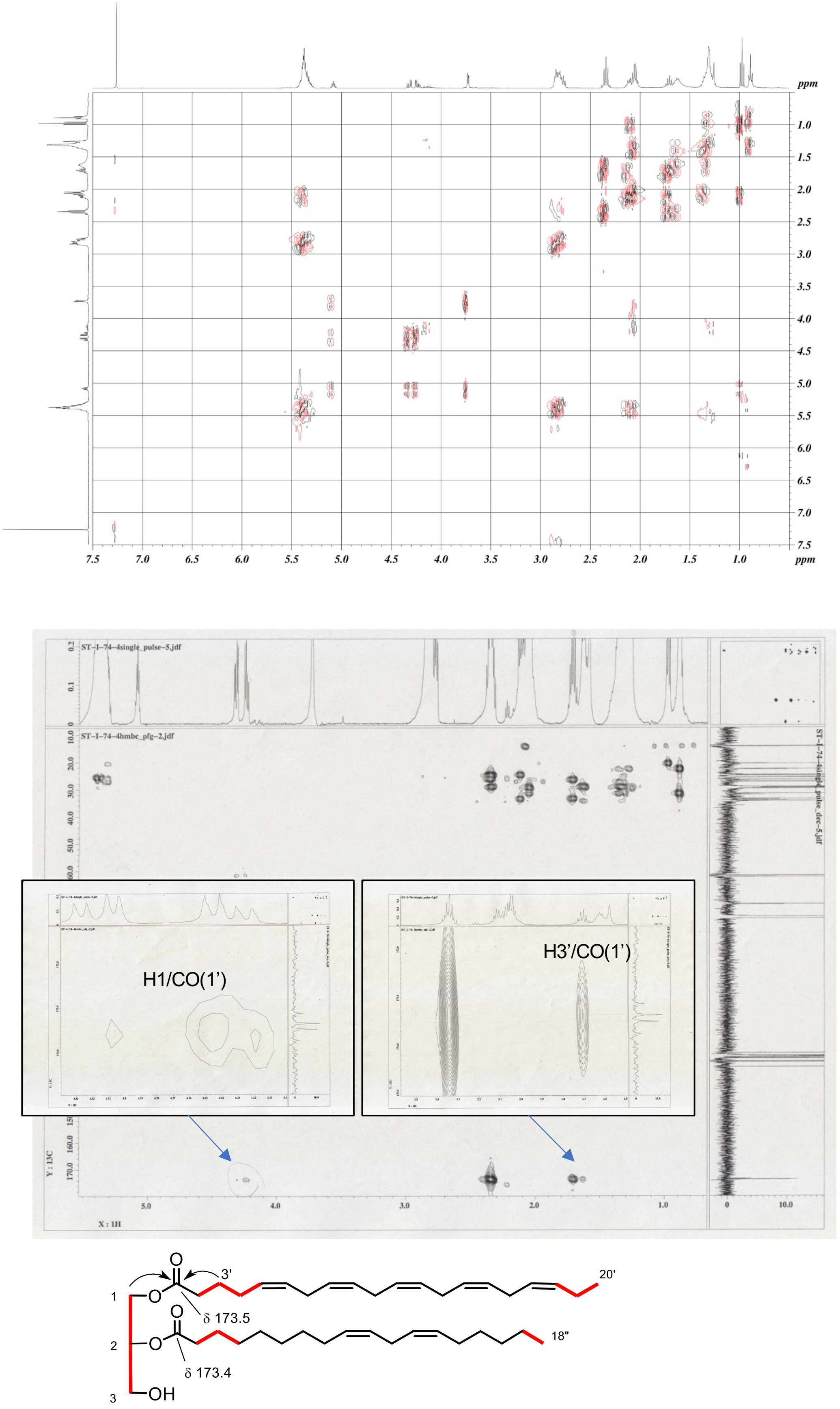


**Supplemental Figure S24.** (continued).
**(C)** Two-dimensional NMR of Pi-DAG A ($CDCl_3$, 400 MHz).
DQF-COSY (top) indicates red-colored connectivity, and the HMBC correlations of H-1/CO(1') and H-3'/CO(1') (bottom, arrows) suggest that eicosapentaenoic acid is located at the position 1.

**D**

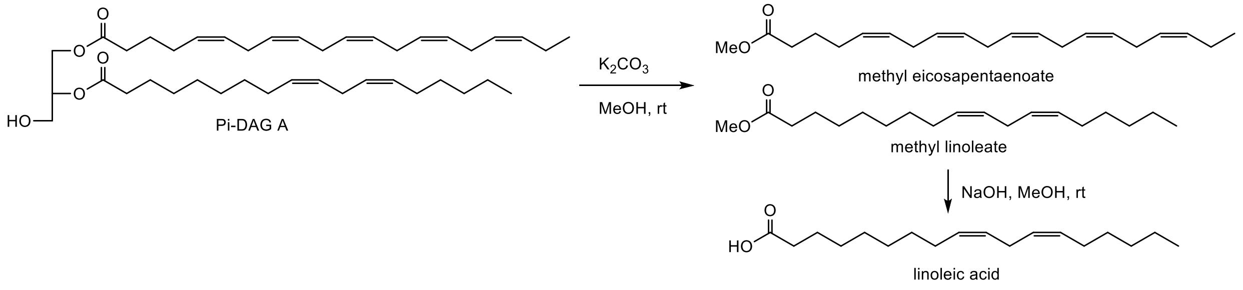


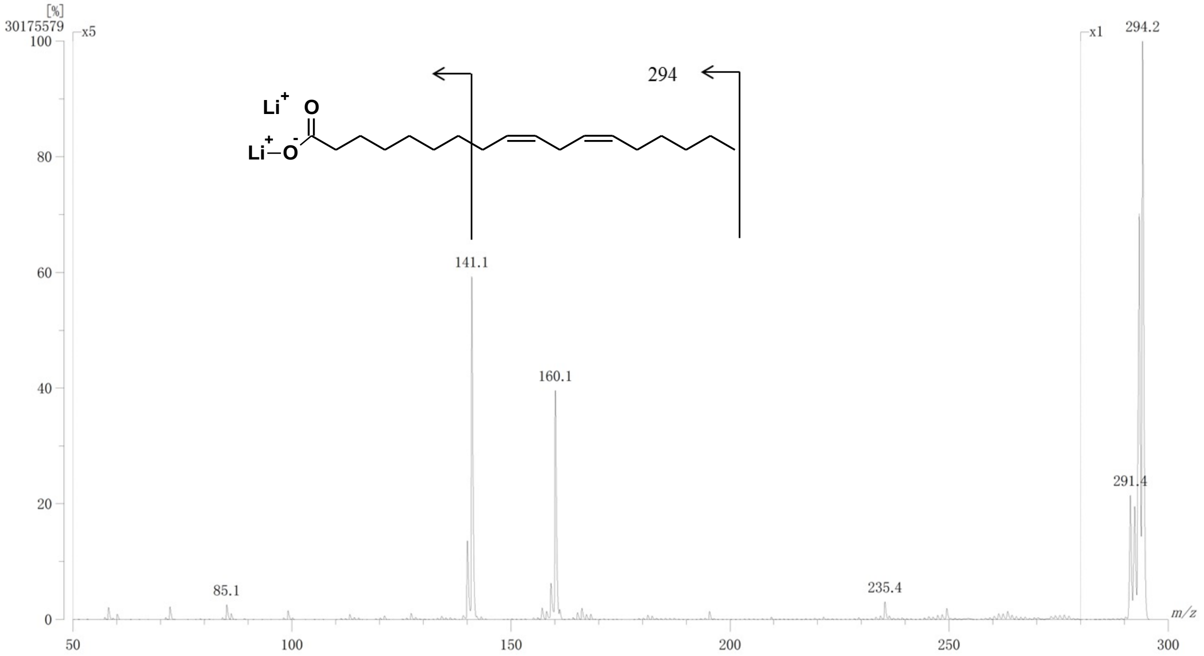


**Supplemental Figure S24.** (continued).
**(D)** Determination of fatty acids of Pi-DAG A by MS/MS.
Methanolysis of Pi-DAG A gave methyl linoleate and methyl eicosapentaenoate. Methyl linoleate was subjected to hydrolysis and the resulting acid was analyzed by MS/MS, revealing the location of the double bonds. The structure of methyl eicosapentaenoate was determined by $^1$H NMR comparison with standard spectrum.

### Pi-DAG B

A

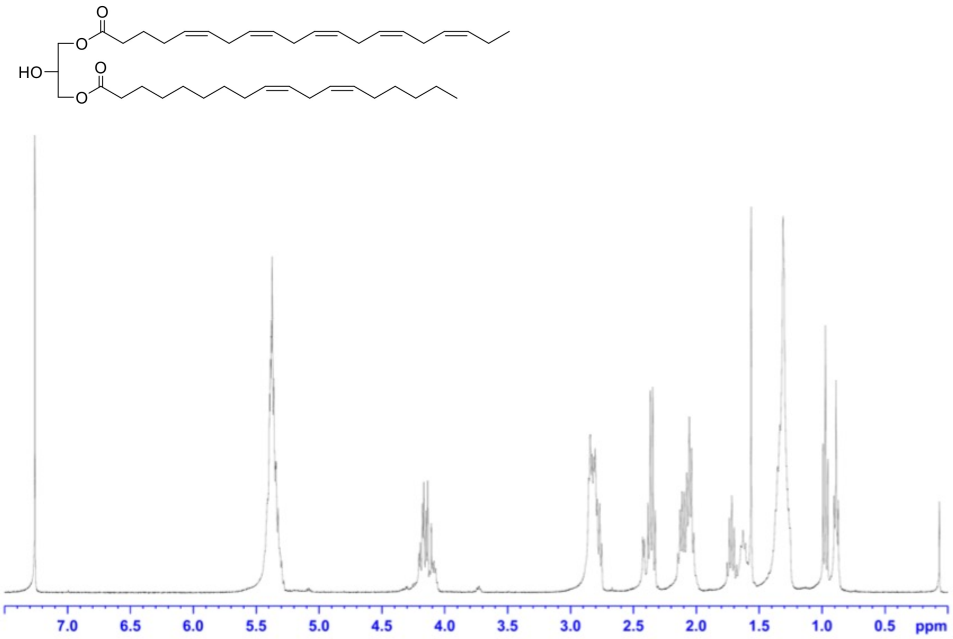


**Supplemental Figure S25. (A)** $^1$H NMR spectrum of Pi-DAG B ($CDCl_3$, 400 MHz). The data indicates that the two fatty acyl moieties are the same as those of Pi-DAG A. The chemical shifts at 4.13 ppm (4H) for H-1 and H-3 indicate that both the primary alcohols of glycerol are acylated.

**B**

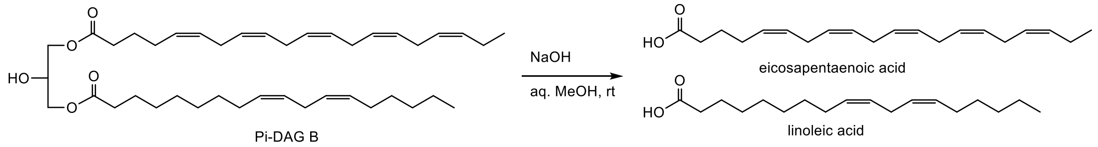


FAB MS(-) of fatty acid mixture

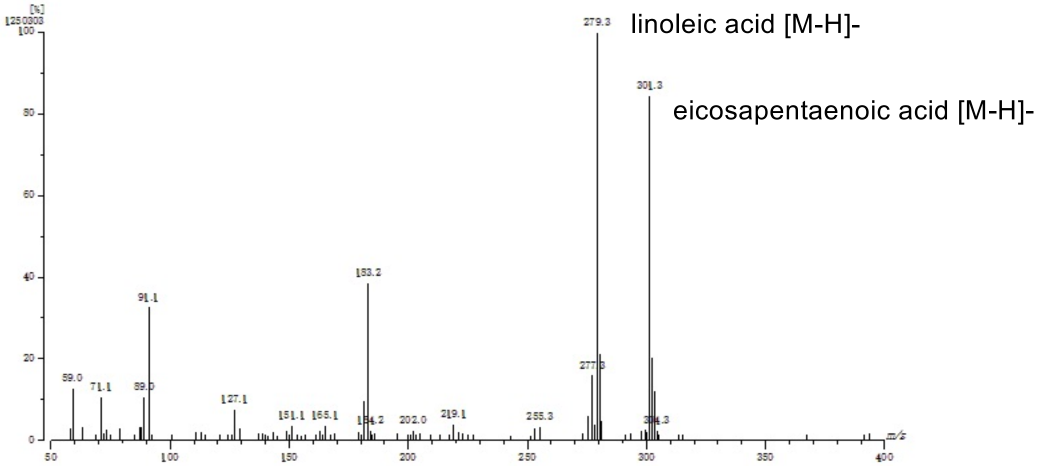


FAB MS/MS(-) of *m/z* 279.3

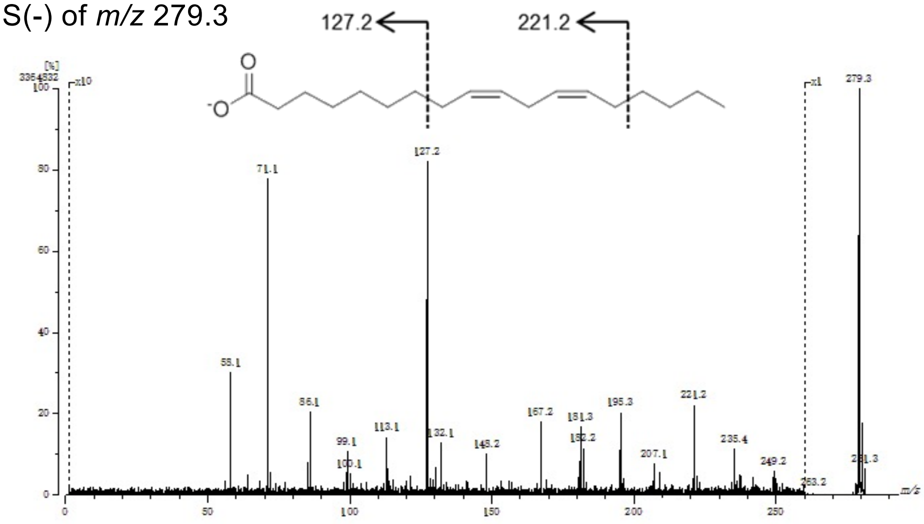


**Figure S25.** (continued).
**(B)** Determination of fatty acids in Pi-DAG B by negative ion FAB MS
Hydrolysis of Pi-DAG B gave two fatty acids as shown in the 1st MS. Linoleic acid was characterized by FAB MS by using the precursor ion m/z 279.3 $[M-H]^-$.

#### Pi-DAG C

A

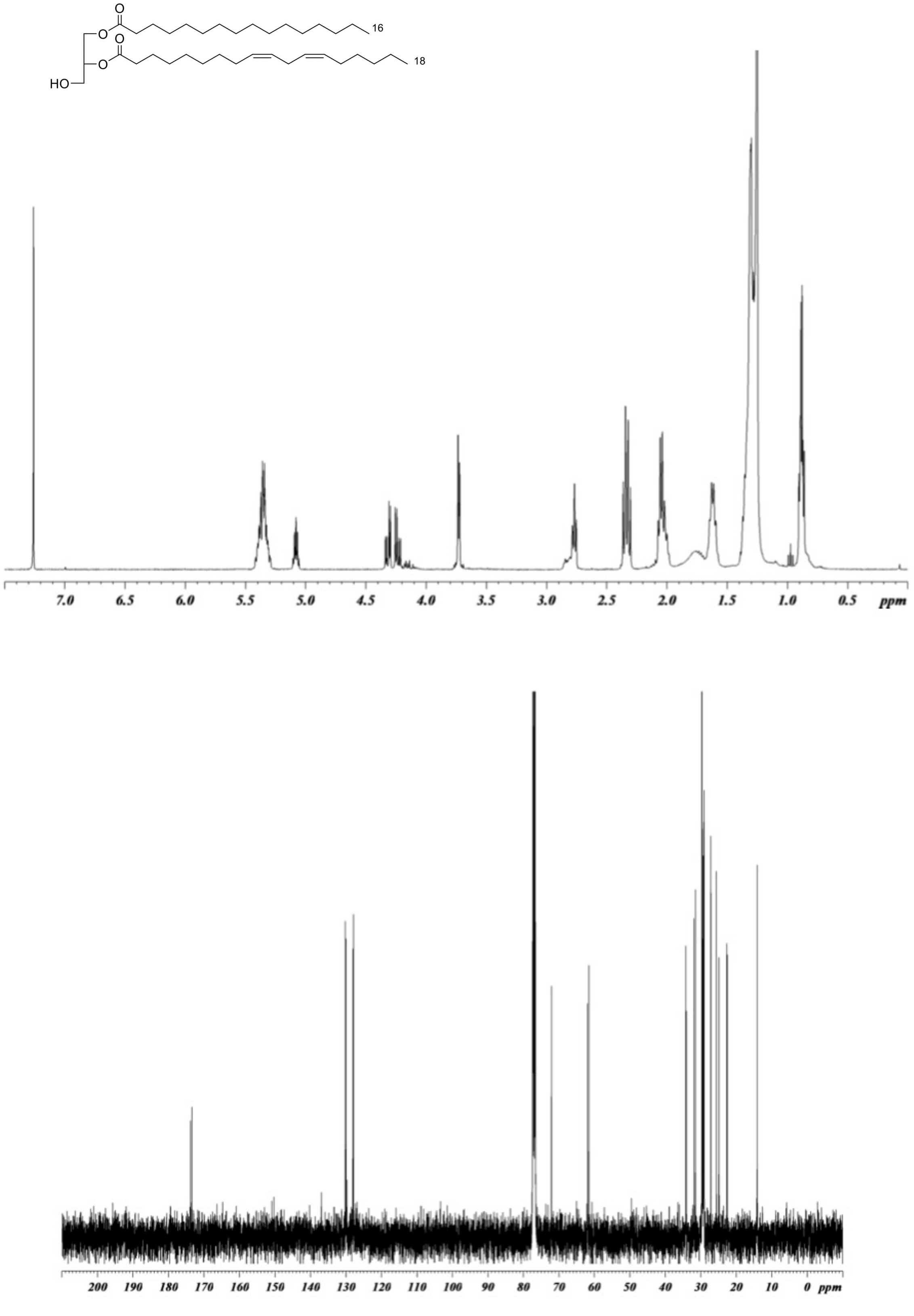


**Supplemental Figure S26. (A)** NMR spectra of Pi-DAG C ($CDCl_3$, 400 MHz for $^1H$, 100 MHz for $^{13}C$).

**B**

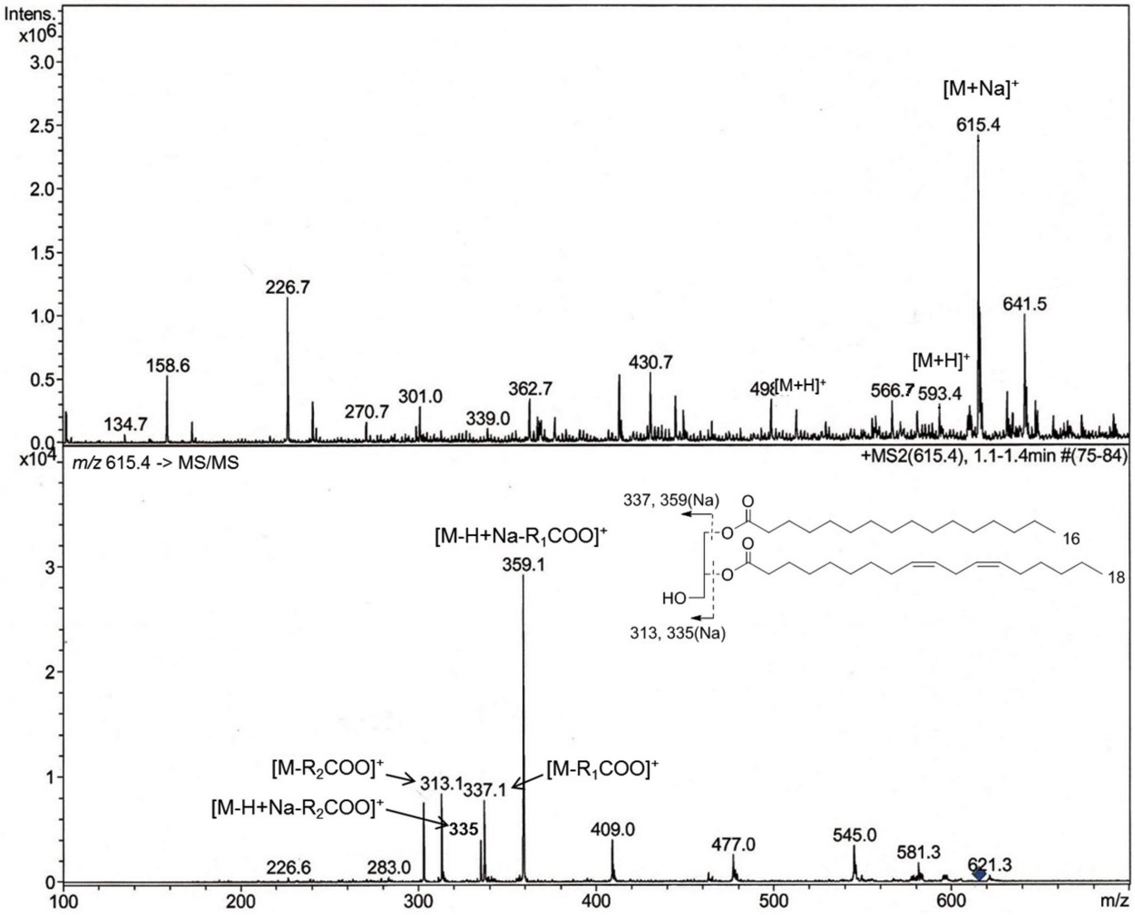


**Figure S26.** (continued).
**(B)** Determination of fatty acids in Pi-DAG C by positive ion ESI MS/MS.
The precursor ion m/z 615.3 [M+Na]+ (top) was used for MS/MS (bottom). The product ions indicate that the fatty acids are palmitic acid and linolein acid. The highest peak at m/z 359.1 suggests that palmitic acid is linked to the position 1, because the ester at position 1 (or 3) of DAG is known to be more easily cleaved.

C

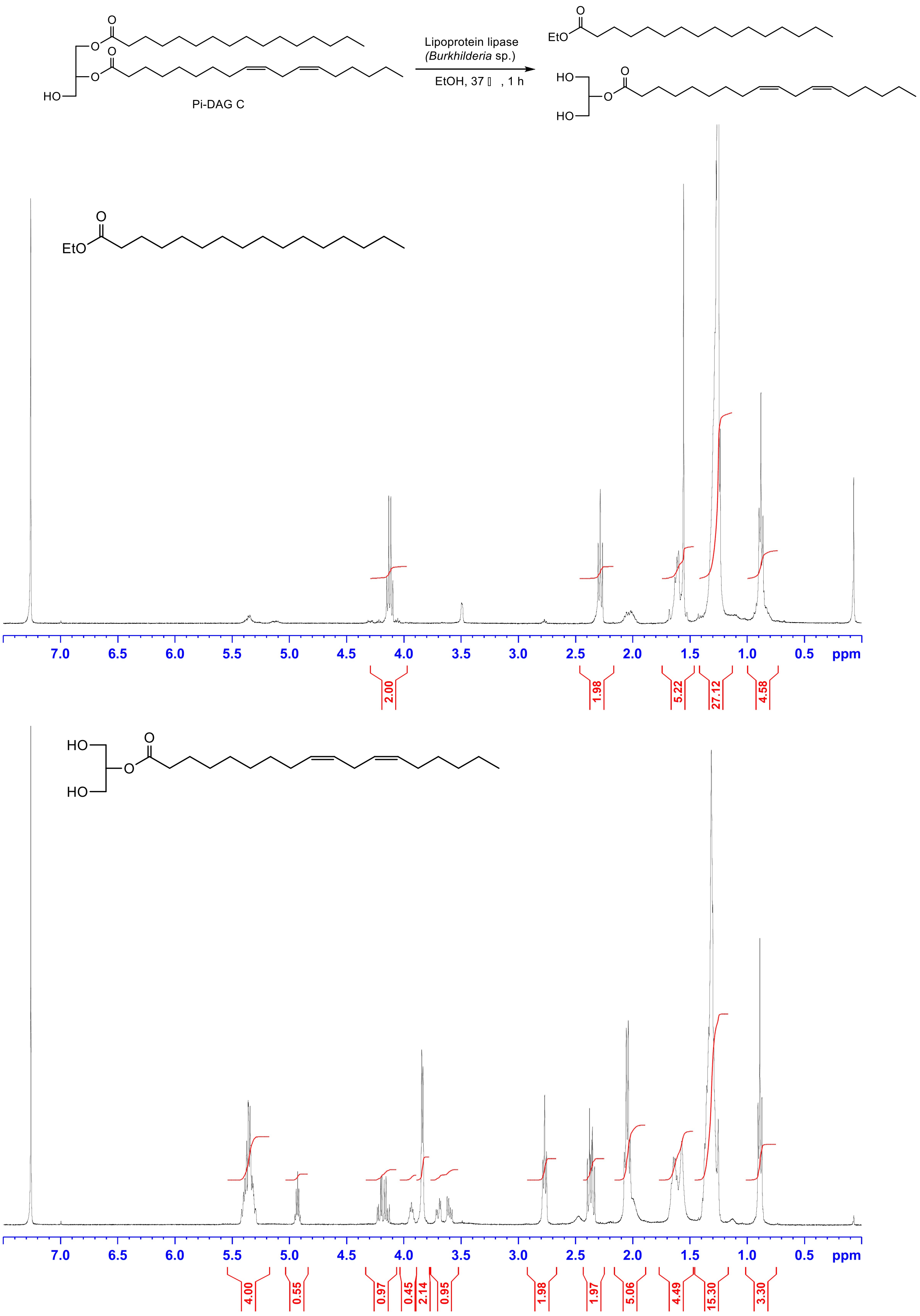


**Figure S26.** (continued).
**(C)** Confirmation of fatty acid linkage in Pi-DAG C.
Pi-DAG C was treated with a lipase to give ethyl palmitate and 2-O-acylglycerol, which were confirmed by $^{1}$H NMR ($CDCl_3$, 400 MHz).

### Pi-DAG D

A

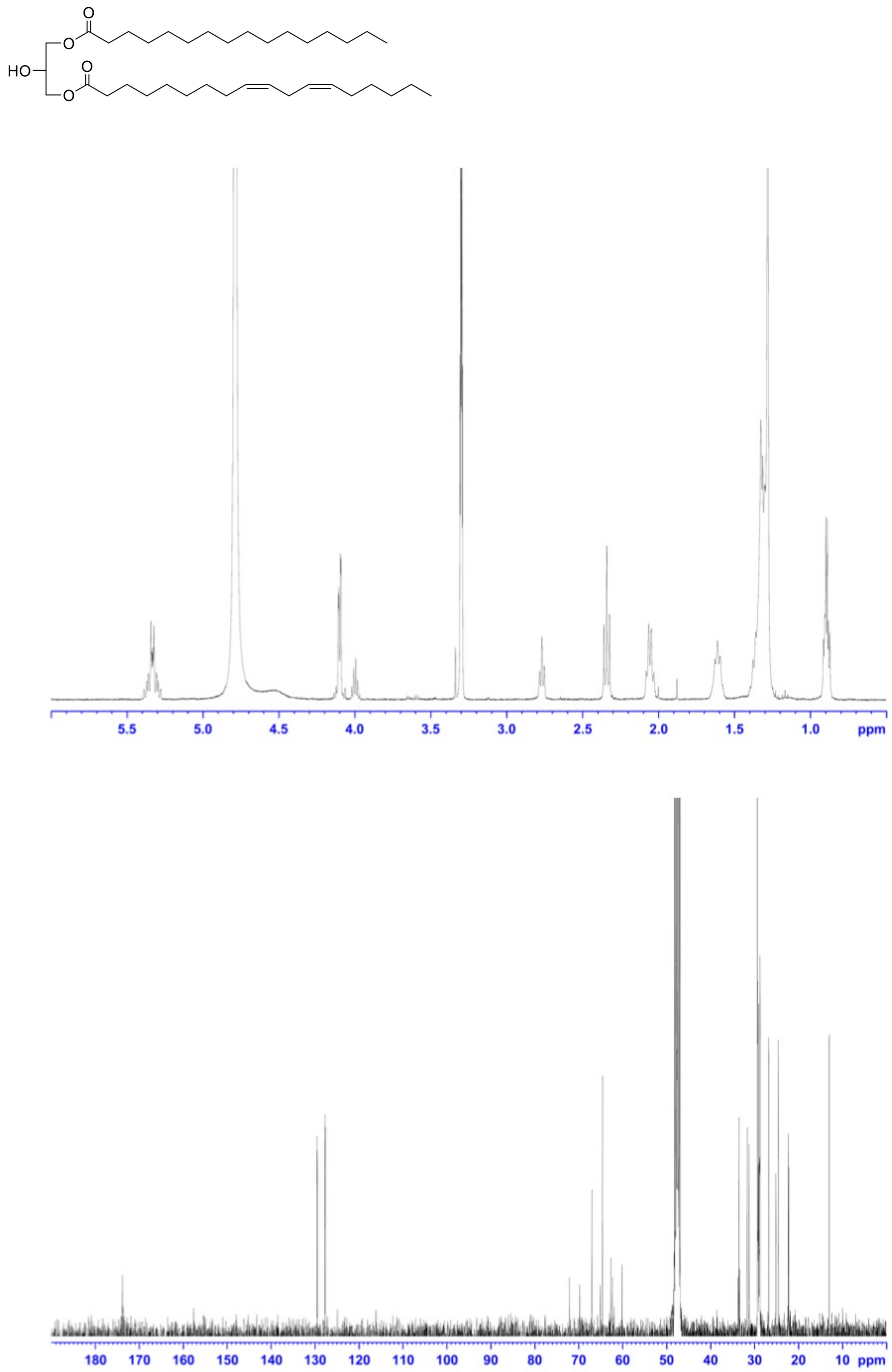


**Supplemental Figure S27. (A)** NMR spectrum of Pi-DAG D (MeOD, 400 MHz for $^{1}$H, 100 MHz for $^{13}$C). The $^{1}$H NMR data (top) indicates that the two fatty acyl moieties are same as those of Pi-DAG C. The chemical shifts at 4.11 ppm (4H) for H-1 and H-3 indicate that both the primary alcohols of glycerol are acylated.

**B**

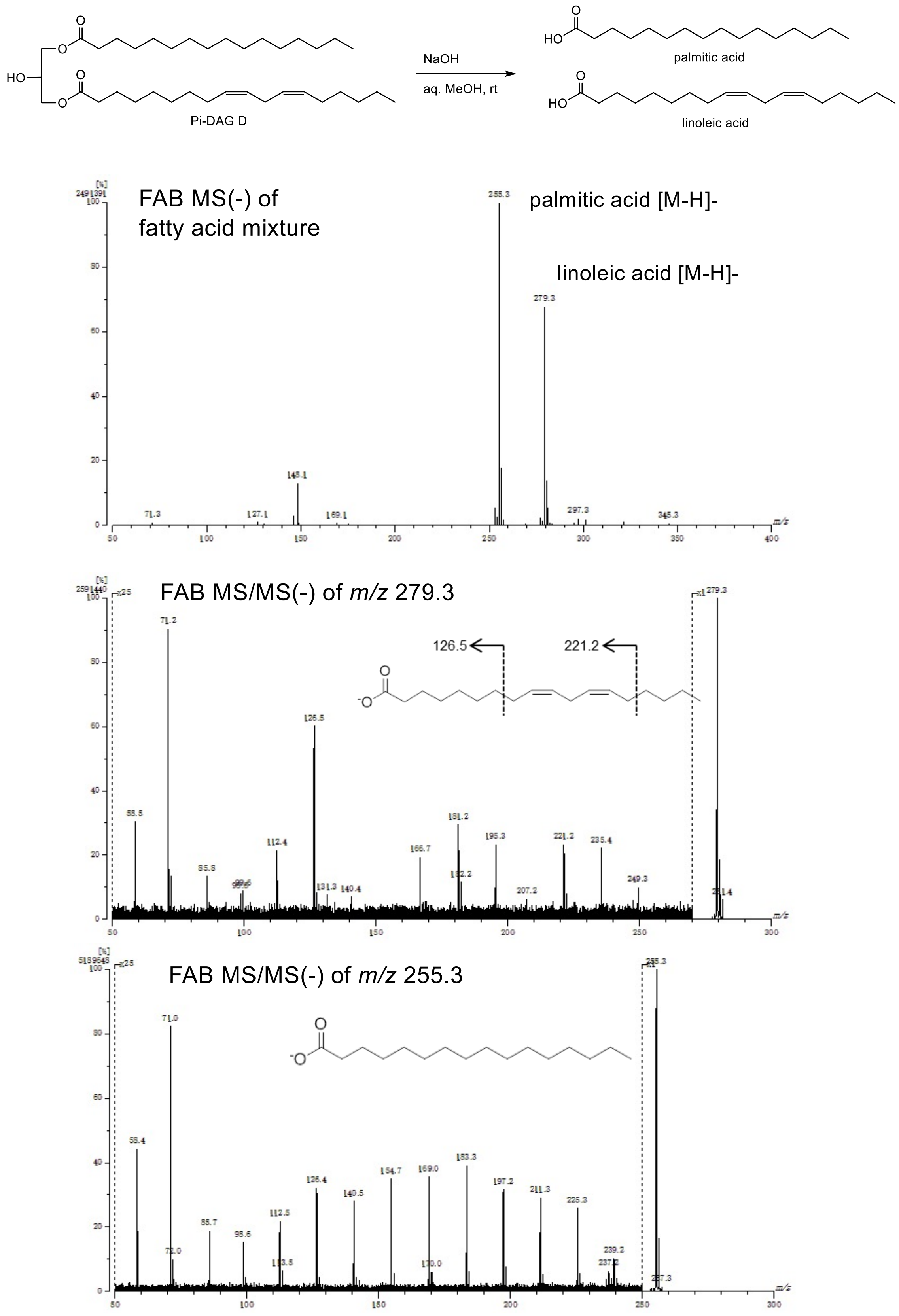


**Figure S27.** (continued).

**(B)** Determination of fatty acids in Pi-DAG D by negative ion FAB MS

Hydrolysis of Pi-DAG D gave two fatty acids as shown in the 1st MS. Both acids were characterized by FAB MS (top) and FAB MS/MS (middle and bottom).

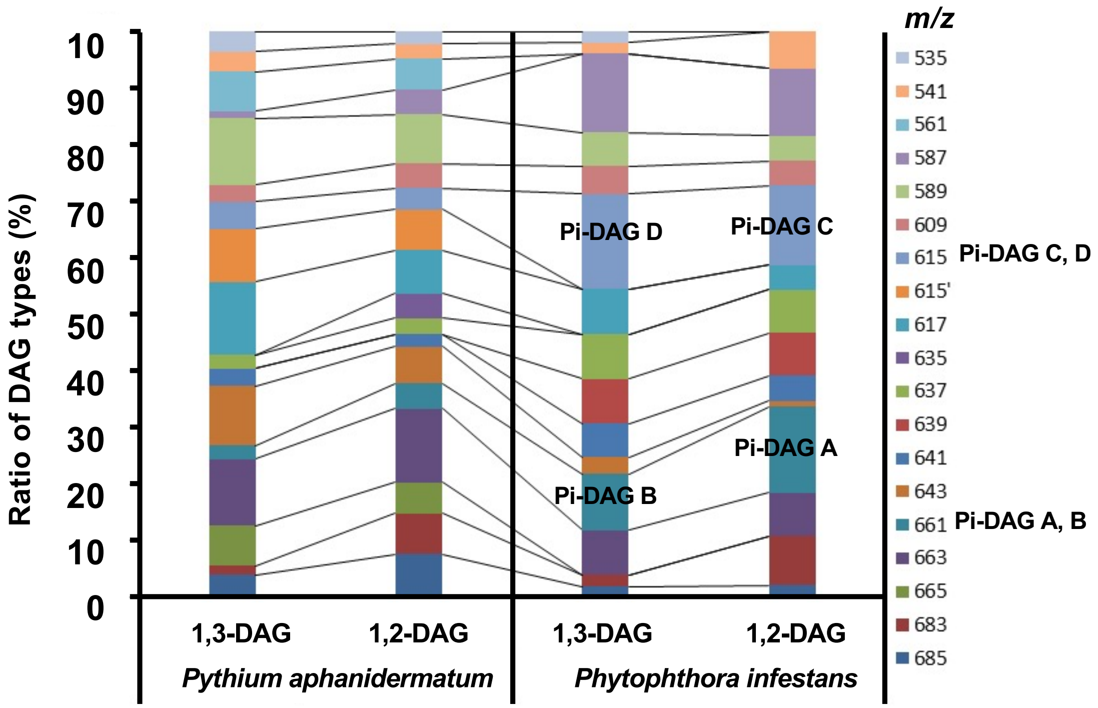


**Supplemental Figure S28.** Pi-DAG contents in mycelia of oomycete plant pathogens. 1,2-DAGs and 1,3-DAGs were partially purified from mycelia of oomycete pathogens *Phytophthora infestans* and *Pythium aphanidermatum*, and Pi-DAGs contents in these pathogens were analyzed by LC/MS. Note that Pi-DAG contents were not quantified but indicated by relative peak area.

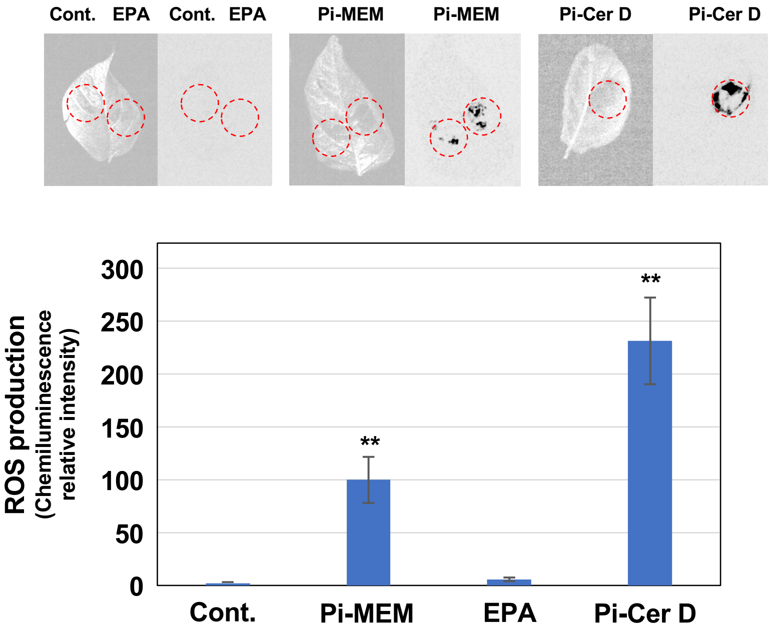


**Supplemental Figure S29.** Eicosapentaenoic acid (EPA) cannot induce the production of reactive oxygen species (ROS) in potato leaves. Potato leaves were treated with 1% DMSO (Cont.), 100 µg/ml methanol extract of *P. infestans* mycelia (Pi-MEM), EPA or Pi-Cer D and production of ROS was detected as L-012-mediated chemiluminescence 12 hrs after treatment. Data are means ± SE (n = 8). Scores are chemiluminescence intensities relative to that of Pi-MEM-treated leaves. Data marked with asterisks are significantly different from control as assessed by two-tailed Student's *t* tests: **$P < 0.01$.

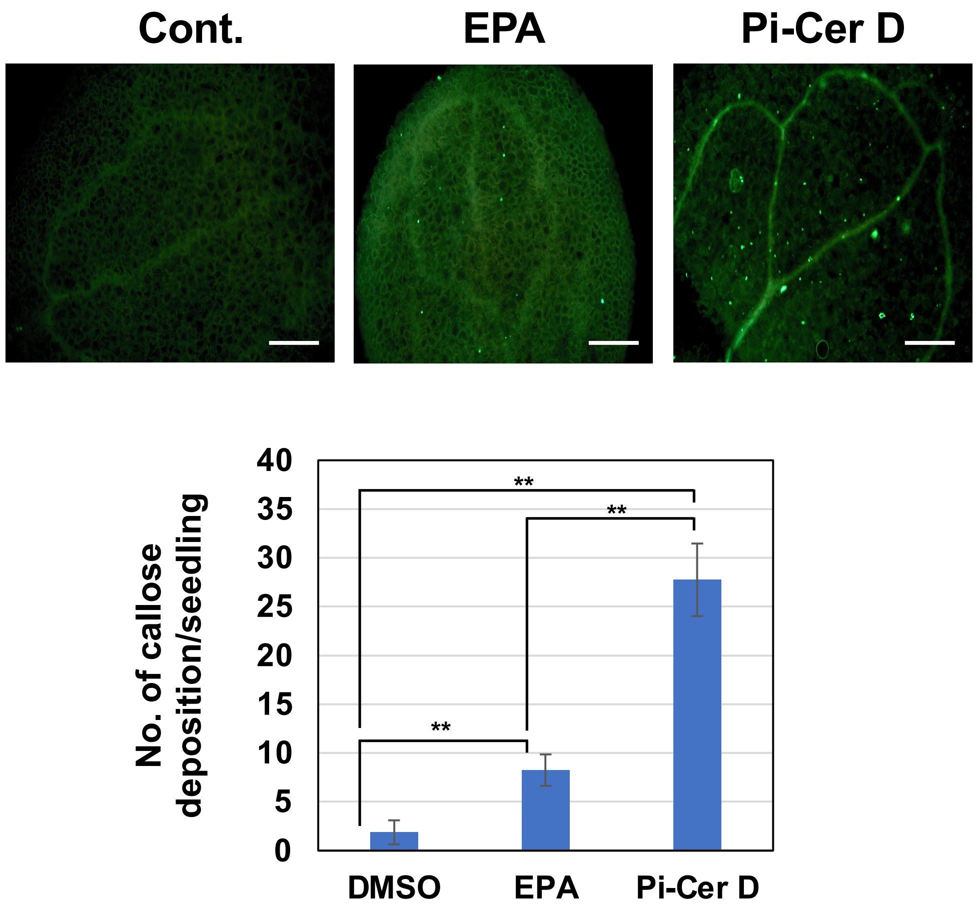


**Supplemental Figure S30.** Callose deposition of Arabidopsis seedlings treated with eicosapentaenoic acid (EPA) or Pi-Cer D. Arabidopsis seedlings (10-days old) were treated with 1% DMSO (Cont.), 100 µg/ml EPA or Pi-Cer D and strained with aniline blue staining at 24 h after treatments. Callose deposition (fluorescence spots) was counted for each treatment (n = 8) by fluorescence microscopy. Bars = 100 µm. Data marked with asterisks are significantly different from control as assessed by two-tailed Student's *t* tests: **$P < 0.01$.

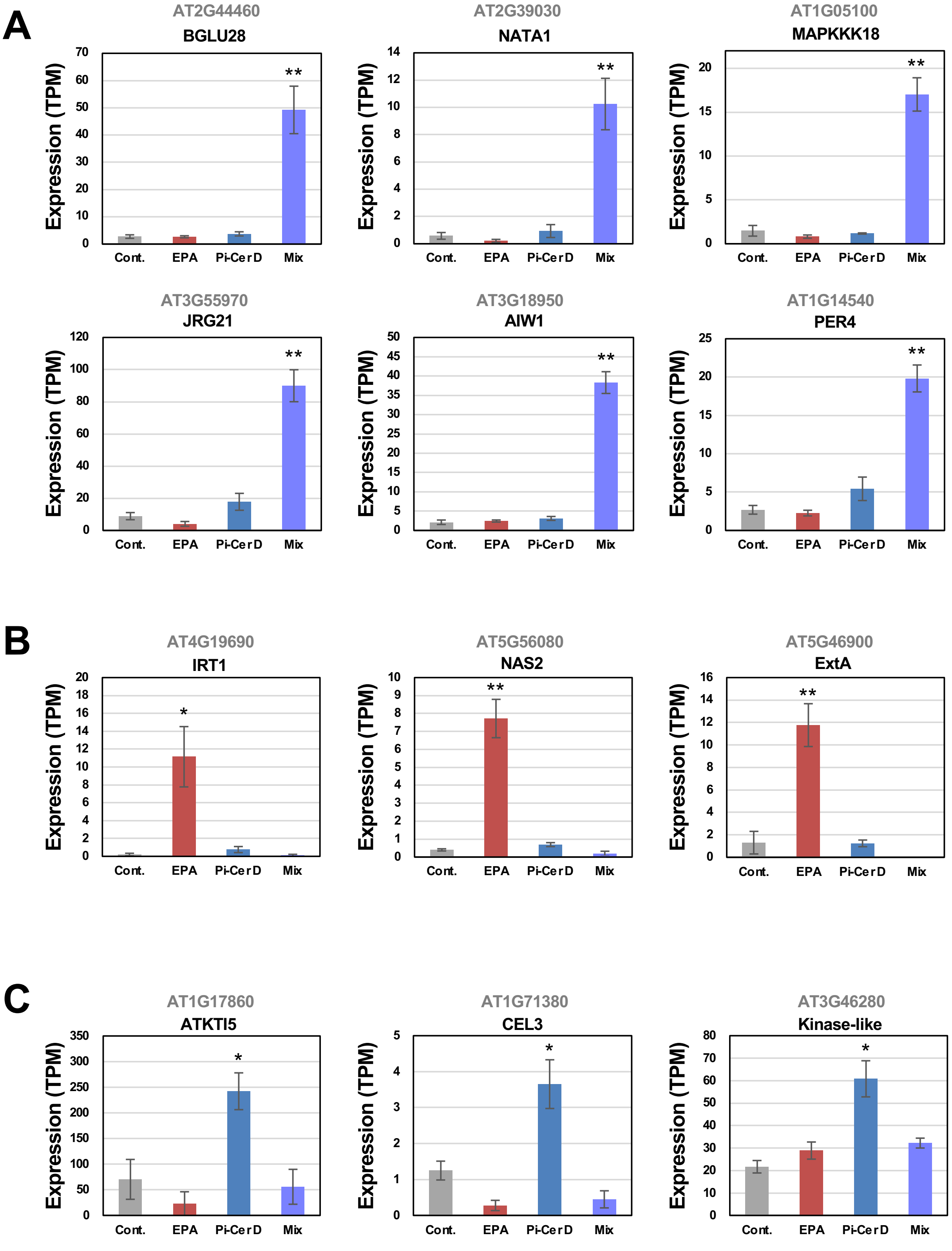


**Supplemental Figure S31.** Expression profiles of Arabidopsis genes upregulated by co-treatment of eicosapentaenoic acid (EPA) and Pi-Cer D (A), EPA (B) or Pi-Cer D (C). Gene expression (TPM value) was determined by RNA-seq analysis of Arabidopsis seedlings treated with 1% DMSO (Cont.), 100 μg/ml EPA, 100 μg/ml Pi-Cer D, or the mixture of 100 μg/ml EPA and 100 μg/ml Pi-Cer D for 12 h. Data are means ±SE (n = 3). Data marked with asterisks are significantly different from control as assessed by the two-tailed Student's *t* test: **p<0.01, *p<0.05.

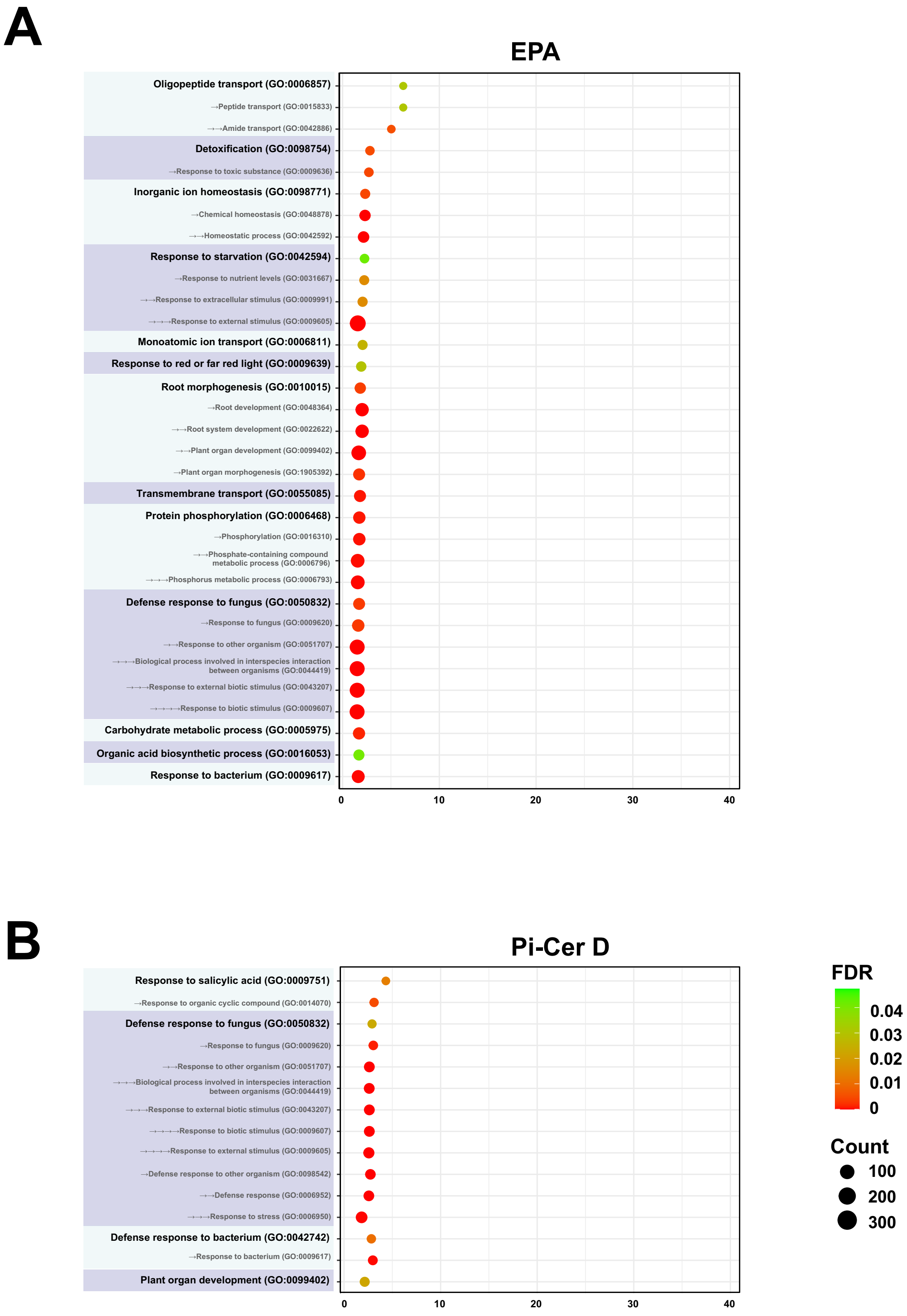


**Supplemental Figure S32.** Gene ontology (GO) enrichment analysis of up-regulated genes in Arabidopsis treated with 100 µg/ml EPA (A), 100 µg/ml Pi-Cer D (B) or their mixcure (C, 100 µg/ml EPA and 100 µg/ml Pi-Cer D) for 12 h. GO term enrichment is expressed as significantly different fold enrichment of mapped genes (FDR p<0.05). The dot size (and numbers beside the dots) indicates the number of significantly up-regulated genes (see Figure 8B) associated with the process and the dot color indicates the significance of the enrichment.

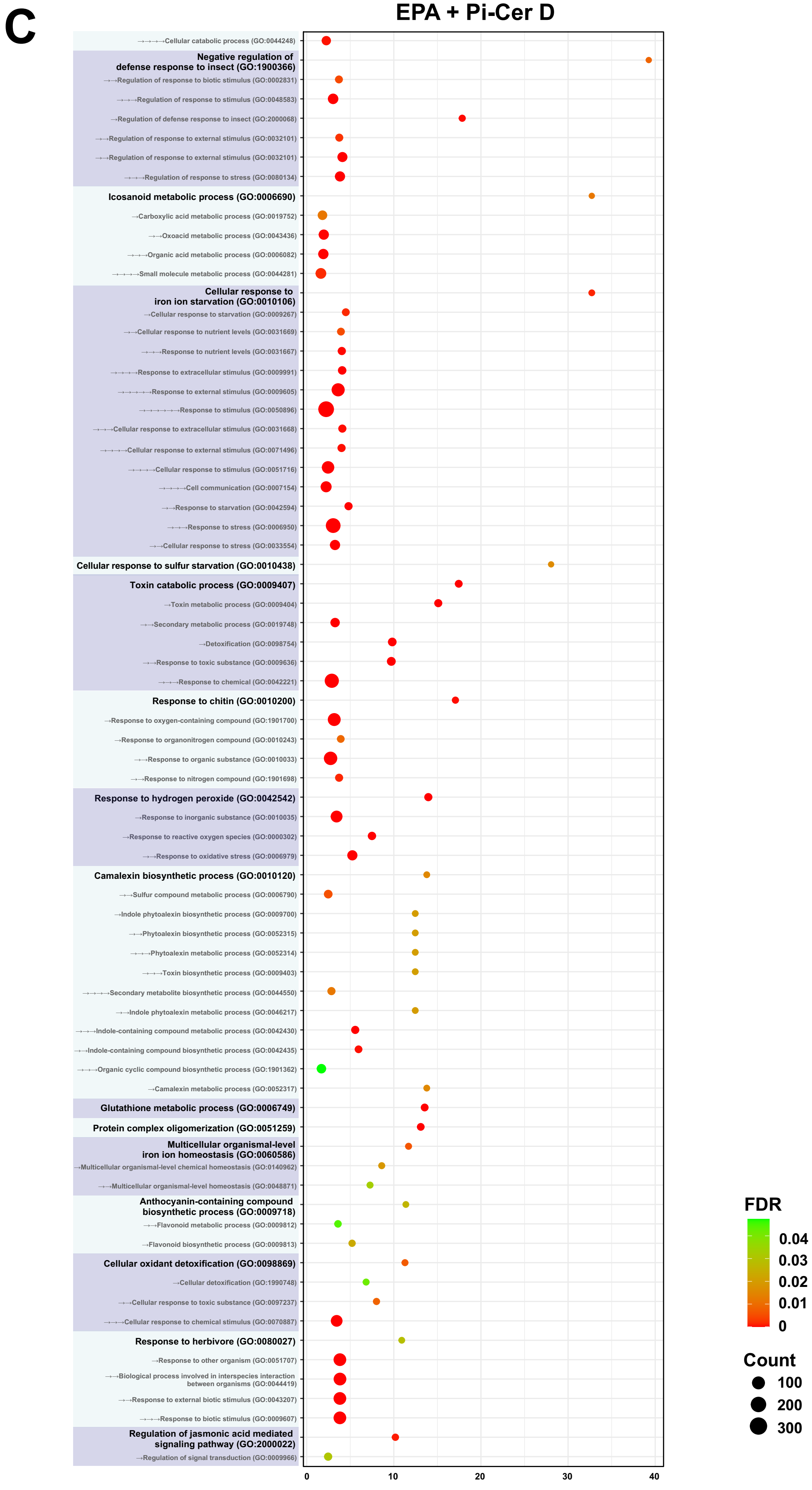


**Supplemental Figure S32.** (continued)

# C

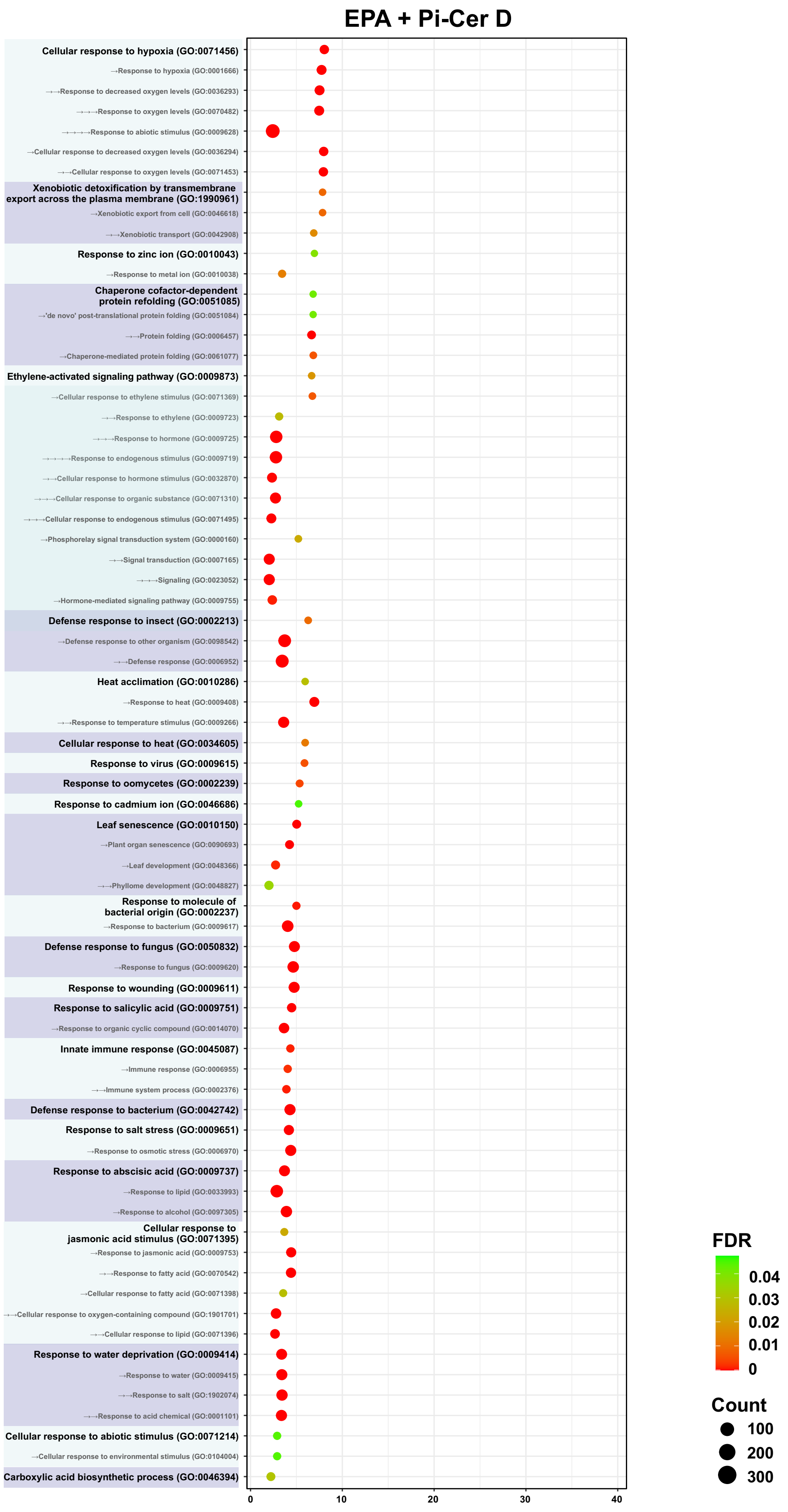


**Supplemental Figure S32.** (continued)
